# Exposure to false cardiac feedback alters pain perception and anticipatory cardiac frequency

**DOI:** 10.1101/2023.06.07.544025

**Authors:** Eleonora Parrotta, Patric Bach, Giovanni Pezzulo, Andrea Zaccaro, Mauro Gianni Perrucci, Marcello Costantini, Francesca Ferri

## Abstract

The experience of pain, like other interoceptive processes, has recently been conceptualized in terms of predictive coding and free energy frameworks. In these views, the brain integrates sensory, proprioceptive, and interoceptive signals to generate probabilistic inferences about upcoming events, which shape both the state and the perception of our inner body. Here we ask whether it is possible to induce pain expectations by providing false faster (vs. slower) acoustic cardiac feedback before administering electrical cutaneous shocks. We test whether these expectations will shape both the perception of pain and the body’s physiological state toward prior predictions. Results confirmed that faster cardiac feedback elicited pain expectations that affected both perceptual pain judgments and the body’s physiological response. Perceptual pain judgments were biased towards the expected level of pain, such that participants illusorily perceived identical noxious stimuli as more intense and unpleasant. Physiological changes mirrored the predicted level of pain, such that participants’ actual cardiac response in anticipation of pain stimuli showed a deceleration in heart rate, in line with the well-known orienting cardiac response in anticipation of threatening stimuli (Experiment 1). In a control experiment, such perceptual and cardiac modulations were dramatically reduced when the feedback reproduced an exteroceptive, instead of interoceptive, cardiac feedback (Experiment 2). These findings show that cardiac perception can be understood as interoceptive inference that modulates both our perception and the physiological state of the body, thereby actively generating the interoceptive and autonomic consequences that have been predicted.

## Introduction

Far more complex than the mere transmission of nociceptive inputs, pain is a component of the interoceptive system (Craig, 2003b). It is defined as an unpleasant sensory and emotional experience associated with, or resembling, actual or potential tissue damage (International Association for the Study of Pain, IASP, 2020). It can occur in the absence of physical harm (Loeser & Treede, 2008) and is permeated by all the available sources of information, including sensory, emotional, cognitive, and social components as well as prior information and expectations (Atlas & Wager, 2012; Tracey, 2010; Williams & Craig, 2016). While the last decades provided important advances in the understanding of pain, developing an overarching theory that accounts for all the dimensions of pain still remains challenging (for a full discussion, see Moayedi and Davis, 2013).

Current views propose predictive processing models as a suitable candidate to account for the multifaceted experience of pain (Büchel et al., 2014; Kiverstein et al., 2022; Song et al., 2019, 2021). Interoceptive inference and the Embodied Predictive Coding frameworks (e.g., Barrett, 2017; Barrett and Simmons, 2015; Pezzulo, 2014; Seth, 2013; Seth and Friston, 2016) conceptualise the brain as an active generator of inferences, which maintains an internal model of both the internal and the external world and attempts to fit it with incoming inputs through a process of Bayesian hypothesis testing and revision. Interoceptive sensations are thought to be derived from the integration of different sources of information into an expectation of the body’s upcoming changes, which are kept in check by the actual state of the body (Barrett & Simmons, 2015). However, people’s predictions do not always mirror reality, but can be inaccurate, leading to a gap between the expected and the actual present state. The goal of inferential processes is to reduce this difference between one’s internal models and the actual interoceptive input, thus converging towards the *brain’s best guess* about the body’s state (Ainley et al., 2016; Seth et al., 2012; Seth & Friston, 2016).

An important feature of these accounts is that the difference between one’s internal models and the sensory data can be minimized not only by updating internal models to better fit the data (i.e., perceptual inference), but also by performing actions, changing bodily states themselves (i.e., active inference, see Friston, 2010, 2005; Parr et al., 2022). For example, an expectation of a threatening, painful stimulus could be realised both by making the painful stimulus *appear* more painful than it really is, thereby fitting one’s internal model to bodily reality, or by *actually* modulating the state of the body and shape the physiologic response to pain towards previous predictions, for example by engaging autonomic reflexes (e.g., typical decrease in the heart rate in relation to pain anticipation, e.g., Bradley et al., 2008, 2005; Colloca et al., 2006; Lykken et al., 1972; Taggart et al., 1976; Tracy et al., 2017).

There is suggestive evidence for both components in the extant literature on pain. The assumption that prior information modulates pain perception underlines decades of work on placebo and nocebo effects (i.e., the expectation-effect; Crombez et al., 1998; Keltner et al., 2006; Petrovic & Ingvar, 2002; Price et al., 1999; Wager, 2005; Wager et al., 2004). On a neuronal level, expecting pain alters the neural mechanisms in pain-processing regions, thought to reflect a combination of nociceptive inputs and top-down information (Atlas et al., 2010; Keltner et al., 2006; Koyama et al., 2005; Wiech et al., 2008), even at a very early stage of processing (Eippert et al., 2009; Geuter & Büchel, 2013). Moreover, such changes can persist - or even grow over time - in the absence of disconfirming evidence (Atlas et al., 2010; Colloca et al., 2010; Craggs et al., 2008; Jepma & Wager, 2015; Koban & Wager, 2016; Montgomery & Kirsch, 1997; Vase et al., 2005, 2011), turning pain experience into potentially a self-fulfilling prophecy (Jepma et al., 2018).

As for the assumption that unfulfilled predictions can be resolved by engaging autonomic reactions, there is evidence that prior pain expectations, and pain itself, modulate physiological responses. Expecting pain induces augmented activity in the sympathetic system (e.g. increases in blood pressure and skin conductance) to prepare potential avoidance responses (Barlow et al., 1996; Oka et al., 2007; Tousignant-Laflamme & Marchand, 2006; Yang et al., 2003), as well as a characteristic decrease in heart rate to promote orienting responses, attention and sensory processing (Bradley et al., 2008, 2005; Colloca et al., 2006; Lykken et al., 1972; Taggart et al., 1976; Tracy et al., 2017; for a full discussion, see Skora et al., 2022). Once pain is actually experienced, however, heart rate rises, reflecting robust sympathetic activation and a well- documented autonomic response to nociceptive input (Tousignant-Laflamme et al., 2005; Terkelsen et al., 2005; for a review, see Forte et al., 2022).

Together, these findings support the idea that the experience of pain may rely on top-down predictive mechanisms, which shape not only the subjective perception of pain, but also its neural and bodily correlates, both when pain is experienced and merely expected. However, to our knowledge, no study exists that tracks both the perceptual and physiological responses to mismatches between pain expectation and bodily reality. Moreover, in previous work, pain expectations were typically induced by exteroceptive cues (e.g., visual, Wiech et al., 2014; Jepma et al., 2019; auditory, Colloca et al., 2006; Atlas et al., 2010). Although interoceptive processes may have contributed to the observed effects, these studies did not specifically target interoceptive sources of information within the inferential process.

If one assumes that pain and bodily states are integrated within a unified interoceptive system, then information about the internal condition of the body should be tightly linked to the experience of pain. Among such interoceptive streams, cardiac activity appears to be closely intertwined with the experience of pain, likely due to a close integration between the pain system and the neural circuits involved in cardiovascular regulation (Craig, 2003b, 2003a, 2008). For example, pain-related evoked potentials and nociception are modulated across the cardiac cycle (Edwards et al., 2001, 2002, 2008; Martins et al., 2009; McIntyre et al., 2006), and both the anticipation of and the response to pain are associated with changes in heart rates (Colloca et al., 2006; Mischkowski et al., 2018; for a review, see Kyle and McNeil, 2014). Moreover, when estimating their own cardiac frequency, people’s estimates integrate the level of threat (i.e., pain) associated with upcoming events (Parrotta et al., 2024), reflecting the (fictitious) belief that anticipating threat increases heart rates. Even though anticipating pain typically decreases real heart rates (the well-known orienting response, Sokolov et al., 2002), people therefore illusorily perceive a higher cardiac frequency when expecting painful stimulation (Parrotta et al., 2024).

The tight coupling between heart rate and pain experience suggests that it should be possible to modulate the experience of pain - and the anticipatory bodily response - by manipulating *interoceptive cardiac* feedback, that is, by misleading people into believing that their heart rate is increasing in anticipation of a noxious stimulus. Under predictive architectures, persuading one’s internal model towards the fictitious evidence of increased heart rates should then induce changes both in *pain perception* (i.e., perceptual inference) and in the *actual state of the body* (i.e., active inference). Specifically, the former process would correspond to an interoceptive illusion of increased pain, whereas the latter would correspond to physiological and autonomic adaptations to the predicted level of pain. We hypothesized that a cardiac feedback manipulation that renders interoceptive streams more similar to those expected during pain would produce an interoceptive illusion of pain as well as ensuing autonomic adjustments - as hypothesized by interoceptive inference and Embodied Predictive Coding theories. To test this idea, Experiment 1 provided participants with auditory cardiac feedback, via headphones, that could diverge from their actual heart rate by being either faster (as when experiencing pain or threat) or slower (as in relaxed states), before the administration of a noxious electrical stimulation, while their ECG was recorded. Once the pain stimulus was administered, subjects were asked to rate its intensity and unpleasantness.

We predicted, first, that the exposure to faster (vs. slower) cardiac feedback would induce expectations of increased levels of pain, such that noxious stimuli are felt and reported as more unpleasant and intense. Accelerated heart rate is a salient, evolutionarily conserved signal of arousal, threat, and nociception, a mapping continually reinforced physiologically when responding to *actual* pain (e.g. for a review, see Forte et al., 2022). If these expectations are integrated with interoceptive inference, they should therefore heighten assessment of pain.

Second, expectations of threat induced by the faster feedback should be fulfilled through anticipatory parasympathetic reflexes, such that participants’ real heart rates decrease when exposed to the faster, compared to the slower cardiac feedback, in line with the well-known orienting cardiac response when expecting threat or pain (Bradley et al., 2008, 2005; Colloca et al., 2006; Lykken et al., 1972; Taggart et al., 1976; Tracy et al., 2017; for a full discussion, see Skora et al., 2022). This anticipatory cardiac deceleration reflects an allostatic adjustment implemented by the brain in response to predicted (instead of experienced) threat.

Our hypotheses were theoretically derived from Embodied Predictive Coding accounts of interoception. Our previous findings (Parrotta et al., 2024) suggest that when the brain estimates cardiac activity, it may assign special relevance to pain and anticipated threat, implying that cardiac signals are weighted within the interoceptive schema of pain (Schoeller et al., 2022). In particular, we previously demonstrated that the mere expectation of pain can induce a cardiac interoceptive illusion: participants reported increased heart rate when anticipating pain, even in the absence of any actual physiological change (Parrotta et al., 2024). Here, we tested the counterpart of this illusion: if pain expectations can bias cardiac perception, then manipulating cardiac feedback should, in turn, influence pain perception. Presenting participants with artificially accelerated heartbeat feedback may act as a misleading prior, enhancing perceived pain intensity despite unchanged nociceptive input. Evidence for such a mechanism is further suggested by the evidence that providing mismatched cardiac feedback in contexts where a faster heart rate is normally expected (i.e. sexual arousal, Rupp & Wallen, 2008; Valin, 1996; physical exercise, Iodice et al 2019) can enhance the perception of physiological states associated with those experiences (Crucian et al., 2000; Iodice et al., 2019).

To rule out that potential modulations in participants’ pain perception and cardiac state were merely associated with the frequency of the feedback, rather than a heartbeat, Experiment 2 replicated the design in a second group of participants, with the only difference being that the congruent or incongruent feedback would not be a heartbeat tone, but an unrelated exteroceptive stimulus. We predicted that, as participants would have no expectation that such an external stimulus signals a preparation for a threatening, painful stimulus, variations in the rate of this exteroceptive stimulus should lead to no (or reduced) perceptual and cardiac changes.

If confirmed, the results would provide the first evidence that predictive processes are generated by the simulation of interoceptive streams, influencing not only our perception of pain but also its heart-related physiological response: two changes that jointly minimize the discrepancy between the expected and the actual homeostatic (interoceptive) state. This evidence would corroborate the assumption that predictions originate along the embodied multisensory sphere by integrating multiple sources of information (i.e., interoceptive) within the inference. Ultimately, the results may provide important insights into how the experience of pain is actively built through predictive processes.

### Experiment 1

Experiment 1 tests whether false cardiac feedback of an accelerated (i.e., faster), relative to a decelerated (i.e., slower) heart rate, alters both pain perception and the cardiac anticipatory response to noxious stimuli.

## Methods

### Participants

Thirty-five participants (mean age 25.11, *SD*= 2.94, 21 women) took part in the experiment, recruited from Gabriele D’Annunzio University and the wider community. All were right-handed with normal or corrected-to-normal vision. Exclusion criteria for taking part were self-reported chronic and acute pain, neurological disease, serious cardiovascular disease (i.e., any type of disease involving the heart or blood vessels that might result in life-threatening medical emergencies, e.g., arrhythmias, infarction, stroke), or conditions that could potentially interfere with pain sensitivity (e.g. drug intake or skin diseases). Participants were instructed not to drink coffee or smoke cigarettes. After taking part in the experiment, participants were excluded if the correlation between the desired five levels of the nociceptive stimulus intensity and their ratings in the no-feedback condition was below *r*=0.75, suggesting insensitivity to the varying nociceptive degrees of the electrical stimulus. No participants were excluded based on this criterion. All gave written informed consent, were unaware of the purposes of the study, and were fully debriefed about it at the end of the experiment. Ethical approval from the local ethics board was obtained. One participant was excluded as, at the end of the experiment, they declared to suffer from a cardiac defect that made them realize the manipulation of the cardiac feedback.

A sensitivity analysis with G*Power 3.1 (Faul et al., 2007) showed that the final sample size of 34 provides .90 power to detect effects with Cohen’s *d* = 0.57 (SESOI of δ = 0.34). Following recommendations, we do not estimate power based on previously reported effect sizes as this neglects uncertainty around reported effect size measurements, especially for new effects for which no reliable effect sizes can be estimated across studies (Albers & Lakens, 2018; Anderson et al., 2017). Instead, we report sensitivity analyses that reveal the effect sizes that the crucial comparison between faster and slower feedback is well-powered to detect, given our experimental parameters (i.e., target power, sample size and type of test), as well as the smallest effect size of interest it can, in principle, reveal (SESOI, Lakens, 2022). Note that these values are well below the observed top-down effects on pain ratings of *d* = .7 reported before (e.g., Iodice et al., 2019).

### Apparatus

Painful stimuli were electrical pulses delivered using a constant-current electrical stimulator (Digitimer DS7A) controlling a pair of neurological electrodes attached to the phalanx of the middle finger of the participant’s left hand, which provide a precise constant current, isolated stimulus, controllable in pulse duration and amplitude. The duration of the electrical stimuli (2 ms) was kept fixed over the experiment and it was established during a calibration phase before the experiment.

Cardiac recording was performed by a Biopac MP 160 system (Biopac Systems Inc., USA). ECG was recorded continuously from two electrodes attached to the lower ribs and one over the right mid-clavicle bone (reference electrode). The ECG signal was sampled at 2 kHz with the Biopac Acqknowledge 3.7.1 software (Biopac Systems Inc., USA) according to the manufacturer’s guidelines. The ECG signal was then fully analysed in MATLAB (R2020a). The tone used for the creation of the feedback was the sound of a single heartbeat, gathered from https://freesound.org and manipulated in Audacity. The feedback audio was then created in MATLAB (R2020a), repeating the single heartbeat sound according to the desired frequency (see *Procedure*).

Stimulus presentation was controlled through E-Prime (Psychology Software Tools Inc., Pittsburgh, USA), which interfaced with the pain stimulators and the Biopac system via a parallel port.

### Procedure

Upon arrival at the lab, participants were briefed by the experimenter. After providing consent, they were placed in a comfortable chair with their left hand placed on a table, and the ECG electrodes were applied after cleaning the skin. To increase the ambiguity and thus predictive influences acting on the pain stimulus (Yon & Frith, 2021), the intensity of the noxious input varied between five intensities, which were identified for each participant in an initial calibration session. To do so, participants were informed that they would undergo a psychophysical calibration procedure to determine their subjective response to increasing stimulus intensities. The first stimulus was delivered at a low intensity, which is below the threshold for pain perception in most people. The intensity increased in a ramping procedure up to a maximum of five volts. Participants verbally rated the pain intensity for each stimulus on a 0-100 Numerical Pain Scale (NPS).

Following previous research (Atlas et al., 2014; Colloca et al., 2006; Hird et al., 2019) a pain intensity rating of NPS 20 denoted “just painful”, NPS 50 denoted “medium pain”, and NPS 80 marked the point at which the stimulus was “just tolerable”. We identified a ‘low’ pain level of 10, a ‘low-medium’ pain level of 30, a ‘medium’ pain level of 50, a ‘medium-high’ pain level of 70, and a ‘high’ pain level of 90. Level 100 was considered the point where the participant did not wish to experience a higher stimulation level in the experimental session and was not used. We repeated this procedure three times and computed the average stimulus intensities over these three repetitions corresponding to NPSs 10, 30, 50, 70, and 90.

Participants then underwent a pre-experiment test procedure: stimulus intensities corresponding to their pain intensity ratings NPS 10 to 90 were delivered in a pseudo-randomised order four times and participants were instructed to identify the intensity of each pulse. Participants had to correctly identify 75% of stimulus intensities to proceed to the main experiment. If they did not achieve this in the test procedure, the intensities were adjusted, and the test was repeated until participants correctly identified 75% of stimulus intensities (Hird et al., 2019). Participants were excluded if the correlation between their ratings of the five levels of the nociceptive stimulus intensities and the actual stimulation intensities within the baseline phase was below *r*=0.75, suggesting insensitivity to the varying nociceptive degrees of the electrical stimulus, and an essentially flat response profile. No participants were excluded with this criterion. This procedure was needed because, although the paradigm introduced ambiguity by using different levels of actual stimulus intensity, it was important to ensure that participants developed a stable internal representation of the pain scale and to ensure that illusion effects were not due to an inability to reliably distinguish between intensities. Note that this correlation criterion only requires that all participants’ ratings generally increase with actual stimulus intensities; participants are still free to vary in absolute values and the range they use on the pain scale. Supplementary Figure 11 shows the stimulation intensities corresponding to each of the five pain levels obtained through this calibration.

Having established the individual levels of intensity for each stimulation, participants were informed that their task was to rate the intensity and unpleasantness of nociceptive stimulations. We carefully described the distinction between intensity and unpleasantness ratings using the standard language developed by Price et al. (1989), emphasizing that pain intensity and unpleasantness should be rated independently.

The experiment started with a no-feedback session in which participants’ heart rate in anticipation of the shock (measured with ECG) and perceived level of pain (measured with Numeric Pain Scale and Likert scale) were assessed. These measures were used to normalize our dependent variables for our analysis (see *Data analysis*).

Participants were seated in front of the computer, with the ECG electrodes attached, their left hand placed on the table with the electrodes of the cutaneous electrical stimulation fixed on the phalanx of the middle finger, and their right hand placed on the mouse, ready to start the task. The ECG was recorded for the entire session.

Each trial of the no-feedback phase started with the presentation of a fixation cross appearing on the screen. After 60 seconds, a single electrical shock was administered, randomly varying the level of intensity in each trial - as established in the previous calibration phase (i.e., NPS 10, 30, 50, 70, and 90). After each stimulation, participants first rated the intensity of the painful stimulus (instruction: “How intense was the painful stimulation?”) on a continuous Numerical Pain Scale (NPS) appearing on the screen, from 0 (“not at all painful”) to 100 (“extremely painful”). They then rated the unpleasantness of the painful stimulus (instruction: “How unpleasant was the painful stimulation?”) on a 5-point Likert scale (1=no pain, 2=weak, 3=moderate, 4=severe, 5=extremely severe). The two scales were presented on the screen in succession, and participants made their judgments through a mouse click on the scales. Between trials, a pause screen was shown for 45 seconds, in order to avoid the subsequent trial to be contaminated by the heart rate response to the shock. Then, a new screen appeared with the instructions to press the spacebar to start the new trial.

The no-feedback phase comprised 10 trials, after which participants were asked to wait and rest for 15 minutes before participating in the second experimental feedback phase. This time was used by the experimenter to prepare the acoustic stimuli for use in the second session for each participant. For this purpose, we assessed the individual mean heart rate during the anticipation of the painful stimulus by measuring the mean R-R intervals of the ECG for the whole 1-minute interval preceding the shock, and by transforming them into frequency (1/R-R, Colloca et al., 2006). The congruent feedback consisted of tones of heartbeats reproduced at the same individual HR frequency recorded in the no feedback phase. The incongruent faster and slower feedback consisted of tones of heartbeats reproduced at a frequency corresponding to an R-R interval obtained by either reducing or increasing the length of the original mean R-R interval over the no-feedback phase of 25%, respectively. The choice of a ±25% variation in heart rate feedback was guided by prior research using false cardiac feedback paradigms. Studies in this field have used a range of modulation strategies, from fixed-frequency symbolic feedback (e.g., Valins, 1966; Tajadura-Jiménez et al., 2008) to real-time individualised adjustments without specific percentage values (e.g., Iodice et al., 2019). More recent work has adopted proportional manipulations of instantaneous heart rate, using variations such as -20% (Azevedo et al., 2017), ±30% (Dey et al., 2018), and ±50% (Gray et al., 2007). Our choice of ±25% falls within this established range, aiming to strike a balance between producing a salient modulation and preserving the plausibility of the feedback as a reflection of one’s actual physiology. Specifically, the generation of feedback was tailored to each participant. The feedback was set to be 25% above or below their baseline heart rate, with the feedback gradually increasing or decreasing. This individualized approach ensured that each participant experienced feedback relative to their own baseline heart rate. Moreover, to make the incongruent feedback manipulation credible, the incongruent acoustic feedback always started at a frequency rate that reproduced the individual’s heart rate recorded over the no feedback phase, to then gradually increase or decrease the R-R length according to whether the feedback was slower or faster, respectively. In the feedback phase, participants were informed that their task was similar to the previous no-feedback procedure, namely, to rate the intensity and unpleasantness of pain stimuli. They were also informed that in the 60 seconds preceding the administration of the shock, they would hear acoustic feedback, which was equivalent to their ongoing heart rate. This phase consisted of 18 trials (6 trials per experimental condition) of 60 seconds (i.e., fixation) each, after which the electrical shock was administered, randomly varying its intensity in each trial (i.e., NPS 30, 50, and 70), either congruent, slower or faster than the participant’s heart rate recorded in the no-feedback phase.

This relatively low number of repetitions was a deliberate design choice based on both theoretical and practical considerations. As observed in other body illusion paradigms (e.g., Botvinick & Cohen, 1998; Ehrsson, 2007; Pratviel et al., 2022), repeated exposures can reduce illusion strength due to habituation, decreased plausibility, or increased awareness of the manipulation. Moreover, sessions lasted approximately 1.5 to 2 hours and involved cognitively demanding tasks, limiting the feasibility of increasing trial numbers without introducing participant fatigue or disengagement. Finally, as we used explicit pain ratings - measures typically less noisy than implicit physiological indices - fewer trials were sufficient to detect reliable effects (e.g., Corneille et al., 2024).

As for the use of only these three pain intensities in the test phase, the rationale was to focus on a manageable subset that still covered a meaningful portion of the stimulus spectrum. Following the approach of Iodice et al. (2019, PNAS), we deliberately excluded the extreme calibrated levels (NPS 10 and NPS 90) to avoid floor- and ceiling-effects. Moreover, by restricting the range we ensured that every test intensity could be paired with both a “slower” and a “faster” cardiac- feedback condition derived from an adjacent level - pairings that would have been impossible at the extremes where no neighbouring level exists.

Specifically, each of the three levels of intensity of the nociceptive stimulus was associated with one of the three different types of acoustic feedback from the previous no-feedback phase. As in the previous no-feedback phase, participants’ individual mean heart rate before the painful stimulus (i.e., ECG) and the perceived intensity and unpleasantness of pain (i.e., NPS and Likert scale) were assessed. The order of the experimental conditions (i.e., congruent, slower and faster) was randomly generated for each participant by a web-based computer program (www.randomization.com).

### Body Perception Questionnaire

After the experiment, all participants completed the Body Perception Questionnaire (Short form BPQ- SF) (Poli et al., 2021), to investigate whether either perceptual or autonomic modulation as induced by our task would be predictive of self-reported measures of bodily awareness and reactivity. The Body Perception Questionnaire is a 22-item self-administered questionnaire that assesses awareness and reactivity of the autonomic nervous system, that is, the subjective ability to perceive bodily states and bodily reactions to stress. High scores on the BPQ reflect high awareness of internal bodily signals (i.e., high interoceptive sensibility) and high perceived reactivity of the visceral nervous system. Items ask participants to rate, on a 5-point scale (from 1 = never to 5 = always), the frequency with which they feel aware of bodily sensations (e.g., body awareness subscale “My mouth being dry”), experience supradiaphragmatic reactivity (e.g., supradiaphragmatic reactivity subscale “I feel shortness of breath”), and subdiaphragmatic reactivity (e.g., subdiaphragmatic reactivity subscale reactivity subscale “I have indigestion”). In this work, we focus on the body awareness and supradiaphragmatic reactivity subscales, following prior research (Petzschner et al., 2019). Body awareness plays a critical role in how individuals perceive and interpret bodily signals, which in turn affects emotional regulation and self-awareness. Supradiaphragmatic reactivity refers specifically to organs located or occurring above the diaphragm - such as the heart - compared to subdiaphragmatic reactivity subscales further down. Our decision to include these subscales was further supported by recent work from Petzschner et al. (2021), showing that supradiaphragmatic reactivity predicts attentional modulation of the heartbeat-evoked potential (HEP), an electroencephalographic signature of the heartbeating. Thus, this subscale, and the more general body awareness scale, most closely reflect the interplay between cardiac attention, bodily awareness and the development of the interoceptive illusion.

### Data Analysis

Measures of the heart rate recorded with the ECG (beats per minute) in the feedback phase were normalized relative to the baseline (i.e., no-feedback phase) by subtracting and then dividing by the baseline (Lambert et al., 2024; Mirmoosavi et al., 2024; Sundararaj et al., 2025) using the formula: normalized value = (X- bX)/bX, where X represents the mean value of each measure assessed in the experimental feedback phase and bX the mean value of the measure calculated over the no-feedback phase. As in prior research (e.g., Bartolo et al., 2013; Cecchini et al., 2020; Riello et al., 2019), the same normalization was also applied to pain intensity and unpleasantness measures. We did not have specific hypotheses about how the different levels of noxious stimulus intensity affect the perceptual illusion.

For all three dependent variables (heart rate, pain intensity, pain unpleasantness), our main hypothesis is tested by comparing the effects of faster against the effects of slower feedback. Statistically, this comparison collapses our prediction onto a single and most powerful test of our hypothesis for each of the dependent measures, as faster and slower feedback are assumed to emerge from changes in opposite direction, in contrast to separate comparisons against the congruent feedback for which effect sizes will by necessity be on average half of the combined influence of the faster vs slower comparison. Moreover, as the slower and faster feedback manipulation reflect analogous deviations from the real heart rate, this comparison keeps any effect of a perceived absolute deviation (in either direction) from the real heart rate constant. Assume, for example, that in one participant, heart rates generally decrease as soon as a deviation from the real heart rate is noticed (i.e., in either direction). The comparison between faster and slower feedback will still measure whether this decrease is larger for faster than slower feedback, just as in a participant who does not show such a general decrease or one who shows an increase.

The data from each of our dependent measures (heart rate, pain intensity ratings, pain unpleasantness ratings) were analysed similarly. First, to confirm that the relevant measure of interest was affected by the different types of heart rate feedback, the data were entered into a 1-factor repeated measures ANOVA (Greenhouse-Geisser corrected) with the levels faster feedback, congruent feedback, and slower feedback.

Second, once such an influence was confirmed, a planned paired t-test tested our main prediction: that there would be a difference in objective heart rate as well as subjective measures of pain intensity and pain unpleasantness between trials with faster and slower feedback.

Finally, we tested whether any such effect was mostly driven by the faster instead of the slower feedback, using separate t-tests comparing the faster feedback against the congruent feedback and the slower feedback against the congruent feedback.

Please note that, as noted above, this latter comparison is not necessary to test our hypothesis, which only requires a difference between faster and slower feedback. Indeed, many influential studies omitted such a control entirely (Hill et al., 2024; Makkar & Grisham, 2013; Phillips et al., 1999; Valins, 1966), or if included, typically treat it as a baseline control (e.g., Patchitt et al., 2025). Nevertheless, we included the congruent condition here to provide an informative baseline reference, and situating our findings clearly within the broader experimental context, which suggests a specific role of heartrate deviations upwards (i.e. faster) than downwards (slower) (Iodice, 2019). For all analyses, the assumption of normality was systematically assessed using the Kolmogorov-Smirnov (K-S) test.

## Results

### ECG Heart Rate measurement

We first tested whether the heart rate feedback was treated as an interoceptive stimulus and induced a compensatory adjustment of the real heart rate. If so, faster feedback should elicit a comparative slowdown of real heart rates compared to slower feedback (see Figure 2c). An initial ANOVA confirmed that the different types of heart rate feedback (faster, congruent, slower) affected heart rates, *F*(2, 66) = 4.071, *p* = .022, η*p²* = .110. A planned paired sample t-test showed that heart rates decreased during faster cardiac feedback compared to slower heart rate feedback, *t*(33) = 2.074, *p* = .046, *d* = .36 (Figure 2c). When compared with congruent feedback, significant heart rate differences emerged for faster feedback, *t*(33) = 2.228, *p* = .033, *d* = .38, but not for slower feedback, *t*(33) = .92, *p* = .364, *d* = .16.

**Figure 1.**
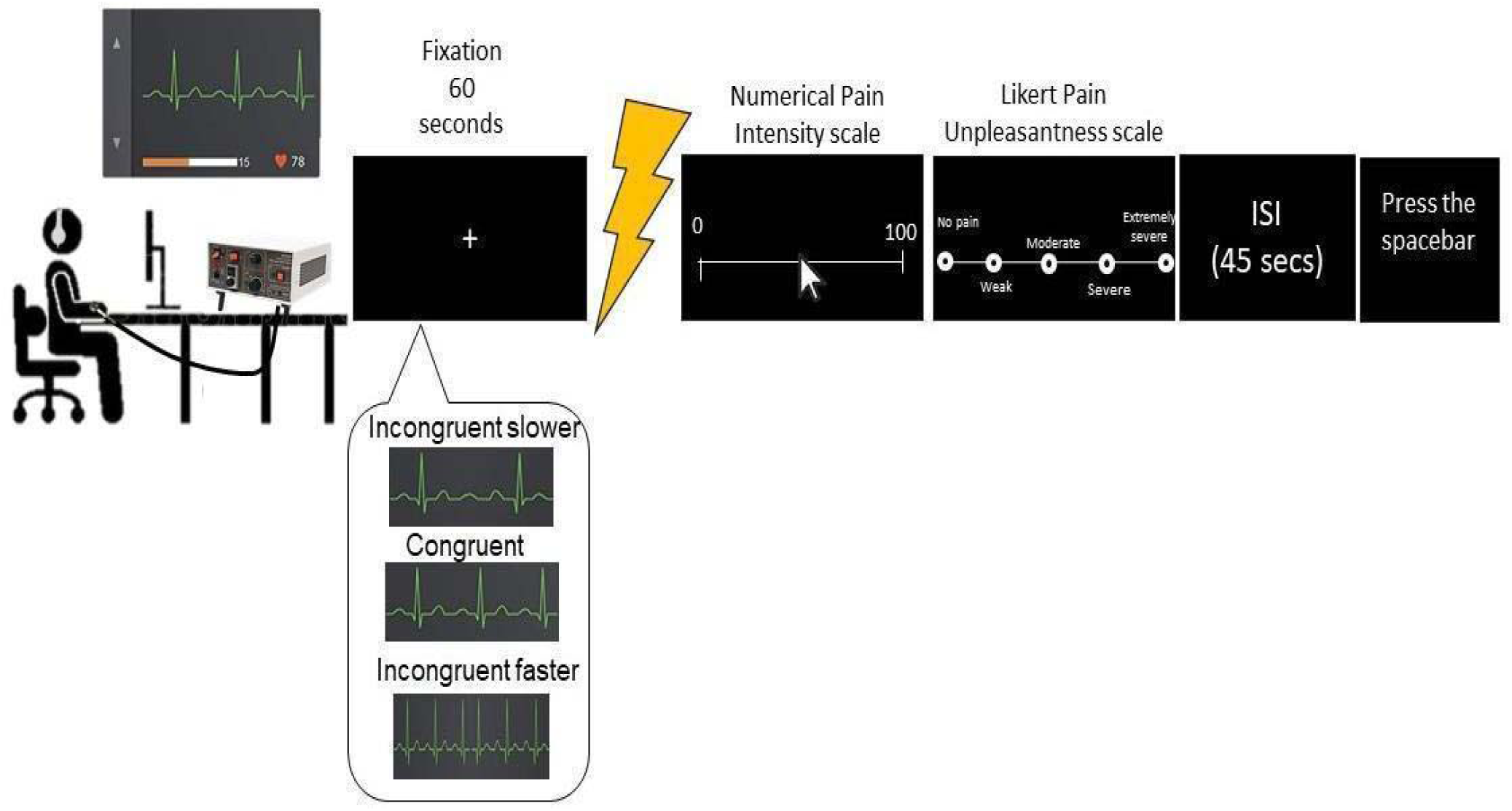
Trial timeline of the experiment. Participants are seated with the electrodes of both the electrical stimulation and the ECG attached. Each trial starts with a fixation cross (60 seconds), after which the noxious stimulation is administered and the pain intensity (i.e., Numerical Rating Scale) and unpleasantness (i.e., Likert scale) ratings are collected. After each trial, a pause screen of 45 seconds was shown (i.e., ISI, Inter Stimulus Interval), after which participants proceeded to the next trial by pressing the spacebar. The trial timeline was identical for both the no-feedback and feedback phases, with the only exception that in the feedback phase participants were provided with the acoustic feedback (i.e., slower, faster, congruent) about their HR, reproduced for the whole fixation period (60 seconds) preceding the shock.

**Figure 2.**
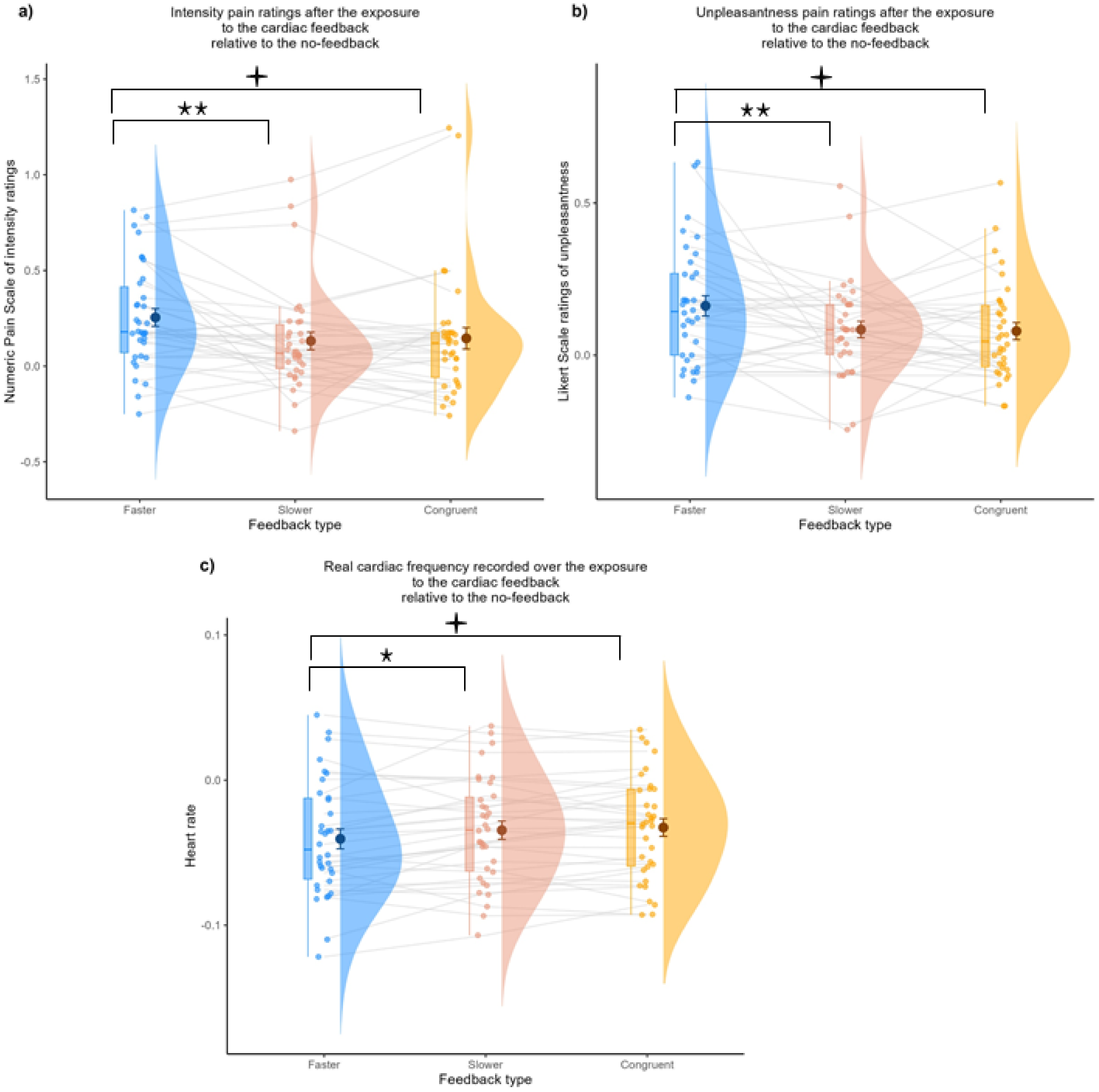
**a** Intensity pain ratings (Numeric Pain Scale), **b** Unpleasantness ratings (5-points Likert Scale), **c** Real cardiac frequency. Values consist of the central tendency of change of pain intensity (a), unpleasantness (b) and heart rate (c) associated with the exposure to the faster, slower and congruent cardiac feedback, relative to the no-feedback. Values of zero on the vertical axis would represent no change relative to the exposure to the no-feedback. Positive and negative values would represent an increase and a decrease, respectively. Brackets denote significance: = significant one-way RM-ANOVA; short bracket = planned Faster–Slower contrast. Asterisks indicate two-tailed *p* values (*< .05, ** < .01, *** < .001, “n.s.” = not significant).

### Subjective pain intensity ratings

Participants’ pain intensity ratings were analysed analogously. If faster heart rates are treated as an interoceptive signal of threat, then faster cardiac feedback should lead to increased pain intensity ratings compared to slower feedback. A repeated measures ANOVA with three levels (faster feedback, congruent feedback, slower feedback) revealed that the types of heart rate feedback affected pain intensity ratings, *F*(2, 66) = 7.017, *p* = .002, η*p²* = .175. A planned t-test then revealed that participants’ perception of pain intensity increased after faster relative to slower cardiac feedback, *t*(33) = 3.339, *p* = .002, *d* = .573 (Figure 2a). Additional t-tests showed that, like in the heart rate recordings, this difference was largely driven by the faster feedback. When compared to congruent feedback, significant differences emerged for faster feedback, *t*(33) = 2.66, *p* = .012, *d* = .457, but not for the slower feedback, which showed a slight numerical decrease, *t*(33) = .477, *p* = .636, *d* = .082.

### Subjective pain unpleasantness ratings

Analogous tests run on pain unpleasantness ratings replicated all findings. The initial ANOVA confirmed that the different types of cardiac feedback (faster, congruent, slower) affected pain unpleasantness ratings, F(2, 66) = 6.092, p = .004, η*p²* = .156. A paired sample t-test showed that pain unpleasantness ratings were higher after faster feedback than slower cardiac feedback, *t*(33) = 2.771, *p* = .009, *d* = .48 (Figure 2b). When compared to congruent feedback, significant differences emerged for faster feedback, *t*(33) = 3.045, *p* = .005, *d* = .522, but not for the slower feedback, which showed a slight numerical decrease, *t*(33) = .206, *p* = .838, *d* = .035.

### Body Perception Questionnaire

The Short form Body Perception Questionnaire (Poli et al., 2021), administered at the end of the experiment, allowed us to explore whether individual differences in body awareness and supradiaphragmatic reactivity were related to the extent to which participants developed (1) the perceptual illusion of pain (2) the physiological changes in their cardiac frequency. Overall, the average score obtained at the Body awareness subscale and Supradiaphragmatic subscale was 5.22 (*SD* = 1.62) and 2.05 (*SD* = 1.96), respectively.

We examined Pearson’s correlations between BPQ scores and changes in heart rate, pain intensity, and pain unpleasantness in response to the *faster vs. slower* feedback conditions - our main contrast of interest which was a key driver of effects. No significant associations were found between either BPQ subscale and the change in heart rate (all p > 0.05). Likewise, no correlations were observed between the BPQ subscales and changes in pain unpleasantness or pain intensity across these two conditions (all p > 0.05).

Overall, these findings do not suggest reliable association between individual differences in self-reported body perception and the perceptual or physiological effects of the experimental manipulation. Please note that the administration of this questionnaire was fully exploratory, as the current sample size is likely underpowered for robust individual difference analyses, which typically require substantially larger samples to detect meaningful associations (Hedge et al., 2018; Schönbrodt & Perugini, 2013).

### Experiment 2

Experiment 1 established that false accelerated cardiac feedback induced both an illusory increase in the perception of pain and a modulation in the pain-anticipatory related cardiac state (i.e., heart rate) towards the expected level of the noxious stimulus, based on the frequency rate of the cardiac feedback. Experiment 2 tests whether the same perceptual and physiological changes are observed if participants are exposed to non-interoceptive feedback (i.e., unrelated to biological human sounds), which should be less likely to induce expectations of threat. To address this question, we substituted the interoceptive cardiac feedback with an exteroceptive tone but kept the experiment otherwise identical. Thus, as in Experiment 1, participants’ pain intensity (i.e., Numeric Pain Scale), unpleasantness (i.e., Likert scale) ratings and cardiac frequency (i.e., ECG) were collected both in a no-feedback as well as in a feedback phase, which could be either congruent, faster or slower than participants’ individual heart rate recorded over the no-feedback phase. We reasoned that participants would have no expectation that an external acoustic sound signals a preparation of their body to a threatening, high painful stimulus. Consequently, perceptual and cardiac modulations associated with the feedback manipulation should be reduced or eliminated with exteroceptive feedback.

## Methods

### Participants

Thirty-five participants (mean age 25.17, SD = 4.02, 23 women) were recruited from Gabriele D’Annunzio University and the wider community. All were right-handed with normal or corrected-to-normal vision. Exclusion criteria were identical to Experiment 1. All gave written informed consent, were unaware of the purposes of the study, and were fully debriefed at the end of the experiment. Ethical approval from the local ethics board was obtained.

As in Experiment 1, participants were excluded if the correlation between the desired five levels of the nociceptive stimulus intensity and their ratings in the no-feedback condition was below *r*=0.75, suggesting insensitivity to the varying nociceptive degrees of the electrical stimulus. Five participants were excluded with this criterion. Moreover, participants were excluded if, for any reason, they linked the exteroceptive sound to a simulation of an interoceptive, cardiac acoustic feedback, in their post-experiment debriefing. One additional participant was excluded with this criterion. Hence, the final sample size was 29. A sensitivity analysis with G*Power 3.1 (Faul et al., 2007) showed that a sample size of 29 provides .90 power to detect effects with Cohen’s *d* >.62 (SESOI of δ = 0.37).

### Apparatus and Stimuli

The apparatus and stimuli were the same as Experiment 1. The only difference was the selection of the tone used to present the heart rate feedback. Instead of using a heartbeat, the sound was a single percussion obtained by knocking two woods, gathered from https://freesound.org and manipulated in Audacity. The feedback audio was then created in MATLAB (R2020a), repeating the single tone according to the desired frequency (see *Procedure*).

### Procedure

The procedure used in Experiment 2 was the same as Experiment 1, with the only exception that the sound used for the feedback was an exteroceptive, instead of interoceptive (i.e., cardiac) tone. Unlike in Experiment 1, where participants were told they would hear their own heartbeats in real time, in Experiment 2 participants were not informed that the sound was meant to represent their heart rate.

### Data Analysis

Data analysis was identical to Experiment 1. Note that we predicted an absence/reduction of effects here, compared to the findings of Experiment 1. In standard analyses, a non-significant result does not allow one to conclude that there is no effect; it simply indicates a lack of evidence for a statistically detectable difference. The predicted reduction/absence of effects with exteroceptive cues here relative to the interoceptive cues in Experiment 1 were tested in two ways.

First, to test for the predicted absence of effects with exteroceptive cues, equivalence tests (i.e., TOST ‘two one-sided t-tests’ procedure; Lakens, 2017; Lakens et al., 2018) were performed for the differences between each dependent (i.e., pain unpleasantness, pain intensity, heart rate). The TOST procedure addresses the above limitation by assessing whether a given (typically non-significant) effect is smaller than the effective smallest effect size of interest (SESOI) that the current design has power to detect, using two one-sided t-tests against positive and negative equivalence bounds. If both tests are statistically significant, it confirms that the differences in heart rate, pain intensity and pain unpleasantness induced by the target conditions (faster and slower feedback) are smaller than the smallest effect size of interest detectable in the current study.

Second, to confirm that the exteroceptively induced changes in heart rate and pain perception differ from those with interoceptive stimuli in Experiment 1, we compared our main contrast of interest - each participant’s difference between faster and slower feedback in each of the three dependent measures - between the two experiments, with independent-samples t-tests. A significant result from this t-test shows that the changes induced by faster vs slower exteroceptive feedback here differ from the analogous changes induced by interoceptive feedback in Experiment 1.

Finally, we checked whether the different sounds (interoceptive heartbeat sounds in Experiment 1, exteroceptive control sounds in Experiment 2) induced any overall changes in heart rates, painfulness or unpleasantness, which would be expected if these sounds differ systematically in the amount of arousal or alertness they induce, using between-group t-tests on overall heart rate, painfulness or unpleasantness averaged across all feedback conditions (equivalent to a main effect of Experiment in an ANOVA).

## Results

### ECG Heartrate measurement

As in Experiment 1, we first tested whether participants’ heartrate was affected by the feedback manipulation, now using non-interoceptive stimuli, using a 1-factor repeated measures ANOVA with three feedback levels (faster, congruent, slower). In contrast to Experiment 1, and as hypothesized, no robust differences between the three conditions emerged, *F*(2, 56) = 2.54, *p* = .088, η*p²* = .083. Our main hypothesis test, which directly compares cardiac frequency in trials with faster and slower feedback revealed no robust differences either, *t*(28) = 1.48, *p* = .317, *d* = .19. A TOST equivalence test was conducted to determine whether the difference in heart rate between the *Incongruent Faster* and *Incongruent Slower* conditions was statistically equivalent within a range of *d* = -0.57 to *d* = 0.57. The observed effect size was *d* = -0.19 (90% CI [-0.36, -0.03]). Both one-sided tests were significant, *t*(28) = 2.05, *p* = .025; *t*(28) = -3.71, *p* < .001, indicating statistical equivalence between conditions, relative to the smallest detectable differences in our design.

Note that, if anything, the induced difference in heart rate went in the numerically opposite direction than in Experiment 1, showing a heart rate *increase* after faster compared to slower exteroceptive feedback, in contrast to the decrease induced by interoceptive feedback cues. Indeed, comparing this increase to the equivalent change in Experiment 1 with a between-groups t-test revealed robust differences, *t*(61) = 2.187, *p* = .033, *d* = .553. Thus, while faster vs. slower interoceptive heart rate feedback led to compensatory changes in participants’ own heart rate, no such effects were obtained for exteroceptive feedback.

As in Experiment 1, we also investigated whether any differences were apparent when faster and slower feedback were compared to congruent feedback. No significant differences emerged for faster feedback *t*(28) = 1.202, *p* = .239, *d* = .22. However, the slower feedback induced a decrease *t*(28) = 2.291, *p* = .030, *d* = .43, again showing the opposite pattern than in Experiment 1.

Finally, we checked whether the interoceptive and exteroceptive stimuli in Experiments 1 and 2 induced overall changes in heart rate response, but no such differences were apparent, *t*(61) =.140, *p* = .889, *d* = .035.

### Subjective pain intensity ratings

Data were analysed as in Experiment 1. We first tested whether pain intensity ratings were generally affected by the feedback manipulation, using a 1-factor repeated measures ANOVA with the levels (faster feedback, congruent feedback, slower feedback). Consistent with the results of the heart rate data, there was no robust difference, *F*(2, 56) = 2.54, *p* = 0.088, η*p²* = .083. In particular, there was no difference in pain ratings between trials with faster and slower feedback, *t*(28) = .64, *p* = .53, *d* = .12, and a A TOST equivalence test was conducted to examine whether the difference in external NPS ratings between the *faster* and *slower* conditions was statistically equivalent within bounds of Cohen’s d = ±0.57. The observed effect size was *d* = -0.12 (89% CI [-0.34, 0.10]). Both one-sided tests were significant, *t*(28) = 2.43, *p* = .011; *t*(28) = -3.33, p = .001, indicating that the effect was statistically equivalent within the bounds of our experimental design.

When the difference in pain intensity ratings induced by faster and slower exteroceptive feedback was compared to that induced by interoceptive feedback in Experiment 1, a significant difference emerged, *t*(61) = 2.052, *p* = .044, *d* = .519. Thus, while faster (relative to slower) heart rate feedback increased pain ratings, delivering the same feedback through an exteroceptive stimulus induced no such changes.

Relative to congruent feedback, intensity ratings were increased for both faster and slower feedback, even though the difference was robust only for the faster feedback, *t*(28) = 2.12, *p* = 0.043, *d* = 0.39, not the slower cardiac feedback, *t*(28) = 1.58, *p* = 0.13, *d* = 0.29.

Finally, we checked whether the interoceptive and exteroceptive stimuli in Experiments 1 and 2 induced overall changes in pain intensity ratings, but no such differences were apparent, *t*(61) =.538, *p* = .593, *d* = .136.

### Subjective pain unpleasantness ratings

Pain unpleasantness ratings showed the same pattern as pain intensity ratings. First, a 1- factor repeated measures ANOVA with the levels faster feedback, congruent feedback, and slower feedback revealed no robust differences, *F*(2, 56) = 1.01, *p* = .371, η*p²* = .035. Moreover, there was no difference between trials with faster and slower feedback, *t*(28) = .864, *p* = .395, *d* = .16 (Figure 3b) and the A TOST procedure (Lakens, 2017) was conducted to test whether the difference in external liking ratings between the *faster* and *slower* conditions was statistically equivalent within bounds of Cohen’s d = ±0.57. The observed effect size was d = -0.02 (89% CI [-0.05, 0.02]). Both one-sided tests were significant, t(28) = 2.21, p = .018; t(28) = -3.56, p < .001, indicating statistical equivalence between the two conditions within the bounds of our experimental design.

**Figure 3.**
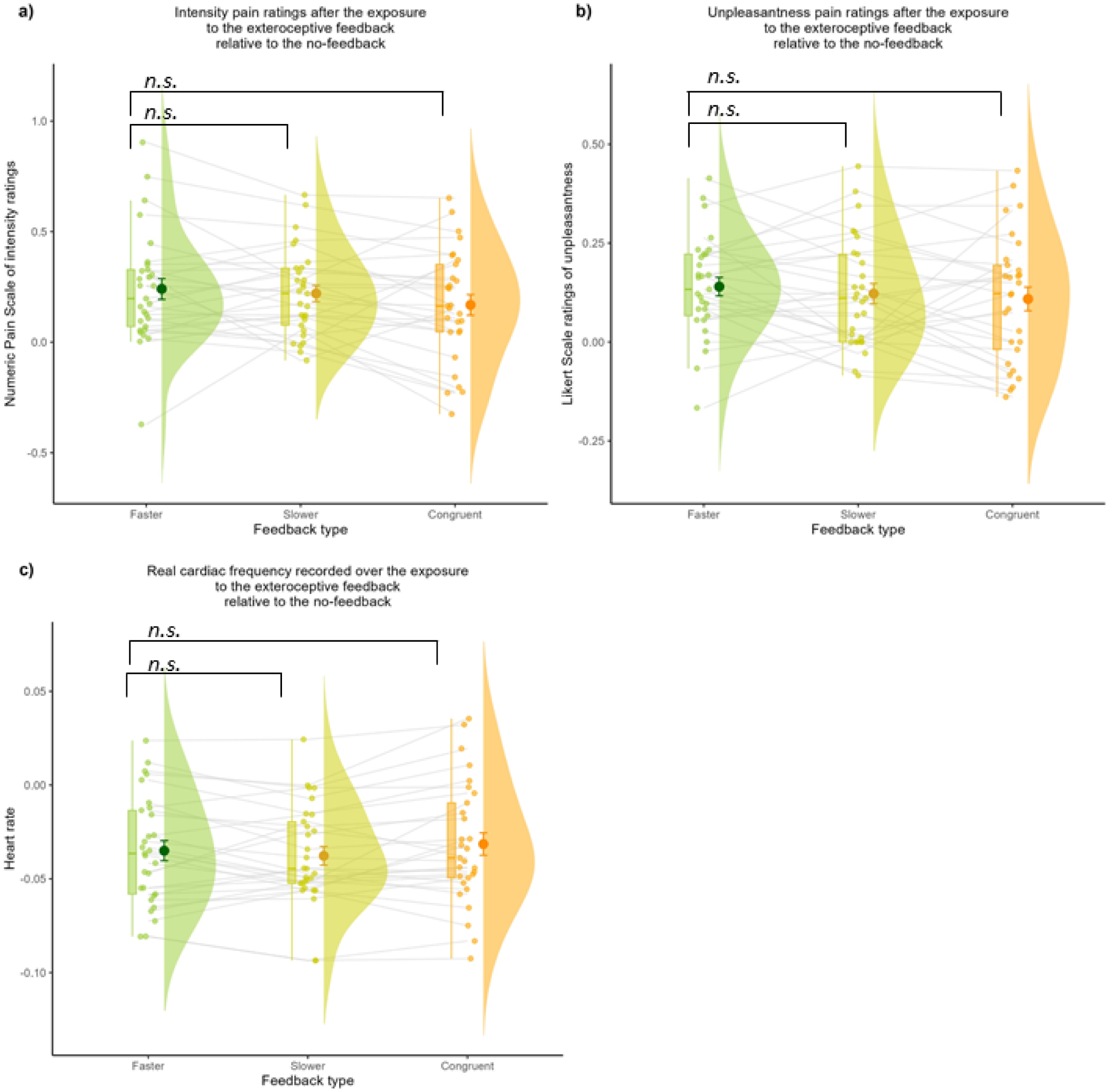
**a** Intensity pain ratings (Numeric Pain Scale), **b** Unpleasantness ratings (5-points Likert Scale), **c** Real cardiac frequency. Values consist of the central tendency of change of pain intensity (a), unpleasantness (b) and heart rate (c) associated with the exposure to the faster, slower and congruent exteroceptive feedback, relative to the no-feedback. Values of zero on the vertical axis would represent no change relative to the exposure to the no-feedback. Positive and negative values would represent an increase and a decrease, respectively. Brackets denote significance: = significant one-way RM-ANOVA; short bracket = planned Faster–Slower contrast. Asterisks indicate two-tailed *p* values (*< .05, ** < .01, *** < .001, “n.s.” = not significant).

Comparing these effects of exteroceptive feedback to the same stimulation delivered interoceptively in Experiment 1 revealed suggestive evidence for a difference in line with the previous measures, but it was not significant, *t*(61) = 1.679, *p* = .098, *d* = .424.

Further simple t-tests showed that, relative to the congruent feedback, differences did neither emerge for faster feedback, *t*(28) = 1.428, *p* = .164, *d* = .27, nor for the slower feedback, *t*(28) = .571, *p* = .573, *d* = .06.

Finally, we checked whether the interoceptive and exteroceptive stimuli in Experiments 1 and 2 induced overall changes in pain unpleasantness ratings, but no such differences were apparent, *t*(61) =.435, *p* = .665, *d* = .110.

### Supplementary analyses

To rule out potential confounds and confirm the robustness of our findings, we conducted additional analyses on the raw data (see *Supplementary materials*), using the mixed linear models approach, which offer increased power and reliability with limited trials and conditions (Kliegl et al., 2011; Baayen et al., 2010; Ambrosini et al., 2019). These analyses included all levels of feedback for heart rate and pain ratings (unpleasantness and intensity), stimulus intensity levels, as well as trial number (i.e., the rank order of each trial) as a variable to assess potential time-related effects such as learning, adaptation, or fatigue (e.g., Möckel et al., 2015). In addition, we incorporated the initial no-feedback phases as condition to test for potential baseline differences, and we analyzed all stimulus intensities separately to test whether feedback effects were specific to particular nociceptive levels or generalized across them. These analyses fully replicate the findings of the main analysis reported above, but offer more detailed insights for interested readers into other, not theoretically relevant, factors that may affect the results. Due to the large number of comparisons, and the associated alpha inflation, these additional, not a priori predicted, results should be interpreted with caution, however, before being replicated in further studies.

### General Discussion

Interoceptive inference and the Embodied Predictive Coding frameworks (Barrett, 2017; Barrett & Simmons, 2015; Pezzulo, 2014; Seth, 2013; Seth & Friston, 2016) propose that interoception is not merely a passive mirror of the internal milieu but rather the outcome of active, probabilistic inferences. Through this mechanism, the brain constantly estimates and regulates homeostatic states by integrating prior expectations with incoming interoceptive, proprioceptive and exteroceptive signals. To test this framework in the context of pain, participants were given auditory cardiac feedback, which they believed reflected their actual physiological state, before the administration of pain stimuli.. Unbeknownst to them, we varied the frequency of this heart rate feedback, making it either faster or slower. This manipulation aimed to create a false interoceptive prior, suggesting an increase or decrease in heart rate just before they received a painful stimulus.

In predictive models of interoception, such differences between expected and actual interoceptive information can be reduced by both changing one’s perceptual state to encompass the new information, and by actively changing the body in response to the unexpected input, by engaging autonomous reflexes (i.e., perceptual and active inference, Parr et al., 2022, Friston, 2005; 2010; Ainley et al., 2016, Paulus et al., 2019). If so, then sensing an accelerated heart rate (relatively to a decelerated one) should have two consequences: first, it may falsely lead participants to expect, and then perceive, a more intense pain stimulus then is really delivered. Second, it may induce changes in autonomic states themselves, compensating for the perceived changes in heart rate by actually down-regulating the real heart rate, towards the expected decreasing orienting cardiac response when expecting threat (Lykken et al., 1972; Taggart et al., 1976; Bradley et al., 2005; Bradley et al., 2008; Colloca et al., 2012; Tracy et al., 2017).

Experiment 1 provides evidence for both components of the hypothesized response. It showed, first, that faster cardiac feedback induced an illusory perception of increased unpleasantness and intensity of subsequent pain stimuli compared to slower cardiac feedback. Thus, as predicted, the mismatch between people’s expectations and the actual input led to changes in the perception of pain, towards the *expected* input associated with increased heart rates, in line with Bayesian/predictive processing accounts of pain perception. Second, false cardiac feedback also changed participants’ real heart rate. When hearing faster (relative to slower) feedback, the real heart rate decreased, assuming the pattern of anticipatory compensatory response typically enacted when humans prepare to face a threatening stimulus (Lykken et al., 1972; Taggart et al., 1976; Bradley et al., 2005; Bradley et al., 2008; Colloca et al., 2006; Tracy et al., 2017).

Importantly, both perceptual and cardiac frequency changes after faster (vs. slower) feedback were not observed when the heartbeat tones were replaced by exteroceptive sounds in Experiment 2. This finding supports the hypothesis that the observed perceptual and physiological modulations are specifically tied to interoceptive feedback, which provides evidence of the current state of the body in preparation for a potential upcoming (threatening) stimulus. In contrast, as the frequency of an external sound is not informative about the current state of the body, it should not induce any nociceptive stimulus expectations. Indeed, no discrepancy in the pain unpleasantness and intensity ratings, as well as in participants’ actual heart rates, were observed in Experiment 2. Importantly, the change from interoceptive to exteroceptive feedback did not induce any overall changes in pain perception or heart rate between experiments, only affecting the sensitivity to its rate (i.e., faster or slower). This suggests that the two types of cues were otherwise similarly processed, and did not vary substantially in overall arousal or sensitivity to pain they induced by themselves.

To our knowledge, this is the first study to successfully demonstrate that both *perceptual* and *active inference* components of pain are modulated through false interoceptive feedback. This provides evidence for the proposed mechanism through which, based on prior knowledge and expectations, the organism infers and actively implements the interoceptive consequences that have been predicted (Barrett & Simmons, 2015; Pezzulo, 2014; Seth & Friston, 2016). Our findings therefore support predictive coding accounts of interoception, which propose such effects on perception and bodily state, and extend them beyond exteroceptive cues (visual, Wiech et al., 2014; Jepma et al., 2019; auditory, Colloca et al., 2006; Atlas et al., 2010). They indicate that cardiac signals are weighted in the brain’s generative model of pain and serve as key variables in allostatic regulation (Barrett et al., 2016), driving both the perception of further interoceptive signals and the regulation of adaptive bodily states.

A conceptual parallel can be drawn with the Rubber Hand Illusion (RHI), where mismatches between seen and felt touch are resolved both perceptually and through autonomic adjustments (Botvinick & Cohen, 1998; Moseley et al., 2008). Crucially, these responses can emerge from expectations alone (Ferri et al., 2013). Similarly, here, faster cardiac feedback created a prediction of threat that was resolved perceptually as higher pain ratings and actively as a parasympathetic deceleration of heart rate. Thus, just as the RHI illustrates predictive integration of visual and proprioceptive signals, the present findings reveal similar mechanisms in the visceral domain, showing that interoceptive predictions about cardiac state can reshape both the subjective and autonomic dimensions of pain.

An interesting observation was that, when compared to congruent cardiac feedback, the current effects on pain perception and autonomic responses were mainly driven by accelerated, not decelerated, feedback. Similar asymmetries of faster cardiac feedback have been reported in other domains. For instance, Iodice et al. (2019) showed that participants overestimate the effort they exerted when provided with faster cardiac feedback, whereas slower feedback had no effect. We speculate that, here, the asymmetry arises because increased heart rates are a biologically conserved (and culturally reinforced) cue for arousal, threat, and pain, which occurs for a wide range of stressors. Faster feedback therefore, conveys the expectation of pain by mimicking the bodily response that follows its actual experience (i.e., heart-rate acceleration). In contrast, a slowed heart rate is the body’s intended autonomic response when *expecting* (but not yet experiencing) pain in both human and non-human species, to prepare for the forthcoming stressor (for reviews, see Livermore et al., 2021; Roelofs, 2017; Vila et al., 2007). It is considered an evolutionarily conserved, adaptive response that allows the organism to orient attention, enhance information processing and action preparation (Hashemi et al., 2019; Klaassen et al., 2021; Lojowska et al., 2015). In our case, therefore, decelerated cardiac feedback signals that this intended autonomic response for dealing with the anticipated pain stimulus is already achieved. An accelerated heart rate, in contrast, signals the opposite, providing interoceptive evidence of a misaligned bodily state that mandates immediate allostatic responses, in terms of both pain perception and cardiac regulation (Barrett & Simmons, 2015; Pezzulo, 2014; Seth & Friston, 2016).Our findings therefore blend well with recent proposals that frame cardiac deceleration under a Bayesian inference approach. Under these views, the cardiac deceleration mechanism allows to adjust precision of sensory evidence accumulation relative to precision of bodily information, optimising perception and action (Skora et al., 2022). Thus, decelerated cardiac activity when exposed to the faster feedback may reflect an autonomic adjustment to prioritize incoming noxious input information in light of the expectation of the threatening painful event.

Together, our evidence therefore supports the hypothesis that not only interoceptive states modulate interoceptive inference and perception, but the converse is also true: inference dynamics can modify or produce new internal states, to the extent that the actual cardiac frequency rate decreased as it would have done over the anticipation of a real threat. More broadly, our findings are in line with recent proposals of *interoceptive schemas* (Barca & Pezzulo, 2020; Iodice et al., 2019; Schoeller et al., 2022; Tschantz et al., 2022), internal models akin to the body schema (Head & Holmes, 1911), which consist of a central representation of interoceptive variables (e.g., body temperature, cardiac activity etc.) along with prior assumptions or "set points" for these variables. This internal model would support homeostatic and allostatic regulation by optimally weighting multiple (i.e., proprioceptive, interoceptive, exteroceptive) streams of information to predict incoming interoceptive signals (Barca & Pezzulo, 2020; Iodice et al., 2019; Schoeller et al., 2022; Tschantz et al., 2022). Interoceptive illusions, i.e., contextual manipulations that shift the precision (i.e., estimated reliability) of interoceptive expectations (Schoeller et al., 2024), offer a direct probe of this schema. Our paradigm is one of such manipulation, showing how a transient change in cardiac evidence reveals how the precision weighting of priors and their prediction errors reshapes both the perceptual and autonomic dimensions of pain. Such illusions open a window onto the internal generative models that guide perception and action. In this regard, the contribution of the study of illusions offers important insights not only for a converging model of brain functioning under the principle of optimal predictive architectures, but it also provides implications for understanding psychopathology in terms of aberrant predictions and compounding allostatic consequences (for a full discussion, see Barrett & Simmons, 2015). Because manipulating contextual cues selectively alters the precision-weighting of interoceptive expectations, such illusions constitute the kind of diagnostic “stress tests” that reveal how much confidence the brain assigns to priors versus bodily evidence, a parameter now believed to underlie dysfunctional conditions, such as anxiety, PTSD and chronic pain (Schoeller et al., 2022). Our experimental paradigm provides a probe of this mechanism: the coupled shift in pain ratings and heart-rate deceleration indexes the degree to which threat-related priors dominate over incoming cardiac signals. Emerging perspectives (Schoeller et al., 2022) frame precisely this imbalance (i.e., excessively precise priors or under-weighted sensory channels) as a form of *false inference* that underlies psychopathology, and identify interoceptive illusions as tools both for phenotyping such aberrant precision control and for retraining it through repeated, controlled exposure.

### Accounting for general unspecific contributions

One possibility is that the different effects in Experiment 1 and 2 may not reflect interoceptive vs exteroceptive processing, but more general features of the sounds used in both experiments. For example, the heartbeat sounds’ cultural meaning and salience (e.g., as signals of anxiety or distress), or their acoustic properties (e.g., loomingness, roughness), could have increased arousal, alertness or attentional engagement more generally. Several aspects of our data argue against this proposal, however. First, if heartbeat sounds had such general alerting/arousing effects, they should have induced robust between-experiment differences in pain ratings or heart rates when such heartbeat (Experiment 1) or non-heart related sounds were presented (Experiment 2). However, no such differences were obtained for any of our measures (see between-experiment t-tests in the main analyses; cross-experiment analysis in supplementary analyses). In contrast, all effects we obtained reflected not the overall presence or absence of cardiac feedback, but responses to a *change* in cardiac feedback (faster or slower), which was absent for identical changes in exteroceptive feedback.

Second, one may suspect that the acceleration of cardiac feedback, rather than cardiac feedback more generally, had simply increased arousal, instead of functioning as a misaligned interoceptive signal. However, if so, one would expect to see an overall *acceleration* in participants’ actual heart rate, as this is a typical physiological marker of heightened arousal (Azarbarzin et al., 2014; Wascher, 2021; Yang et al., 2017), not the slowing of the heart rate we had predicted and observed. Thus, our findings are *opposite* to what would be expected under a generic arousal-based explanation but consistent with the well-known anticipatory physiological regulatory response predicted by interoceptive inference frameworks. Note also that for such changes in heart rate to be obtained, very strong manipulations of threat or arousal are required. For example, in Parrotta et al. (2024), even highly salient and reliable threat cues (i.e., images that predicted upcoming pain with 100% certainty) did not produce any measurable change in heart rate. Our pattern of results is therefore not readily accounted for by general arousal or salience mechanisms, but is fully in line with the intrinsic capacity of cardiac feedback to engage autonomic regulation.

Finally, the idea that the interoceptive relevance of the cue, rather than its low-level acoustic properties or general human-related salience, is the key driver of the observed effects is also reflected in neurophysiological studies. For example, fMRI studies have shown that heartbeat sounds selectively activate interoceptive regions (e.g., anterior insula, frontal operculum; Kleint et al., 2015), while EEG work (Vicentin et al., 2024) has demonstrated enhanced cortical activation to faster heartbeats, particularly over frontocentral regions, suggesting enhanced processing in networks associated with interoceptive attention. Moreover, heartbeat sounds have been shown to attenuate the auditory N1 component (van Elk et al., 2014), a neural signature typically linked to self-generated or predicted bodily signals.

### Limitations and future directions

The current results provide a platform for further investigation of the link between perceptual and active inference in interoception. As this was the first study using our paradigm, it was optimised to detect robust within-subject effects of pain expectation. By definition, such robust within-subject effects minimize between-participant differences and are therefore difficult to link to cross-participant variability (i.e., the “reliability paradox”, Hedge, Powell & Sumner, 2018), requiring substantially larger samples than used here and in the literature (i.e., N>200). Potentially informative associations, such as those between anticipatory heart-rate dynamics and pain ratings, could not be tested here. Similarly, the absence of a robust relationship between pain modulation and BPQ subscales of body awareness and supradiaphragmatic reactivity should be interpreted with caution, given that these measures were included for exploratory purposes and not sufficiently powered to draw strong conclusions, especially about the absence of relationships. Future work designed and powered for individual differences should determine how interoceptive awareness and cardiac attention shape pain experience. Such studies could also investigate how the current findings depend on sex/gender. In our sample, like in most psychology research, women were somewhat over-represented. Prior research suggests that men and women differ in pain processing and interoceptive sensitivity (Sorge et al., 2018; Mogil, 2012). Future studies with larger samples could investigate whether such differences modulate the integration of interoceptive signals and pain perception, especially as predictive processing accounts assume that the precision of sensory information determines how much it is weighted against prior expectations (Yon & Frith, 2021).

A promising avenue is integrating perceptual, neuroimaging and physiological measures (e.g., the Heartbeat Evoked Potential, Petzschner et al., 2019) into computational models of interoceptive inference (e.g., Allen & Friston, 2018; Owens et al., 2018; Allen et al., 2022; Eckert et al., 2022). Similar inferential frameworks as proposed here have been implemented in computational models of cardiac (Smith et al., 2020) and gastrointestinal (Smith et al., 2021) interoception, thermoregulation (Tschantz et al., 2022), and psychopathological conditions, such as anorexia nervosa (Barca & Pezzulo, 2020), panic disorder (Maisto et al., 2021), self-injury (Barca et al., 2023), depression (Barrett et al., 2016; Stephan et al., 2016), and others (Paulus et al., 2019). Such frameworks would allow latent parameters like prior precision and sensory gain to be estimated for each participant (Unal et al., 2021), allowing us to test whether individuals who rely more heavily on cardiac priors also exhibit larger pain illusions and autonomic adjustments, further enabling individual differences research and extending it to clinical applications.

Model comparison could also establish whether faster cardiac feedback shifts the mean of the pain prior or instead alters the precision of nociceptive evidence. Systematically manipulating the precision of priors or prediction errors, or explicitly measuring participants’ interoceptive beliefs and their precision, offers a powerful means to uncover the mechanisms that facilitate the development of the interoceptive illusion. They may reveal whether perceptual biases in the experience of pain emerge from overlearned priors that do not update to incoming sensory data, or whether it is possible to directly act on interoceptive illusions and their autonomic consequences by changing prior knowledge.

The latter would have far-reaching implications, making it possible to both modify the perception of our internal milieu and impact autonomic states by simply reconfiguring individuals’ beliefs and knowledge. Such findings would have profound consequences for understanding and potentially intervening in psychopathological conditions. On this basis, manipulating contextual cues can selectively up- or down-weight interoceptive expectations. In our paradigm, the coupled shift in pain ratings and heart-rate deceleration quantifies the extent to which threat-related priors override incoming bodily signals (i.e., cardiac input), a precision parameter that recent accounts frame as the very locus of dysfunctional predictive processes in anxiety, PTSD and chronic pain.

From this perspective, the pain illusion documented here, together with other interoceptive illusions, provide a direct window into the brain’s predictive architecture (Barrett & Simmons, 2015), offering a mechanistic framework for understanding psychopathology as a failure to revise inaccurate interoceptive priors, leading to maladaptive regulation and compounding allostatic consequences. At the same time, they serve as diagnostic “stress tests,” revealing how much confidence the brain assigns to priors versus bodily evidence and thereby offering a mechanistic foothold for targeted intervention. Finally, we note that in our unpleasantness scale, the lowest anchor was labeled “no pain.” Although this label was not ideal, as unpleasantness and pain can in principle be dissociated, the effect of this choice is likely minimal given that unpleasantness ratings were always provided after intensity ratings, and our main findings rely on the interplay between intensity measures and physiological cardiac modulations.

## Conclusions

This study is the first to show that false cardiac signals reshape both pain perception and anticipatory autonomic responses. False cardiac feedback amplified pain intensity and unpleasantness while inducing parasympathetic deceleration of heart rate, revealing how expectations actively construct interoceptive experiences and drive allostatic adjustments. These findings support predictive models by emphasizing the pivotal role of interoceptive schemas in predictive architectures and highlight cardiac signals as key variables in the generative models governing pain and autonomic regulation. Importantly, the lack of effects with non-interoceptive feedback highlights the specificity of this mechanism to cardiac interoceptive signals, providing a platform to explore whether other visceral channels, such as respiratory cues, similarly recalibrate pain perception and physiological states. Beyond advancing theoretical models of interoception, our paradigm offers a novel tool for probing the precision-weighting of priors and sensory evidence in healthy and clinical populations, which is consistent with proposals that interoception provides actionable targets for mental-health interventions (Nord & Garfinkel, 2022). Future work may explore its translational potential in understanding and modulating maladaptive pain experiences, where aberrant interoceptive predictions contribute to chronic pain and related disorders.

## Acknowledgements

The work was funded by Leverhulme Trust grant RPG-2019-248 to Patric Bach, and PhD studentship awarded to Eleonora Parrotta from the Universities of Plymouth and Aberdeen. This research received funding from the European Union’s Horizon 2020 Framework Programme for Research and Innovation under the Specific Grant Agreement 945539 (Human Brain Project SGA3) and 952215 (TAILOR) to Giovanni Pezzulo, the European Research Council under the Grant Agreement No. 820213 (ThinkAhead) to Giovanni Pezzulo and the PNRR MUR project PE0000013-FAIR to Giovanni Pezzulo. This work was supported by the "Departments of Excellence 2018-2022" initiative of the Italian Ministry of Education, University and Research for the Department of Neuroscience, Imaging and Clinical Sciences (DNISC) of the University of Chieti-Pescara, and by the "Search for Excellence" initiative of the University of Chieti-Pescara.

Below we list the main authors’ contribution, in accordance with the CRediT taxonomy:

Eleonora Parrotta: Conceptualization, Methodology, Software, Formal Analysis, Investigation, Writing - Original Draft, Visualization.

Patric Bach: Conceptualization, Methodology, Writing – Original Draft, Resources, Supervision, Project Administration, Funding Acquisition.

Giovanni Pezzulo: Conceptualization, Methodology, Writing - Review & Editing, Resources, Supervision, Funding Acquisition.

Andrea Zaccaro: Methodology, Software, Formal Analysis, Writing - Review & Editing

Mauro Gianni Perrucci: Methodology, Software, Formal Analysis, Investigation, Writing - Review & Editing

Marcello Costantini: Conceptualization, Methodology, Writing - Review & Editing, Supervision, Resources, Project Administration, Funding Acquisition.

Francesca Ferri: Conceptualization, Methodology, Writing - Review & Editing, Supervision, Resources, Project Administration, Funding Acquisition.

## Data availability

The data used for the analyses is available upon publication in an open repository at the following link: https://github.com/eleonorap17/painperception. The behavioural and physiological raw data can be shared by the corresponding author upon request if data privacy can be guaranteed according to the rules of the European General Data Protection Regulation (EU GDPR).

