## Supplementary materials for "Exposure to false cardiac feedback alters pain perception and anticipatory cardiac frequency"

Supplementary material	1

Linear mixed-effects model	1

1.Cross-experiment analysis (between-subjects model)	2

1.1.Heart rate	2

1.2 Likert Pain Unpleasantness Ratings Scale	5

1.3 Numeric Pain Scale of intensity ratings	9

2. Interoceptive experiment (within-subjects model)	13

2.1 Heart rate	13

2.2 Likert pain unpleasantness	15

2.3 Numeric Pain Scale of intensity ratings	20

3.Exteroceptive experiment (within-subjects model)	24

3.1 Heart rate	24

3.2 Likert pain unpleasantness	25

3.3 Numeric Pain Scale of intensity ratings	30

References	34

### Supplementary material

#### Linear mixed-effects model

We used linear mixed-effects models to investigate the effects of Feedback, Stimulus Intensity, and Trial on our dependent measures (Numeric Pain Scale intensity ratings, Likert ratings, and heart rate). In all models, the variable Trial was included as a continuous predictor reflecting the relative position of each trial within the session. Trial was linearly rescaled within each participant to range from –0.5 (first trial) to +0.5 (last trial), using min–max normalization. This transformation enables consistent interpretation of temporal effects across sessions of different lengths and aligns with best practices in linear mixed-effects modeling (Meteyard & Davies, 2020; Schad et al., 2020).

Stimulus Intensity (StimInt) was originally derived from three discrete stimulation levels (3, 4, and 5), corresponding to subjective pain ratings of NPS 30, 50, and 70, respectively. These levels were selected to induce variability in pain perception while avoiding floor and ceiling effects, in line with previous work on pain illusions (Atlas et al., 2014; Colloca et al., 2006; Hird et al., 2019). For statistical modeling, these levels were recoded as a continuous centered variable: –0.5 (low = NPS 30), 0 (medium = NPS 50), and +0.5 (high = NPS 70), allowing for linear trend analysis.

Feedback was modelled as a categorical factor with four levels (Congruent, Slower, Faster, No Feedback), and Experiment (Exp) was included as a fixed factor in between-subject models comparing the interoceptive and exteroceptive conditions (Exteroceptive, Interoceptive).

To increase the robustness of our analyses, we excluded outliers based on standardized residuals: observations with residuals exceeding ±2 standard deviations were removed prior to model fitting (Baayen & Milin, 2010). Residual diagnostics were then performed by visually inspecting quantile–quantile (Q–Q) plots to assess the normality assumption.

Random effects structures included by-subject random intercepts and random slopes for both Trial and Stimulus Intensity (StimInt), following recommendations for maximizing model generalizability (Barr et al., 2013). For each model, we compared the full random-effects structure to simpler models using likelihood ratio tests, and retained random slopes only when they significantly improved model fit (see also Matuschek et al., 2017).

We did not include *Experiment* (Exp) or *Feedback* as random effects in our models. *Experiment* was a between-subjects factor reflecting two distinct, theoretically defined groups (Interoceptive vs. Exteroceptive). Because participants were nested within a single experiment, this factor lacked within-subject variability and was therefore modeled as a fixed effect. This approach is consistent with standard practices for modeling between-subject manipulations that represent specific experimental conditions rather than a sample from a broader population (Barr et al., 2013; Brauer & Curtin, 2018).

Similarly, *Feedback* was modeled as a fixed effect rather than a random factor. This within-subject manipulation comprised four experimentally designed levels (Congruent, Slower, Faster, No Feedback), each corresponding to a distinct prediction condition derived from our theoretical framework. These levels do not represent a random sample from a larger population of feedback types, but rather a set of conditions defined a priori based on predictive coding and interoceptive inference models. As such, modeling *Feedback* as a random factor would violate key assumptions of hierarchical modeling, while also obscuring the interpretation of fixed effects and their interactions. In line with best practices in mixed-effects modeling (Barr et al., 2013; Matuschek et al., 2017), random effects should be specified only for factors whose levels are sampled from a larger population, which was not the case in our design.

While some modeling approaches adopt a stepwise procedure, adding or removing predictors to assess incremental improvements in model fit, our models were not constructed in this way. Instead, we specified a full-factorial fixed-effects structure a priori, in order to comprehensively test whether the originally observed effects of feedback on pain ratings and heart rate remained robust when including all levels of Feedback (including No Feedback and Congruent) and all levels of the effective stimulus intensity. Although these analyses go beyond the scope of the original confirmatory models, they were conducted in part in response to a reviewer’s request for a more detailed examination of variability across trials and conditions. Their structure, however, was determined by theoretical considerations rather than by data-driven model selection. This approach aligns with best practices in mixed modeling that prioritize hypothesis-driven model specification and theoretical transparency over purely exploratory, stepwise procedures (Gelman & Hill, 2007; Barr et al., 2013; Harrell, 2015). Moreover, stepwise model building can be useful in exploratory contexts, but it has also been criticized for inflating Type I error rates, encouraging overfitting, and obscuring interpretability (Harrell, 2015). In contrast, *a priori* model specification allows for clearer hypothesis testing, maintains the integrity of confirmatory analyses, and ensures that all theoretically relevant terms, including interactions, are directly modeled. Accordingly, our models included the full factorial structure of fixed effects necessary to test the hypothesized interactions (e.g., Exp × Feedback × Trial), and random effects were tested and retained based on improvements in model fit via likelihood ratio tests (Matuschek et al., 2017).

All models were fitted using restricted maximum likelihood estimation (REML), and p-values were computed using Satterthwaite’s approximation of degrees of freedom. For each fixed effect, we report the estimated coefficient (b), standard error (SE), t-statistic (t), and p-value (p).

#### 1. Cross-experiment analysis (between-subjects model)

##### 1.1. Heart rate

To examine heart rate (HR) variations across experimental conditions, we applied general linear mixed-effects modeling (GLMM), which accounts for both between- and within-participant variability by estimating fixed effects while allowing subject-specific random variations. The model was fitted using **R** (lme4 package; Bates et al., 2015) and was specified as follows:

𝐻𝑅∼𝐸𝑥𝑝𝑒𝑟𝑖𝑚𝑒𝑛𝑡×𝐹𝑒𝑒𝑑𝑏𝑎𝑐𝑘×𝑇𝑟𝑖𝑎𝑙+(𝑇𝑟𝑖𝑎𝑙∣𝑆𝑢𝑏𝑗𝑒𝑐𝑡)

where Experiment, Feedback, and Trial were included as fixed effects. A random slope for Trial was included at the subject level (Trial | Subject) to account for individual differences in temporal HR changes. Model comparison confirmed that including the random slope for Trial significantly improved model fit (χ² = 11.462, p = 0.003), indicating that the trajectory of HR changes over time varied across participants.

###### 1.1.2 Results

###### 1.1.2.1 Effects of interest

The results of the Type III ANOVA with Satterthwaite’s approximation revealed a significant **main effect of Feedback**, *F* (3,1305.28) = 102.68,*p*<0.001, indicating that HR varied across feedback conditions. Compared to the reference condition (i.e., Congruent feedback), HR was significantly higher in the No Feedback condition (*b*=1.97, *SE*=0.24, *t*=8.21, *p*<0.001), significantly lower in the Faster feedback condition (*b*=-1.10, *SE*=0.24, *t*=-4.55, *p*<0.001), and showed a non-significant decrease in the Slower feedback condition (*b*=-0.42, *SE*=0.24, *t*=-1.76, *p*=0.079). *Post hoc comparisons* further clarified these effects. HR was significantly lower in the Faster feedback condition compared to Congruent feedback (*p*=0.0008), and significantly higher in the No Feedback condition compared to all other conditions (*p*<0.0001 for all comparisons). The Slower feedback condition did not differ significantly from the Faster condition (*p*=0.216), but showed a significant reduction in HR compared to the No Feedback condition (*p*<0.0001). These results suggest that the absence of feedback elicited the highest HR, while faster feedback was associated with the lowest HR.

Crucially, the **Feedback × Experiment** interaction was also significant, F(3, 1305.28) = 3.58, p = .013, indicating that the effect of Feedback on HR differed between the Interoceptive and Exteroceptive experiments.

Post hoc comparisons confirmed that in the *Interoceptive experiment*, heart rate (HR) in the No Feedback condition (*M* = 76.2, *SE* = 1.45) was significantly higher than in all other feedback conditions. Specifically, HR was significantly higher than in the Faster condition (*M* = 73.1, *SE* = 1.45; *estimate* = -3.07, *SE* = 0.24, *t*(1310.3) = -12.94, *p* < .0001), in the Slower condition (*M* = 73.8, *SE* = 1.45; *estimate* = -2.38, *SE* = 0.23, *t*(1305.5) = -10.31, *p* < .0001), and in the Congruent condition (*M* = 74.2, *SE* = 1.45; *estimate* = -1.96, *SE* = 0.24, *t*(1307.1) = -8.16, *p* < .0001). Importantly, HR in the Faster condition was significantly lower than in the Slower condition (*estimate* = 0.68, *SE* = 0.23, *t*(1309.2) = 2.92, *p* = .0043) and than in the Congruent condition (*estimate* = 1.10, *SE* = 0.24, *t*(1307.6) = 4.55, *p* < .0001) .

In the *Exteroceptive experiment*, HR in the No Feedback condition (*M* = 78.8, *SE* = 1.57) was also significantly higher than in all other feedback conditions. It was significantly higher than in the Congruent condition (*M* = 76.6, *SE* = 1.57; *estimate* = -2.22, *SE* = 0.25, *t*(1305.9) = -8.74, *p* < .0001), the Slower condition (*M* = 76.2, *SE* = 1.57; *estimate* = -2.58, *SE* = 0.25, *t*(1306.4) = -10.26, *p* < .0001), and the Faster condition (*M* = 76.5, *SE* = 1.57; *estimate* = -2.33, *SE* = 0.25, *t*(1306.0) = -9.25, *p* < .0001). However, in contrast to the Interoceptive experiment, no significant differences emerged between the Congruent, Slower, and Faster feedback conditions (Congruent vs. Slower: *estimate* = 0.36, *SE* = 0.25, *t*(1302.4) = 1.45, *p* = . 2204; Congruent vs. Faster: *estimate* = 0.11, *SE* = 0.25, *t*(1302.5) = 0.42, *p* = .674; Slower vs. Faster: *estimate* = -0.26, *SE* = 0.25, *t*(1301.6) = -1.04, *p* = . 3566).

Importantly, the crucial difference in HR between the Slower and Faster feedback conditions also differed significantly across experiments (*b* = 0.941, *SE* = 0.34, *t* = 2.76, *p* = .005). Specifically, the Slower–Faster contrast was larger in the Interoceptive experiment, indicating a more pronounced reduction in HR from the Slower to the Faster condition when participants were exposed to the interoceptive relative to the exteroceptive feedback. Moreover, interaction contrasts also revealed that the difference in HR between the Faster and No feedback conditions varied significantly across experiments, with a greater HR reduction in the Interoceptive experiment compared to the Exteroceptive one (b = -0.7411, SE = 0.346, t = -2.145, p = 0.032). No other between-experiment differences reached significance. Finally, interaction contrasts revealed that the difference in HR between the Faster and Congruent feedback conditions varied significantly across experiments (*b* = 0.998, *SE* = 0.35, *t* = 2.86, *p* = .004), with a greater HR reduction in the Interoceptive experiment compared to the Exteroceptive one.

###### 1.1.2.2 Additional unpredicted effects

Due to the large number of comparisons the results here should be treated with caution before being replicated.

Although the **main effect of Trial** did not reach significance, F(1, 64.44) = 3.51, p = 0.066, the fixed-effect estimate indicated a general increase in HR across trials (b = 2.89, SE = 0.71, t = 4.08, p < 0.001).

A significant interaction between **Trial × Experiment** emerged, *F*(1, 64.44) = 5.65, *p* = .020, indicating that the trajectory of heart rate (HR) over time differed between the two experiments. HR increased significantly across trials in the Interoceptive experiment, *b* = 1.40, *SE* = 0.45, 95% CI [0.51, 2.29]. In contrast, no significant change in HR over time was observed in the Exteroceptive experiment, *b* = –0.17, *SE* = 0.48, 95% CI [–1.13, 0.80]. The difference between these temporal trends was statistically significant, *b* = 1.57, *SE* = 0.66, *t*(64.8) = 2.38, *p* = .020, with a more positive HR slope in the Interoceptive than in the Exteroceptive experiment.

Moreover, the three-way interaction between **Trial × Feedback × Experiment** was significant, *F*(3, 1309.10) = 9.44, *p* < 0.001, indicating that the trajectory of HR over time depended on both feedback type and experimental condition.

In the *Interoceptive experiment*, HR significantly increased over trials in the Congruent feedback condition (*b* = 2.89, *SE* = 0.71, 95% CI [1.50, 4.29], *t*=4.07, *p*=0.0001), as well as in the Faster condition (*b* = 2.66, *SE* = 0.64, 95% CI [1.41, 3.92], *t*=4.171, *p*<0.0001). In contrast, no significant increase was observed in the Slower condition (*b* = 0.66, *SE* = 0.61, 95% CI [–0.55, 1.86], *t*=1.072, *p*=0.285), and HR even showed a non-significant decreasing trend in the No Feedback condition (*b* = –0.61, *SE* = 0.72, 95% CI [–2.03, 0.80], *t*= -0.853, *p=* 0.3940).
Post hoc contrasts confirmed that the HR increase in the Congruent condition was significantly greater than in both No Feedback (*estimate* = 3.51, *SE* = 0.89, *t*(1320) = 3.95, *p* = .0003) and Slower feedback (*estimate* = 2.24, *SE* = 0.80, *t*(1307) = 2.79, *p* = .0103), but did not differ significantly from Faster feedback (*estimate* = 0.23, *SE* = 0.82, *t*(1313) = 0.28, *p* = 0.78).

In the *Exteroceptive experiment*, HR showed a non-significant decreasing trend over trials in the Congruent condition (*b* = –1.13, *SE* = 0.68, 95% CI [–2.46, 0.20]), and remained flat in the Slower condition (*b* = –0.001, *SE* = 0.65, 95% CI [–1.29, 1.29]). Similarly, HR did not significantly change over time in the Faster condition (*b* = –0.82, *SE* = 0.74, 95% CI [–2.28, 0.64]) nor in the No Feedback condition (*b* = 1.28, *SE* = 0.81, 95% CI [–0.30, 2.87]).
However, post hoc contrasts showed that the No Feedback condition was associated with a significantly different HR trajectory compared to Congruent feedback (*estimate* = –2.42, *SE* = 0.91, *t*(1310) = –2.65, *p* = 0.049), while no other significant differences emerged between Congruent and Slower (*p* = 0.227) or Faster feedback (*p* = 0.714).

Finally, interaction contrasts on the HR slope revealed several significant differences in feedback-related trajectories across the two experimental contexts.

The difference in HR slope between Congruent and No Feedback conditions was significantly larger in the Interoceptive experiment than in the Exteroceptive one (*b* = 5.92, *SE* = 1.27, *t* = 4.65, *p* < .0001). Specifically, HR increased over time in the Congruent condition but remained flat or even decreased in the No Feedback condition, and this contrast was markedly stronger in the Interoceptive context.

A similar interaction was found for the comparison between Faster and No Feedback (*b* = 5.38, *SE* = 1.27, *t* = 4.23, *p* < .0001), indicating that the presence of temporally accelerated feedback boosted HR increase significantly more in the Interoceptive experiment than in the Exteroceptive one.

Additionally, the contrast between Slower and No Feedback was also significantly greater in the Interoceptive context (*b* = 2.56, *SE* = 1.21, *t* = 2.12, *p* = .0345). Beyond these comparisons with No Feedback, additional interaction effects were observed among active feedback conditions.

The contrast between Slower and Faster feedback differed significantly across experiments (*b* = -2.82, *SE* = 1.12, *t* = -2.52, *p* = .0117), reflecting a stronger HR increase under Faster than under Slower feedback in the Interoceptive experiment, a pattern not observed in the Exteroceptive experiment.

The contrast between Congruent and Slower was also significantly greater in Interoceptive compared to Exteroceptive experiment (*b* = 3.37, *SE* = 1.12, *t* = 3.01, *p* = .0027), while no significant difference emerged between Congruent and Faster feedback conditions across experiments (*b* = 0.54, *SE* = 1.19, *t* = 0.46, *p* = .647).

##### 1.2 Likert Pain Unpleasantness Ratings Scale

To examine variations in pain unpleasantness ratings across experimental conditions, we applied general linear mixed-effects modeling (GLMM), which accounts for both between- and within-participant variability. The model was fitted using the lmer function from the *lme4* package (Bates et al., 2015) in R and specified as follows:

LIKERT PAIN UNPLEASANTNESS RATINGS ~ Experiment × Feedback × StimInt × Trial + (StimInt + Trial | Subject)

Experiment, Feedback, StimInt and Trial were included as fixed effects. Random slopes for Stimulus Intensity and Trial were specified at the subject level to account for individual variability. Model comparison confirmed that including these random slopes significantly improved model fit (χ² = 117.73, p < 0.001).

###### 1.2.1 Results

###### 1.2.1.1 Effects of interest

The Type III ANOVA revealed a significant **main effect of** **Feedback**, F(3, 1228.10) = 40.30, p < .001, indicating that unpleasantness ratings differed across feedback conditions. Compared to the Congruent condition (reference), unpleasantness ratings were significantly higher in the Faster feedback condition (b = 0.14, SE = 0.048, t(1231.07) = 2.87, p = .004), and significantly lower in the No Feedback condition (b = –0.23, SE = 0.047, t(1228.15) = –4.97, p < .0001). The Slower condition did not differ significantly from the Congruent condition (p = .17). Post hoc comparisons further clarified these effects. Likert pain unpleasantness ratings were significantly higher in the Faster feedback condition compared to Congruent feedback (p= 0.0096), Slower feedback (p= 0.0006) and No Feedback (p<0.001). The Slower feedback condition did not differ significantly from the Congruent condition (p=0.415). Moreover, the No Feedback yielded a reduction in Numeric Pain Unpleasantness ratings compared to all other feedback conditions (all p<0.0001).

Although the **Feedback × Experiment interaction** did not reach conventional significance in the omnibus ANOVA (F(3, 1228.10) = 2.32, p = .074), the results suggested a trend indicating that the effect of feedback on pain unpleasantness may differ between the Interoceptive and Exteroceptive experiments. Given that this was one of the main hypotheses of the study, we conducted follow-up analyses to further this interaction.

In the *Interoceptive experiment*, unpleasantness ratings in the No Feedback condition (M = 2.80, SE = 0.105) were significantly lower than in all other feedback conditions. Specifically, ratings were significantly lower than in the Faster condition (M = 3.17, SE = 0.105; estimate = –0.37, SE = 0.046, t(1232) = –7.97, p < .0001), the Slower condition (M = 2.97, SE = 0.105; estimate = –0.17, SE = 0.046, t(1232) = –3.66, p = .0004), and the Congruent condition (M = 3.03, SE = 0.105; estimate = –0.23, SE = 0.047, t(1229) = –4.95, p < .0001). Importantly, unpleasantness ratings in the Faster condition were significantly higher than in the Slower conditions (p < .0001), as well as in the Congruent (p = .0051) and, while no significant difference emerged between the Congruent and Slower conditions (p = .17).

In the *Exteroceptive experiment*, unpleasantness ratings in the No Feedback condition (M = 2.75, SE = 0.113) were again significantly lower than in all other feedback conditions. Ratings in the No Feedback condition were significantly lower than in the Faster (M = 3.10, SE = 0.113; estimate = –0.34, SE = 0.049, p < .0001), Slower (M = 3.06, SE = 0.113; estimate = –0.30, SE = 0.049, p < .0001), and Congruent conditions (M = 3.05, SE = 0.113; estimate = –0.29, SE = 0.050, p < .0001). In contrast to the Interoceptive experiment, no significant differences emerged between the Congruent, Slower, and Faster feedback conditions (all ps > .52).

To directly compare the feedback-related differences across experiments, interaction contrasts were computed.

Importantly, the contrast between Slower and Faster feedback was significantly more pronounced in the Interoceptive than in the Exteroceptive experiment (b = –0.16, SE = 0.067, t = –2.43, p = .015), indicating a greater differentiation in unpleasantness between these conditions when the feedback was interoceptive. Similarly, the contrast between Slower and No Feedback was also significantly larger in the Interoceptive context (b = –0.14, SE = 0.067, t = –2.01, p = .0445). No other feedback comparisons showed significant interaction effects between experiments (all ps > .18).

###### 1.2.1.2 Additional unpredicted effects

A Type III ANOVA revealed a significant **main effect of Trial** on pain unpleasantness ratings, F(1, 64.50) = 19.63, p < .001, suggesting that unpleasantness ratings varied throughout the experimental session. However, the fixed-effect estimate of the Trial coefficient did not reach statistical significance (b = 0.27, SE = 0.15, t(269.04) = 1.76, p = .079), indicating a non-significant trend toward increasing unpleasantness ratings over time. This apparent discrepancy reflects the fact that Trial was involved in several higher-order interactions, particularly with Stimulus Intensity and Feedback, that modulated its effect on unpleasantness ratings. These interactions are reported in detail below.

There was also a significant **main effect of Stimulus Intensity (StimInt)** on unpleasantness ratings, F(1, 62.30) = 352.15, p < .001. Specifically, unpleasantness ratings increased significantly with higher stimulation levels, as confirmed by a strong and significant positive coefficient (b = 1.52, SE = 0.12, t(172.73) = 12.82, p < .0001), indicating that higher stimulus intensity reliably evoked greater unpleasantness.

Additionally, the effect of trial on pain unpleasantness ratings varied as a function of the actual intensity of the nociceptive stimulation, as indicated by a significant **Stimulus Intensity × Trial** interaction, F(1, 1253.50) = 17.84, p < .001. To further investigate this interaction, estimated marginal trends of unpleasantness ratings across trials were computed at three levels of stimulus intensity: low (–0.5), medium (0), and high (+0.5). Unpleasantness ratings increased progressively over trials at all intensity levels, but the rate of increase became steeper as stimulus intensity increased.

At low intensity, the slope was positive but did not reach statistical significance (b = 0.12, SE = 0.09, 95% CI [–0.06, 0.30] , t(123.4) = 1.32, p = .191). A significant positive trend emerged at medium intensity (b = 0.34, SE = 0.08, 95% CI [0.19, 0.49], t(64.6) = 4.43, p < .0001) and was even more pronounced at high intensity (b = 0.56, SE = 0.09, 95% CI [0.37, 0.74] , t(147.4) = 5.90, p < .0001).

Pairwise comparisons between slopes (FDR-corrected) confirmed that the increase in unpleasantness ratings over trials was significantly steeper at higher stimulus intensity levels. Specifically, the slope at low intensity was significantly lower than both the medium (estimate = –0.22, SE = 0.052, t(1254) = –4.22, p < .0001) and high intensity levels (estimate = –0.44, SE = 0.10, t(1254) = –4.22, p < .0001). Moreover, the slope at medium intensity was also significantly lower than at high intensity (estimate = –0.22, SE = 0.052, t(1254) = –4.22, p < .0001).

The **Feedback × Stimulus Intensity** interaction was also significant, F(3, 1237.96) = 10.50, p < .001, indicating that the effect of stimulus intensity on pain unpleasantness ratings varied across feedback conditions. To further this interaction, estimated marginal means (EMMs) of unpleasantness ratings were computed at the three levels of stimulus intensity: low (–0.5), medium (0), and high (+0.5).

At *low stimulus intensity* (–0.5), the Faster feedback condition elicited the highest unpleasantness ratings (M = 2.49, SE = 0.08), followed closely by Slower (M = 2.47, SE = 0.08), Congruent (M = 2.28, SE = 0.08), and No Feedback (M = 2.20, SE = 0.08). Ratings in the Faster condition were significantly higher than those in the Congruent (estimate = –0.22, SE = 0.06, t(1238) = –3.92, p = .0002) and No Feedback condition (estimate = 0.29, SE = 0.05, t(1235) = 5.39, p < .0001). Likewise, Slower feedback elicited significantly higher ratings than both Congruent (estimate = –0.19, SE = 0.05, t(1236) = –3.50, p = .0007) and No Feedback (estimate = 0.26, SE = 0.05, t(1233) = 4.98, p < .0001). However, the difference between Congruent and No Feedback did not reach significance (p = .2213)

At *medium intensity* (0), the Faster feedback produced the highest unpleasantness ratings (M = 3.13, SE = 0.08), followed by Congruent (M = 3.04, SE = 0.08), Slower (M = 3.01, SE = 0.08), and No Feedback (M = 2.78, SE = 0.08). Post hoc comparisons revealed that ratings were significantly higher in the Faster condition compared to both Congruent (estimate = –0.09, SE = 0.03, t(1229) = –2.66, p = .0096) and Slower (estimate = –0.12, SE = 0.03, t(1228) = –3.53, p = .0006). All three active feedback conditions also elicited significantly higher ratings than No Feedback (all ps < .0001).

At *high intensity* (+0.5), unpleasantness ratings were highest under the Congruent feedback (M = 3.80, SE = 0.10), followed by the Faster (M = 3.77, SE = 0.10), Slower (M = 3.56, SE = 0.10), and No Feedback (M = 3.35, SE = 0.10). Ratings in the No Feedback condition were significantly lower than those in all other conditions (all ps < .0001), and ratings in the Faster condition were significantly higher than Slower (estimate = –0.21, SE = 0.05, t(1232) = –3.99, p = .0001). Congruent also differed significantly from Slower (estimate = 0.25, SE = 0.05, t(1232) = 4.54, p < .0001), whereas no significant difference emerged between Congruent and Faster (p = .5377).

Unpleasantness ratings increased significantly as a function of stimulus intensity in all feedback conditions. However, the rate of increase (slope) varied depending on feedback type. The steepest increase was observed in the Congruent condition (b = 1.53, SE = 0.09, t(164) = 17.76, p < .0001), followed by Faster (b = 1.28, SE = 0.08, t(154) = 15.10, p < .0001), No Feedback (b = 1.14, SE = 0.08, t(148) = 13.68, p < .0001), and Slower (b = 1.09, SE = 0.08, t(149) = 12.99, p < .0001). Pairwise comparisons between slopes revealed that the slope was significantly steeper in the Congruent condition compared to Slower (estimate = 0.44, SE = 0.08, t(1239) = 5.17, p < .0001), Faster (estimate = 0.25, SE = 0.09, t(1239) = 2.93, p = .0069), and No Feedback (estimate = 0.38, SE = 0.08, t(1237) = 4.52, p < .0001). The difference between Faster and Slower also reached significance (estimate = 0.19, SE = 0.08, t(1239) = 2.25, p = .0367), whereas other comparisons did not (all ps > .13).

The interaction between **Feedback × Trial** was significant, F(3, 1239.19) = 14.39, p < .0001, indicating that the trajectory of unpleasantness ratings across trials varied depending on the type of feedback received.

To further unpack this interaction, the estimated slopes of Trial were computed for each feedback condition. Unpleasantness ratings increased significantly across trials in both the Congruent (b = 0.24, SE = 0.11, 95% CI [0.03, 0.45] , t(235) = 2.22, p = .0272) and Slower feedback conditions (b = 0.29, SE = 0.10, 95% CI [0.10, 0.48] , t(168) = 2.96, p = .0036), indicating a moderate but consistent increase in perceived unpleasantness over time. In contrast, no significant change was observed in the Faster condition (b = 0.01, SE = 0.10, 95% CI [–0.20, 0.22] , t(221) = 0.08, p = .9339). The steepest and most robust increase in unpleasantness ratings over time was found in the No Feedback condition (b = 0.82, SE = 0.11, 95% CI [0.59, 1.04] , t(286) = 7.22, p < .0001).

Post hoc pairwise comparisons of slopes (FDR-corrected) confirmed that the rate of increase in unpleasantness over trials was significantly greater in the No Feedback condition compared to all other feedback types (all ps < .0001). Specifically, the slope in the No Feedback condition was significantly greater than in the Congruent (estimate = –0.58, SE = 0.13, t(1245) = –4.56, p < .0001), Slower (estimate = –0.53, SE = 0.12, t(1240) = –4.45, p < .0001), and Faster conditions (estimate = –0.81, SE = 0.13, t(1242) = –6.44, p < .0001). Additionally, the slope in the Faster condition was significantly lower than both the Congruent (p = .0682, marginal) and Slower conditions (estimate = –0.28, SE = 0.11, t(1236) = –2.50, p = .0186), while no significant difference emerged between the Congruent and Slower conditions (p = .6562).

To further explore how the effect of feedback on unpleasantness evolved over time, estimated marginal means were computed at three levels of the standardized Trial variable (–0.5, 0, +0.5), representing early, middle, and late stages of the session, respectively. This analysis allowed us to assess whether the differences between feedback conditions were stable or changed over time.

At *early trials (*Trial = –0.5), unpleasantness ratings were significantly higher in the Faster condition (M = 3.13, SE = 0.09) compared to both the Congruent (M = 2.92, SE = 0.09; estimate = –0.21, SE = 0.07, t(1234) = –3.10, p = .0024) and Slower conditions (M = 2.87, SE = 0.09; estimate = –0.26, SE = 0.07, t(1231) = –3.81, p = .0002). The No Feedback condition elicited the lowest ratings (M = 2.37, SE = 0.10), significantly lower than all other feedback conditions (all ps < .0001).

At *middle trials* (Trial = 0), the pattern remained consistent: Faster feedback yielded significantly higher ratings (M = 3.13, SE = 0.08) compared to Congruent (M = 3.04, SE = 0.08; estimate = –0.09, SE = 0.03, t(1229) = –2.66, p = .0094) and Slower (M = 3.01, SE = 0.08; estimate = –0.12, SE = 0.03, t(1227) = –3.55, p = .0006), while the No Feedback condition (M = 2.77, SE = 0.08) was again significantly lower than all other conditions (all ps < .0001).

By *late trials* (Trial = 0.5), feedback-related differences in unpleasantness ratings had disappeared. No significant differences were found among any of the feedback conditions (all ps > .90), and ratings converged: Congruent (M = 3.16, SE = 0.10), Slower (M = 3.15, SE = 0.09), Faster (M = 3.14, SE = 0.09), and No Feedback (M = 3.18, SE = 0.10). This pattern suggests that, as the session progressed, the modulatory effect of feedback on unpleasantness diminished.

##### 1.3 Numeric Pain Scale of intensity ratings

To examine variations in pain intensity ratings (Numeric pain scale of intensity ratings) across experimental conditions, we applied general linear mixed-effects modeling (GLMM), which accounts for both between- and within-participant variability by estimating fixed effects while allowing subject-specific random variations. The model was fitted using R (lme4 package; Bates et al., 2015) and was specified as follows:

NUMERIC PAIN SCALE OF INTENSITY RATINGS~ Experiment × Feedback × StimInt × Trial + (StimInt + Trial | Subject)

where Experiment, Feedback. StimInt, and Trial were included as fixed effects. A random slope for StimInt and Trial was included at the subject level (StimInt + Trial | Subject) to account for individual differences in pain perception across stimulus intensities and over time. Model comparison confirmed that including these random slopes significantly improved model fit (χ² = 134.14, p < 0.001). The statistical significance of predictors was assessed using t-tests with Satterthwaite’s approximation to degrees of freedom. For each predictor, we report the estimated coefficient (b), standard error (SE), t-statistic (t), and p-value (p).

###### 1.3.1 Results

###### 1.3.1.1 Effects of interest

The results of the Type III ANOVA with Satterthwaite’s approximation revealed a significant **main effect of Feedback,** F(3,1228.44) = 54.41, p < 0.001, indicating that pain intensity ratings varied across feedback conditions. Compared to the reference condition (i.e., Congruent feedback), pain ratings were significantly lower in the No Feedback condition (b = -4.67, SE = 1.08, t = -4.31, p < 0.001) and significantly higher in the Faster feedback condition (b = 3.88, SE = 1.10, t = 3.53, p < 0.001). The Slower feedback condition showed no significant difference compared to the Congruent (p = 0.53). Post hoc comparisons further clarified these effects. Numeric pain scale of intensity ratings were significantly higher in the Faster feedback condition compared to Congruent feedback (p=0.0002), Slower feedback (p= 0.0002) and No Feedback (p<0.001). The Slower feedback condition did not differ significantly from the Congruent condition (p=0.8722). Moreover, the No Feedback yielded a reduction in Numeric pain scale of intensity ratings compared to all other feedback conditions (all p<0.0001).

Crucially, the **Feedback × Experiment interaction** was also significant, F(3, 1228.44) = 4.48, p = 0.004, indicating that the effect of Feedback on pain ratings differed between the Interoceptive and Exteroceptive experiments.

In the *Interoceptive experiment*, pain ratings in the No Feedback condition (*M* = 51.1, *SE* = 2.65) were significantly lower than in all other feedback conditions. Specifically, pain ratings were significantly lower than in the Faster condition (*M* = 59.6, *SE* = 2.66; *estimate* = –8.55, *SE* = 1.07, *t*(1234) = –7.99, *p* < .0001), the Slower condition (*M* = 55.1, *SE* = 2.65; *estimate* = –3.99, *SE* = 1.05, *t*(1231) = –3.79, *p* = .0002), and the Congruent condition (*M* = 55.8, *SE* = 2.66; *estimate* = –4.70, *SE* = 1.08, *t*(1230) = –4.34, *p* < .0001). Importantly, pain ratings in the Faster condition were significantly higher than in the Slower condition (*estimate* = –4.56, *SE* = 1.07, *t*(1233) = –4.26, *p* < .0001), and in the Congruent condition (*estimate* = –3.86, *SE* = 1.10, *t*(1234) = –3.51, *p* = .0006). No significant difference was observed between the Slower and Congruent feedback conditions (*estimate* = 0.71, *SE* = 1.08, *t*(1230) = 0.65, *p* = .515).

In the *Exteroceptive experiment*, pain ratings in the No Feedback condition (*M* = 51.0, *SE* = 2.87) were also significantly lower than in all other feedback conditions. Specifically, pain ratings were lower than in the Congruent condition (*M* = 59.5, *SE* = 2.87; *estimate* = –8.48, *SE* = 1.15, *t*(1231) = –7.35, *p* < .0001), the Slower condition (*M* = 60.5, *SE* = 2.87; *estimate* = –9.44, *SE* = 1.14, *t*(1230) = –8.24, *p* < .0001), and the Faster condition (*M* = 61.8, *SE* = 2.87; *estimate* = –10.73, *SE* = 1.14, *t*(1230) = –9.41, *p* < .0001). In contrast to the Interoceptive experiment, no significant differences emerged between the Congruent, Slower, and Faster feedback conditions (Congruent vs. Slower: *estimate* = –0.96, *SE* = 1.15, *t*(1229) = –0.83, *p* = .406; Congruent vs. Faster: *estimate* = –2.25, *SE* = 1.15, *t*(1229) = –1.96, *p* = .075; Slower vs. Faster: *estimate* = –1.29, *SE* = 1.14, *t*(1230) = –1.13, *p* = .309).

Importantly, interaction contrasts revealed a significant difference in the effect of Slower vs. Faster feedback across experiments (b = -3.27, SE = 1.56, t = -2.09, p = 0.0368). This effect was strongest in the Interoceptive experiment, where Faster feedback led to a markedly greater increase in pain intensity. The effect of Slower vs. No feedback differed significantly between experiments, with a stronger difference in the Interoceptive experiment (b = -5.45, SE = 1.56, t = -3.501, p = 0.0005). Likewise, the Congruent vs. No feedback contrast was more pronounced in the Interoceptive experiment (b = -3.78, SE = 1.58, t = -2.390, p = 0.0170). No other between-experiment differences were significant.

###### 1.3.1.2 Additional unpredicted effects

A Type III ANOVA revealed a significant **main effect of Trial** on Numeric pain scale of intensity ratings, F(1, 61.94) = 17.56, p < .001, suggesting that pain ratings varied throughout the experimental session. However, the fixed-effect estimate of the Trial coefficient did not reach statistical significance (b = 5.30, SE = 3.67, t(210.37) = 1.45, p = .15), indicating a non-significant trend toward increasing intensity ratings over time. These interactions are reported in detail below.

There was also a significant main effect of **Stimulus Intensity (StimInt)** on pain ratings, F(1, 62.32) = 386.62, p < 0.001. Specifically, pain ratings increased significantly as the intensity of the stimulus increased, with a coefficient estimate of 37.54 (SE = 2.90), t = 12.94, p < 0.001. This indicates a strong and significant positive relationship between stimulus intensity and reported pain levels.

Additionally, the effect of trial on pain ratings varied as a function of the actual intensity of the nociceptive stimulation, as indicated by a significant **Stimulus Intensity × Trial** **interaction**, *F*(1, 1249.76) = 14.07, *p* < .001.

To further investigate this interaction, estimated marginal trends of the Numeric pain scale of intensity ratings across trials were computed at three levels of stimulus intensity: low (-0.5), medium (0), and high (+0.5). Pain ratings increased progressively over trials at all intensity levels, but the rate of increase became steeper as stimulus intensity increased. At low intensity, the slope was positive but only marginally significant (*b* = 3.77, *SE* = 2.26, 95% CI [–0.72, 8.25], *t*(109.3) = 1.67, *p* = .099). A significant positive trend emerged at medium intensity (*b* = 8.26, *SE* = 1.97, 95% CI [4.32, 12.20] , *t*(63.8) = 4.19, *p* = .0001) and was even more pronounced at high intensity (*b* = 12.75, *SE* = 2.35, 95% CI [8.10, 17.40], *t*(126.6) = 5.43, *p* < .0001).

Pairwise comparisons between slopes (FDR-corrected) confirmed that the increase in Numeric pain scale of intensity ratings over trials was significantly steeper at higher levels of stimulus intensity. Specifically, the slope at low intensity was significantly lower than both the medium (*estimate* = –4.49, *SE* = 1.20, *t*(1251) = –3.75, *p* = .0002) and high intensity levels (*estimate* = –8.99, *SE* = 2.40, *t*(1251) = –3.75, *p* = .0002). Moreover, the slope at medium intensity was significantly lower than at high intensity (*estimate* = –4.49, *SE* = 1.20, *t*(1251) = –3.75, *p* = .0002).

The **Feedback × Stimulus Intensity** **interaction** was also significant, *F*(3, 1237.89) = 5.27, *p* = .001, indicating that the effect of stimulus intensity on pain ratings varied across feedback conditions. To further unpack this interaction, estimated marginal means (EMMs) of Numeric pain scale of intensity ratings were computed at the three levels of stimulus intensity: low (–0.5), medium (0), and high (+0.5).

At *low stimulus intensity* (–0.5), the Faster feedback condition elicited the highest pain ratings (M = 43.5, SE = 2.09), followed by Slower (M = 42.5, SE = 2.09), Congruent (M = 39.0, SE = 2.11), and No Feedback (M = 34.6, SE = 2.09). Ratings in the Faster condition were significantly higher than those in the Congruent condition (estimate = –4.42, SE = 1.50, t(1309) = –2.94, p = .0050), and also significantly higher than those in the No Feedback condition (estimate = 8.85, SE = 1.48, t(1308) = 5.99, p < .0001). Likewise, the Slower condition elicited significantly higher ratings than Congruent, t(1306) = -2.284, p = .0271, and No Feedback, t(1307) = 5.346, p < 0.0001. Ratings in the No Feedback condition were significantly lower than in all other feedback conditions (all ps < .0006).

At *medium intensity* (0), Faster again produced the highest ratings (M = 60.5, SE = 1.93), followed by Slower (M = 57.3, SE = 1.92), Congruent (M = 57.3, SE = 1.93), and No Feedback (M = 50.5, SE = 1.92). Post hoc comparisons confirmed that Faster was significantly higher than both Congruent (estimate = –3.25, SE = 0.95, t(1301) = –3.44, p = .0007) and Slower (estimate = –3.20, SE = 0.93, t(1300) = –3.45, p = .0007), while all three active feedback conditions yielded significantly higher ratings than No Feedback (all ps < .0001).

At *high intensity* (+0.5), Faster again elicited the highest ratings (M = 77.6, SE = 2.32), followed by Congruent (M = 75.6, SE = 2.34), Slower (M = 72.2, SE = 2.31), and No Feedback (M = 66.3, SE = 2.31). The No Feedback condition was again significantly lower than all others (all ps < .0002), and Faster was significantly higher than Slower (estimate = –5.40, SE = 1.47, t(1306) = –3.67, p = .0004). A significant difference was also observed between Congruent and Slower (estimate = 3.33, SE = 1.49, t(1307) = 2.23, p = .0311), while the difference between Congruent and Faster was not significant (p = .1706).

Numeric pain scale of intensity ratings increased as a function of stimulus intensity in all feedback conditions. However, the rate of increase (slope) differed depending on the feedback type. The steepest increase was observed in the Congruent condition (b = 36.5, SE = 2.20, t(197) = 16.59, p < .0001), followed by Faster (b = 34.2, SE = 2.17, t(186) = 15.77, p < .0001), No Feedback (b = 31.7, SE = 2.15, t(178) = 14.75, p < .0001), and Slower (b = 29.8, SE = 2.15, t(178) = 13.86, p < .0001). Pairwise comparisons between slopes revealed that the increase in pain ratings was significantly steeper in the Congruent condition compared to Slower (estimate = 6.74, SE = 2.33, t(1312) = 2.90, p = .0229). Differences between other slopes did not reach statistical significance (all ps > .11).

The interaction between **Feedback x Trial** was significant, *F*(3, 1238.72) = 20.77, *p* < .0001, indicating that the trajectory of pain ratings across trials varied depending on the type of feedback received.

To further unpack this interaction, the estimated slopes of Trial were computed for each feedback condition. Pain ratings increased significantly across trials in both the Congruent (*b* = 6.58, *SE* = 2.61, 95% CI [1.42, 11.73] , *t*(190) = 2.52, *p* = .0127) and Slower feedback conditions (*b* = 6.85, *SE* = 2.42, 95% CI [2.06, 11.63] , *t*(142) = 2.83, *p* = .0053), indicating a moderate but consistent increase in perceived pain over time. In contrast, no significant change across trials was observed in the Faster condition (*b* = –1.51, *SE* = 2.58, 95% CI [–6.59, 3.57] , *t*(181) = –0.59, *p* = .558). The steepest and most robust increase in pain ratings over time was found in the No Feedback condition (*b* = 21.27, *SE* = 2.77, 95% CI [15.80, 26.73] , *t*(235) = 7.67, *p* < .0001).

Post hoc pairwise comparisons of slopes (FDR-corrected) confirmed that the rate of pain increase over trials was significantly greater in the No Feedback condition compared to all other feedback types (all *p*s < .0001). Specifically, the slope in the No Feedback condition was significantly higher than that of the Congruent (*estimate* = –14.69, *SE* = 2.94, *t*(1244) = –4.99, *p* < .0001), Slower (*estimate* = –14.42, *SE* = 2.76, *t*(1241) = –5.22, *p* < .0001), and Faster conditions (*estimate* = –22.78, *SE* = 2.91, *t*(1241) = –7.83, *p* < .0001). Additionally, the slope in the Faster condition was significantly lower than that of both the Congruent (*estimate* = 8.090, *SE* = 2.76 , *t*(1240) = 2.931, *p* = .0041) and Slower conditions (*estimate* = 8.359, *SE* = 2.57, *t*(1236) = 3.251, *p* = . 0018), while no significant difference emerged between the Congruent and Slower conditions (*p* = .918).

To further explore how the effect of feedback on pain ratings evolved over time, estimated marginal means of Numeric pain scale of intensity ratings were computed at three levels of the standardized Trial variable (–0.5, 0, +0.5), representing earlier, middle, and later stages of the experimental session, respectively. This analysis allowed us to probe whether the differences in pain ratings across feedback conditions were stable or varied as the experiment progressed.

At *early trials* (Trial = –0.5), pain ratings differed markedly across feedback conditions. Ratings were significantly higher in the Faster condition (M = 61.5, SE = 2.35) compared to both the Congruent (M = 54.4, SE = 2.27; estimate = –7.10, SE = 1.54, t(1235) = –4.62, p < .0001) and Slower conditions (M = 54.4, SE = 2.29; estimate = –7.11, SE = 1.57, t(1233) = –4.53, p < .0001). In contrast, the No Feedback condition elicited the lowest pain ratings (M = 40.4, SE = 2.40), significantly lower than all other conditions (all ps < .0001), with the largest difference observed when compared to the Faster condition (estimate = –21.03, SE = 1.72, t(1239) = –12.23, p < .0001).

At *middle trials* (Trial = 0), the pattern of feedback differences remained largely consistent. Pain ratings were again significantly higher in the Faster condition (M = 60.7, SE = 1.95) than in the Congruent (M = 57.7, SE = 1.96; estimate = –3.06, SE = 0.80, t(1231) = –3.84, p = .0002) and Slower conditions (M = 57.8, SE = 1.95; estimate = –2.93, SE = 0.78, t(1231) = –3.75, p = .0002). The No Feedback condition continued to yield significantly lower ratings (M = 51.1, SE = 1.95) compared to all other feedback conditions (all ps < .0001).

At *later trials* (Trial = 0.5), the feedback-related differences in pain ratings disappeared. No significant differences emerged among any of the feedback conditions (all ps > .87), and pain ratings converged across all feedback types: Congruent (M = 60.9, SE = 2.44), Slower (M = 61.2, SE = 2.30), Faster (M = 60.0, SE = 2.33), and No Feedback (M = 61.7, SE = 2.39). This pattern suggests that while feedback modulated pain ratings in the earlier and middle stages of the experiment, these effects dissipated over time, resulting in equivalent pain ratings by the end of the session.

#### 2. Interoceptive experiment (within-subjects model)

##### 2.1 Heart rate

To examine variations in heart rate (HR) within the interoceptive experiment, we applied general linear mixed-effects modeling (GLMM), the model was fitted using R (lme4 package; Bates et al., 2015; Bates et al., 2015) and was specified as follows:

𝐻𝑅∼𝐹𝑒𝑒𝑑𝑏𝑎𝑐𝑘×𝑇𝑟𝑖𝑎𝑙+(𝑇𝑟𝑖𝑎𝑙∣𝑆𝑢𝑏𝑗𝑒𝑐𝑡)

where Feedback and Trial were included as fixed effects, and a random slope for Trial at the subject level (Trial | Subject) to account for individual variability in HR trajectories over time. Model comparison indicated that the inclusion of a random slope significantly improved model fit (χ² = 13.493, p = .001).

###### 2.1.1 Results

###### 2.1.1.1 Effects of interest

The Type III ANOVA revealed a significant **main effect of Feedback**, F(3, 707.58) = 60.70, p < .0001, indicating that HR levels varied depending on the type of feedback received. Compared to the Congruent condition (M = 74.2, SE = 1.37), HR was significantly lower in the Faster feedback condition (M = 73.1, SE = 1.37; estimate = -1.05, SE = 0.25, t(708) = - 4.28, p < .0001), and significantly higher in the No Feedback condition (M = 76.2, SE = 1.37; estimate = 2.03, SE = 0.24, t(707) = 8.32, p < .0001). In contrast, the Slower feedback condition did not significantly differ from the Congruent (M = 73.8, SE = 1.37, estimate = -0.37, SE = 0.24, t(705.6) = –1.53, p = .1277). *Post hoc comparisons* further clarified these effects. Relative to all feedback conditions, the No feedback elicited a significantly increased heart rate (all ps<0.0001). Additionally, heart rate decreased significantly under Faster feedback relative to the Slower (estimate = 0.694, SE = 0.240, t(708) = 2.898, p = .0046) and Congruent feedback (estimate = 1.058, SE = 0.247, t(708) = 4.284, p < .0001). The Slower feedback condition did not differ significantly from the Congruent condition (*p*=0.14).

###### 2.1.1.2 Additional unpredicted effects

A significant **main effect of Trial** also emerged, F(1, 34.95) = 8.23, p = .0069, indicating that HR increased throughout the experiment (b = 2.66, SE = 0.74, t(164.7) = 3.58, p = .0004).

The interaction between **Feedback × Trial** was significant, F(3, 711.19) = 6.16, p < .001, indicating that the trajectory of heart rate (HR) across trials varied depending on the type of feedback received (See Supplementary Figure 1).

To further unpack this interaction, the estimated slopes of Trial were computed for each feedback condition. HR increased significantly across trials in both the Congruent (b = 2.67, SE = 0.74, 95% CI [1.20, 4.13], t(164) = 3.58, p = .0005) and Faster feedback conditions (b = 2.66, SE = 0.68, 95% CI [1.31, 4.00], t(119) = 3.91, p = .0002), indicating a consistent rise in autonomic arousal over time. In contrast, the Slower condition did not show a significant change across trials (b = 0.68, SE = 0.65, 95% CI [–0.61, 1.98], t(104) = 1.04, p = .2993), while the No Feedback condition showed a flat-to-decreasing pattern that was also non-significant (b = –0.34, SE = 0.75, 95% CI [–1.82, 1.15], t(169) = –0.44, p = .6574).

Post hoc pairwise comparisons of slopes (FDR-corrected) confirmed that the rate of HR increase over trials was significantly higher in the Congruent condition compared to both Slower (estimate = 1.98, SE = 0.81, t(707) = 2.44, p = .0227) and No Feedback conditions (estimate = 3.00, SE = 0.90, t(715) = 3.34, p = .0026). The slope in the Faster condition was also significantly greater than that of Slower (estimate = 1.98, SE = 0.76, t(707) = 2.61, p = .0188) and No Feedback (estimate = 2.99, SE = 0.85, t(714) = 3.54, p = .0026). No significant difference emerged between Congruent and Faster feedback (p = .9928), nor between Slower and No Feedback (p = .2611).

To further explore how the effect of feedback on heart rate (HR) evolved over time, estimated marginal means of HR were computed at three levels of the standardized Trial variable (–0.5, 0, +0.5), representing early, middle, and late stages of the experimental session. This analysis allowed us to assess whether differences in HR across feedback conditions were stable or changed as the session progressed.

At *early trials* (Trial = –0.5), HR was significantly higher in the No Feedback condition (M = 76.4, SE = 1.33) compared to all other feedback conditions. Specifically, HR was significantly higher than in the Faster (M = 71.8, SE = 1.33; estimate = –4.57, SE = 0.52, t(713) = –8.86, p < .0001), Slower (M = 73.5, SE = 1.32; estimate = –2.90, SE = 0.48, t(713) = –6.08, p < .0001), and Congruent conditions (M = 72.8, SE = 1.32; estimate = –3.52, SE = 0.47, t(715) = –7.43, p < .0001). Additionally, the Faster condition elicited significantly lower HR than both the Slower (estimate = 1.68, SE = 0.48, t(705) = 3.52, p = .0007) and Congruent conditions (estimate = 1.05, SE = 0.47, t(707) = 2.23, p = .0315), while no significant difference emerged between Congruent and Slower feedback (p = .1472).

At *middle trials* (Trial = 0), the same pattern of differences was observed. HR in the No Feedback condition (M = 76.2, SE = 1.37) remained significantly higher than in all other conditions (all ps < .0001). The Faster condition (M = 73.1, SE = 1.37) was again significantly lower than Slower (M = 73.8, SE = 1.37; estimate = –0.69, SE = 0.24, t(708) = –2.89, p = .0048) and Congruent (M = 74.2, SE = 1.37; estimate = –1.06, SE = 0.25, t(708) = –4.28, p < .0001). Congruent and Slower did not differ significantly (p = .1277).

By *later trials* (Trial = 0.5), HR values across conditions began to converge. Although No Feedback (M = 76.0, SE = 1.50) was still numerically higher, it no longer differed significantly from Congruent (M = 75.5, SE = 1.51; estimate = –0.52, SE = 0.55, t(712) = –0.95, p = .4092). However, Faster (M = 74.4, SE = 1.48) and Slower (M = 74.1, SE = 1.49) still showed significantly lower HR compared to No Feedback (Faster vs. No Feedback: estimate = –1.58, SE = 0.46, t(712) = –3.46, p = .0017; Slower vs. No Feedback: estimate = –1.88, SE = 0.47, t(711) = –3.97, p = .0005). The Congruent condition was significantly higher than Slower (estimate = 1.36, SE = 0.51, t(706) = 2.65, p = .0163) and marginally higher than Faster (estimate = 1.06, SE = 0.50, t(712) = 2.13, p = .0507).

Taken together, these results indicate that the Faster feedback condition consistently elicited lower heart rate (HR) compared to the other feedback types, particularly during the early and middle phases of the experimental session, and this effect tended to vanish over later trials.

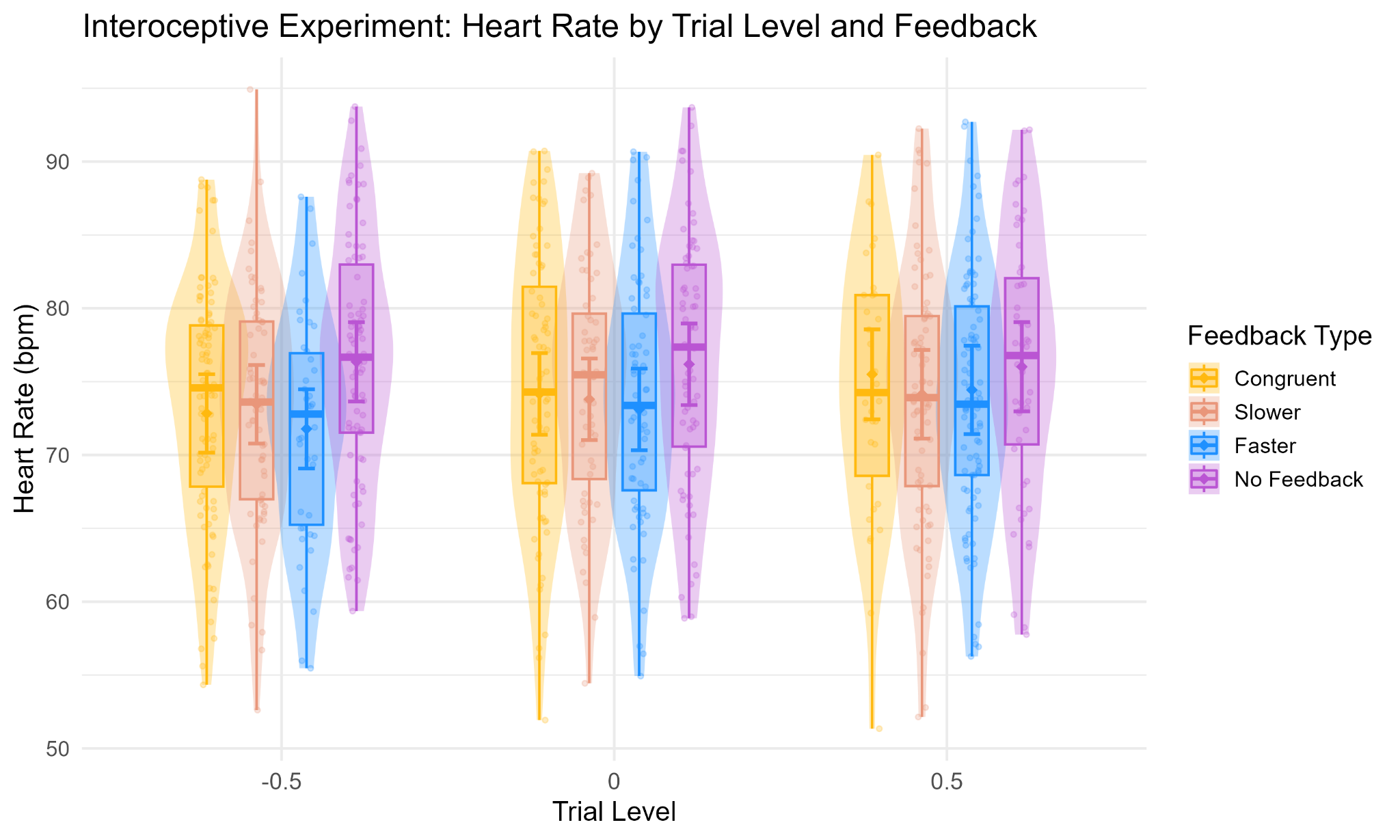

*Supplementary Figure 1.* Interoceptive experiment: Model-predicted heart rate (HR) responses as a function of standardized trial progression (Trial) across feedback conditions (Feedback × Trial was significant, F(3, 711.19) = 6.16, p < .001). The Trial variable is modeled as a continuous, mean-centered predictor capturing temporal dynamics within the session (from -0.5 = early trials to +0.5 = late trials). Violin plots show the distribution and density of predicted HR values at early, middle, and late trial points.

##### 2.2 Likert pain unpleasantness

To examine variations in pain unpleasantness ratings within the interoceptive experiment, we applied general linear mixed-effects modeling (GLMM), the model was fitted using R (lme4 package; Bates et al., 2015; Bates et al., 2015) and was specified as follows:

LIKERT PAIN UNPLEASANTNESS RATINGS ~ Feedback × StimInt × Trial + (StimInt + Trial | Subject)

where Feedback, StimInt, and Trial were included as fixed effects. A random slope for StimInt and Trial was included at the subject level (StimInt + Trial | Subject) to account for individual differences in pain perception across stimulus intensities and over time. Model comparison confirmed that including these random slopes significantly improved model fit (χ² = 13.493 p = 0.001).

###### 2.2.1 Results

###### 2.2.1.1 Effects of interest

The Type III ANOVA revealed a significant **main effect of Feedback**, F(3, 669.05) = 19.60, p < .0001, indicating that pain unpleasantness ratings varied as a function of the type of feedback received. Compared to the Congruent condition (M = 3.02, SE = 0.11), unpleasantness ratings were significantly higher in the Faster feedback condition (M = 3.17, SE = 0.11; estimate = –0.14, SE = 0.05, t(670) = –2.92, p = .0044), and significantly lower in the No Feedback condition (M = 2.80, SE = 0.11; estimate = 0.22, SE = 0.05, t(669) = 4.58, p < .0001). In contrast, the Slower feedback condition (M = 2.97, SE = 0.11) did not significantly differ from the Congruent condition (estimate = -0.05, SE = 0.05, t(668) = -0.99, p = .3112).

Post hoc comparisons further clarified these effects. Ratings in the No Feedback condition were significantly lower than in all other feedback conditions (all ps < .0005). Additionally, unpleasantness ratings were significantly higher in the Faster feedback condition compared to both the Slower (estimate = 0.19, SE = 0.05, t(669) = 4.02, p = .0001) and Congruent conditions (p = .0044). The Slower condition did not significantly differ from Congruent (p = .3112), but was associated with significantly higher unpleasantness than the No Feedback condition (estimate = 0.17, SE = 0.05, t(669) = 3.67, p = .0004).

###### 2.2.1.1 Additional unpredicted effects

A robust **main effect of Stimulus Intensity** also emerged, F(1, 34.04) = 169.12, p < .0001, indicating that unpleasantness ratings increased significantly with higher stimulus intensities. The fixed-effect estimate confirmed a strong positive relationship (b = 1.52, SE = 0.12, t(89.6) = 12.18, p < .0001), suggesting that stronger pain stimulation was reliably associated with higher unpleasantness evaluations.

A Type III ANOVA revealed a significant **main effect of Trial** on pain unpleasantness ratings, F(1, 34.46) = 11.30, p = .0019, suggesting that unpleasantness ratings varied throughout the experimental session. However, the fixed-effect estimate of the Trial coefficient did not reach statistical significance (b = 0.25, SE = 0.16, t(140.90) = 1.57, p = .118), indicating a non-significant trend toward increasing unpleasantness ratings over time. These interactions are reported in detail below.

The interaction between **Stimulus Intensity × Feedback** was significant, F(3, 672.32) = 3.90, p = .0088, indicating that the effect of stimulus intensity on pain unpleasantness ratings varied depending on the type of feedback received (See Supplementary Figure 2). To further examine this interaction, estimated marginal means (EMMs) of unpleasantness ratings were computed at three levels of stimulus intensity: low (–0.5), medium (0), and high (+0.5).

The interaction between Stimulus Intensity and Feedback was significant, *F*(3, 672.32) = 3.90, *p* = .0088, indicating that the effect of stimulus intensity on pain unpleasantness ratings varied as a function of the type of feedback received. To explore this interaction, estimated marginal means (EMMs) of unpleasantness were examined at three levels of stimulus intensity: low (–0.5), medium (0), and high (+0.5).

At low stimulus intensity (–0.5), the highest unpleasantness ratings were observed under the Faster feedback condition (*M* = 2.58, *SE* = 0.10), followed by Slower (*M* = 2.36, *SE* = 0.10), Congruent (*M* = 2.26, *SE* = 0.11), and No Feedback (*M* = 2.23, *SE* = 0.10). Post hoc comparisons showed that Faster feedback elicited significantly higher ratings than both Congruent (estimate = –0.32, *SE* = 0.08, *t*(675) = –3.97, *p* = .0002) and No Feedback (estimate = 0.35, *SE* = 0.08, *t*(673) = 4.55, *p* < .0001). Ratings under Faster were also significantly higher than Slower (estimate = 0.22, *SE* = 0.08, *t*(675) = –2.84, *p* = .0094). No other contrasts reached significance.

At medium intensity (0), the Faster condition again yielded the highest ratings (*M* = 3.17, *SE* = 0.11), followed by Congruent (*M* = 3.02, *SE* = 0.11), Slower (*M* = 2.97, *SE* = 0.11), and No Feedback (*M* = 2.80, *SE* = 0.11). Ratings under Faster were significantly higher than Congruent (estimate = –0.14, *SE* = 0.05, *t*(670) = –2.93, *p* = .0042), Slower (estimate = –0.19, *SE* = 0.05, *t*(669) = –4.02, *p* = .0001), and No Feedback (estimate = 0.37, *SE* = 0.05, *t*(671) = 7.61, *p* < .0001). Both Congruent and Slower feedback also produced significantly higher ratings than No Feedback (all *ps* < .001).

At high intensity (+0.5), unpleasantness ratings peaked under Congruent feedback (*M* = 3.78, *SE* = 0.14), followed by Faster (*M* = 3.76, *SE* = 0.14), Slower (*M* = 3.59, *SE* = 0.14), and No Feedback (*M* = 3.37, *SE* = 0.14). No Feedback elicited significantly lower ratings than all other conditions (all *ps* < .01). In addition, both Congruent (estimate = 0.20, *SE* = 0.08, *t*(671) = 2.52, *p* = .0179) and Faster (estimate = 0.17, *SE* = 0.08, *t*(671) = 2.21, *p* = .0326) conditions elicited significantly higher ratings than Slower. No significant difference was found between Congruent and Faster (estimate = 0.03, *SE* = 0.08, *t*(670) = 0.33, *p* = .7415).

Unpleasantness ratings increased significantly with stimulus intensity across all feedback conditions. However, the rate of increase (slope) varied by feedback type. The steepest increase was observed in the Congruent condition (b = 1.52, SE = 0.13, t(88.6) = 12.19, p < .0001), followed by Slower (b = 1.23, SE = 0.12, t(76.6) = 10.23, p < .0001), Faster (b = 1.18, SE = 0.12, t(85.0) = 9.56, p < .0001), and No Feedback (b = 1.14, SE = 0.12, t(77.5) = 9.51, p < .0001). Pairwise comparisons of slopes (FDR-corrected) confirmed that the slope in the Congruent condition was significantly steeper than in Slower (estimate = 0.29, SE = 0.12, t(672) = 2.43, p = .0305), Faster (estimate = 0.34, SE = 0.12, t(673) = 2.76, p = .0177), and No Feedback (estimate = 0.38, SE = 0.12, t(672) = 3.12, p = .0114). No other pairwise slope differences reached significance (all ps > .70).

The interaction between **Feedback × Trial** was significant, F(3, 675.33) = 3.44, p = .0165, indicating that the trajectory of unpleasantness ratings across trials varied depending on the type of feedback received (See Supplementary Figure 3).
To further unpack this interaction, the estimated slopes of Trial were computed for each feedback condition. Unpleasantness ratings increased significantly over time in the Slower feedback condition (b = 0.35, SE = 0.14, 95% CI [0.07, 0.62], t(91) = 2.50, p = .0143), and even more robustly in the No Feedback condition (b = 0.69, SE = 0.16, 95% CI [0.38, 1.00], t(134) = 4.43, p < .0001). In contrast, the increase observed in the Congruent (b = 0.25, SE = 0.16, 95% CI [–0.06, 0.56], t(142) = 1.58, p = .1174) and Faster conditions (b = 0.18, SE = 0.14, 95% CI [–0.10, 0.46], t(101) = 1.26, p = .2122) did not reach statistical significance.

Post hoc pairwise comparisons of slopes (FDR-corrected) confirmed that the rate of increase in unpleasantness over trials was significantly higher in the No Feedback condition compared to both Congruent (estimate = –0.44, SE = 0.18, t(679) = –2.46, p = .0430) and Faster feedback (estimate = –0.51, SE = 0.17, t(677) = –3.05, p = .0144). The difference between No Feedback and Slower was marginally significant (estimate = –0.34, SE = 0.16, t(676) = –2.10, p = .0730), while no significant differences were found among the active feedback conditions (all ps > .40).

To further explore how the effect of feedback on unpleasantness evolved over time, estimated marginal means were computed at three levels of the standardized Trial variable (–0.5, 0, +0.5), representing early, middle, and late stages of the session, respectively. This analysis allowed us to assess whether the differences between feedback conditions were stable or changed over time.

At *early trials* (Trial = –0.5), unpleasantness ratings were highest in the Faster condition (M = 3.08, SE = 0.13), followed by Congruent (M = 2.90, SE = 0.13), Slower (M = 2.80, SE = 0.13), and No Feedback (M = 2.46, SE = 0.13). Ratings in the No Feedback condition were significantly lower than in all other feedback conditions (all ps < .0007). Moreover, Faster feedback elicited significantly higher ratings than Slower (estimate = –0.28, SE = 0.10, t(671) = –2.90, p = .0058) and marginally higher ratings than Congruent (estimate = –0.18, SE = 0.09, t(671) = –1.88, p = .0733).

At *middle trials* (Trial = 0), the pattern remained consistent. Unpleasantness ratings were again highest in the Faster condition (M = 3.17, SE = 0.11), followed by Congruent (M = 3.02, SE = 0.11), Slower (M = 2.98, SE = 0.11), and No Feedback (M = 2.80, SE = 0.11). Ratings in the No Feedback condition were significantly lower than in all other feedback types (all ps < .0005), and Faster was significantly higher than both Congruent (estimate = –0.14, SE = 0.05, t(670) = –2.91, p = .0045) and Slower (estimate = –0.19, SE = 0.05, t(669) = –4.01, p = .0001).

By *late trials* (Trial = 0.5), feedback-related differences in unpleasantness ratings had largely dissipated, with no statistically significant differences among feedback conditions (all ps > .55). Nonetheless, the Faster condition continued to elicit the highest unpleasantness ratings (M = 3.26, SE = 0.12), slightly above Congruent (M = 3.15, SE = 0.14), Slower (M = 3.15, SE = 0.12), and No Feedback (M = 3.15, SE = 0.13). Although these differences were no longer significant, the numerical pattern suggests a persistent trend toward heightened unpleasantness in the presence of accelerated feedback, even in the final stage of the session. This convergence indicates that, over time, the modulatory effect of feedback on unpleasantness attenuated.

Additionally, the effect of Trial on pain unpleasantness ratings varied as a function of the actual intensity of the nociceptive stimulation, as indicated by a significant **Stimulus Intensity × Trial** interaction, *F*(1, 680.64) = 5.44, *p* = .0199. To further investigate this interaction, estimated marginal trends of unpleasantness ratings across trials were computed at three levels of stimulus intensity: low (–0.5), medium (0), and high (+0.5). Unpleasantness ratings increased progressively over trials at all intensity levels, but the rate of increase was steeper as stimulus intensity increased.

At low intensity, the slope was positive but did not reach statistical significance (*b* = 0.20, SE = 0.13, 95% CI [–0.06, 0.45], *t*(68.8) = 1.50, *p* = .1377). A significant upward trend emerged at medium intensity (*b* = 0.37, SE = 0.11, 95% CI [0.14, 0.59], *t*(34.6) = 3.36, *p* = .0019), which became even more pronounced at high intensity (*b* = 0.54, SE = 0.13, 95% CI [0.27, 0.80], *t*(74.7) = 4.04, *p* = .0001).

Pairwise comparisons between slopes (FDR-corrected) confirmed that the increase in unpleasantness ratings over trials was significantly steeper at higher stimulus intensities. Specifically, the slope at low intensity was significantly shallower than both medium (estimate = –0.17, SE = 0.073, *t*(681) = –2.33, *p* = .0201) and high intensity (estimate = –0.34, SE = 0.15, *t*(681) = –2.33, *p* = .0201). Moreover, the slope at medium intensity was also significantly lower than at high intensity (estimate = –0.17, SE = 0.073, *t*(681) = –2.33, *p* = .0201). These results suggest that the impact of repeated nociceptive stimulation on perceived unpleasantness intensifies progressively with the strength of the stimulation.

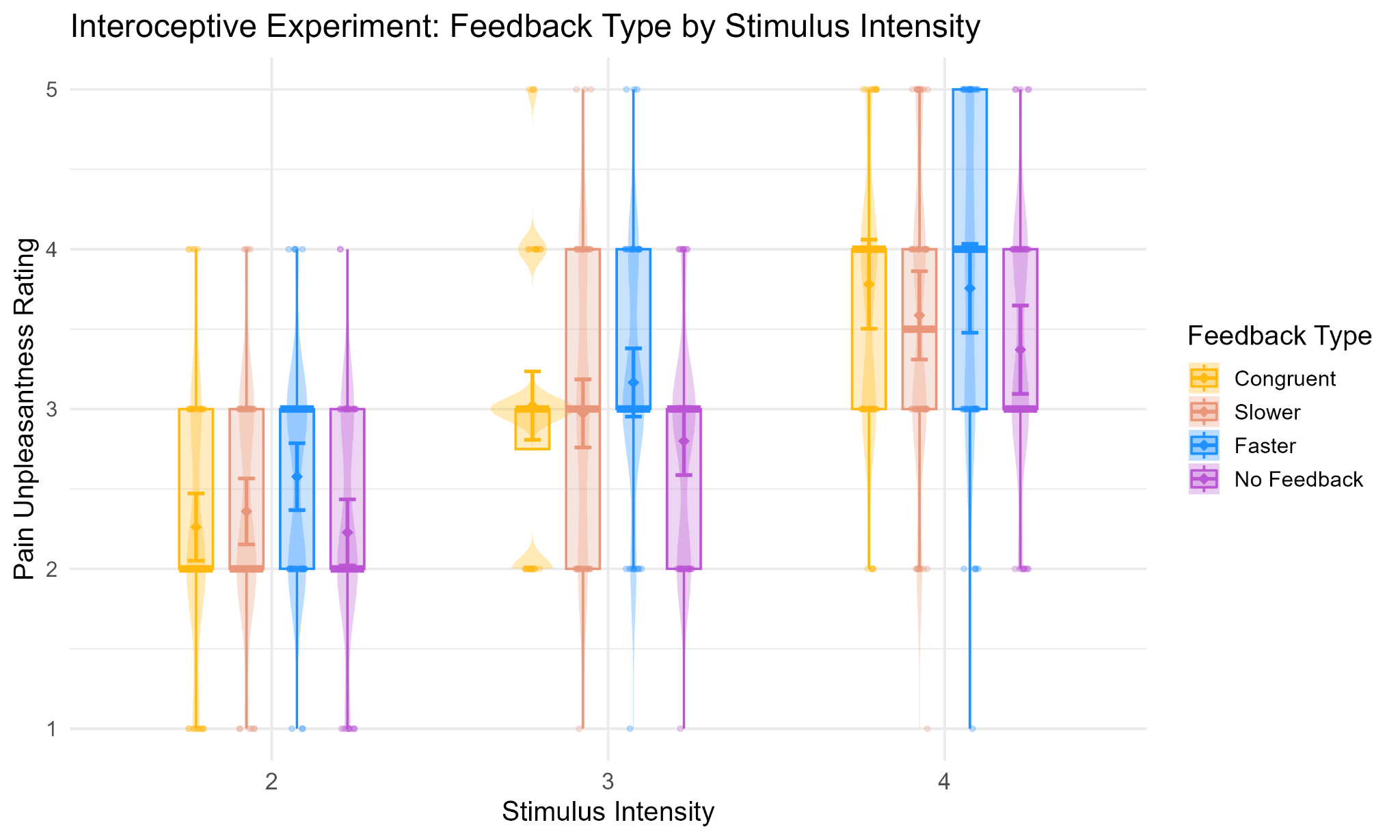

*Supplementary Figure 2.* Interoceptive experiment: Model-predicted Pain Unpleasantness ratings for each stimulation intensity (StimInt 2–4), across feedback conditions (Stimulus Intensity × Feedback was significant, F(3, 672.32) = 3.90, p = .0088). Violin plots and overlaid boxplots depict the distribution (violin) and interquartile range with median (box) of the observed data for each condition. Boxes are centred on the x-axis categories because they summarise the data within each stimulation intensity and feedback condition. Large coloured dots and error bars show the model-predicted estimated marginal means (EMMs) ± standard errors for each condition.

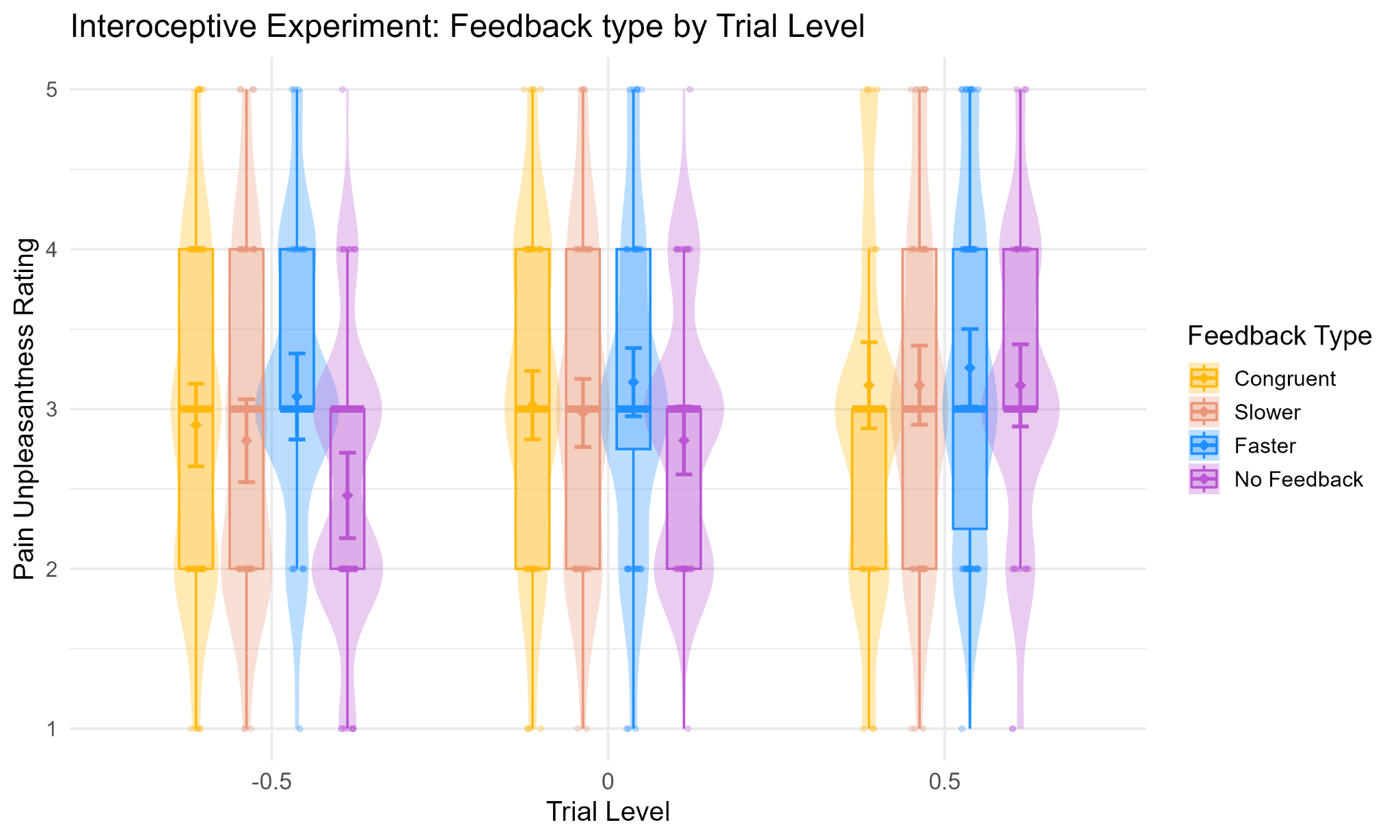

*Supplementary Figure 3.* Interoceptive experiment: Model-predicted pain unplesantness ratings as a function of standardized trial progression (Trial) across feedback conditions (Feedback × Trial, *F*(3, 675.33) = 3.44, *p* = .0165). The Trial variable is modeled as a continuous, mean-centered predictor reflecting the temporal progression of the task (from -0.5 = early trials to +0.5 = late trials). Violin plots display the distribution and density of predicted ratings at early, middle, and late stages of the session

##### 2.3 Numeric Pain Scale of intensity ratings

To examine variations in pain intensity ratings within the interoceptive experiment, we applied general linear mixed-effects modeling (GLMM), the model was fitted using R (lme4 package; Bates et al., 2015; Bates et al., 2015) and was specified as follows:

NUMERIC PAIN SCALE OF INTENSITY RATINGS~ Feedback × StimInt × Trial + (StimInt + Trial | Subject)

where Feedback, StimInt and Trial were included as fixed effects. A random slope for StimInt and Trial was included at the subject level (StimInt + Trial | Subject) to account for individual differences in pain perception across stimulus intensities and over time. Model comparison confirmed that including these random slopes significantly improved model fit (χ² = 83.437, p < 0.0001).

###### 2.3.1 Results

###### 2.3.1.1 Effects of interest

The results of the Type III ANOVA with Satterthwaite’s approximation revealed a significant **main effect of Feedback**, F(3, 663.89) = 22.46, p < .001, indicating that Numeric pain scale of intensity ratings varied across feedback conditions. Compared to the reference condition (i.e., Congruent feedback), Numeric pain scale of intensity ratings were significantly higher in the Faster feedback condition (b = 3.69, SE = 1.08, t(664.80) = 3.40, p = .0008), and significantly lower in the No Feedback condition (b = –4.90, SE = 1.07, t(663.18) = –4.59, p < .0001). The Slower feedback condition did not differ significantly from the Congruent condition (b = –0.79, SE = 1.07, t(663.03) = –0.74, p = .4623).

Post hoc comparisons further clarified these effects. Numeric pain scale of intensity ratings were significantly higher in the Faster feedback condition compared to Congruent (*estimate* = -3.691, *SE* = 1.08, *t*(665 )=-3.406, *p* = .0008), Slower (*estimate* = -4.522, *SE* = 1.06, *t*(665 )=-4.285, p < .0001), and No Feedback (*estimate* = 8.711, *SE* = 1.05, *t*(665)=8.260, p < .0001). The Slower condition did not differ significantly from the Congruent condition (p = .4350). Moreover, No Feedback yielded significantly lower Numeric pain scale of intensity ratings compared to all other feedback conditions (all ps < .0001).

###### 2.3.1.2 Additional unpredicted effects

A Type III ANOVA revealed a significant **main effect of Trial** on Numeric pain scale of intensity ratings, F(1, 32.90) = 9.01, p = .0051, suggesting that pain ratings varied throughout the experimental session. However, the fixed-effect estimate of the Trial coefficient from the mixed model indicated a positive but non-significant trend (b = 6.05, SE = 3.73, t(103.82) = 1.62, p = .108), suggesting that pain intensity ratings tended to increase over time, although this trend was not statistically reliable when averaged across all other factors in the model. This apparent discrepancy likely reflects the fact that Trial was involved in several higher-order interactions, particularly with Stimulus Intensity and Feedback, that modulated its effect on pain ratings. These interactions are reported in detail below.

Critically, there was also a robust **main effect of Stimulus Intensity**, F(1, 34.22) = 204.11, p < .001, indicating that pain ratings increased as a function of the objective intensity of nociceptive stimulation. The fixed-effect estimate showed a strong positive association (b = 37.12, SE = 2.82, t(83.01) = 13.14, p < .001), consistent with the expected relationship between stimulus intensity and perceived pain.

The **Feedback × Stimulus Intensity** interaction was also significant, F(3, 667.07) = 3.4631, p = .016, indicating that the effect of stimulus intensity on pain ratings varied across feedback conditions (See Supplementary Figure 4). To further unpack this interaction, estimated marginal means (EMMs) of Numeric pain scale of intensity ratings were computed at the three levels of stimulus intensity: low (–0.5), medium (0), and high (+0.5).

At *low stimulus intensity* (–0.5), the Faster feedback condition elicited the highest pain ratings (M = 43.4, SE = 2.76), followed by Slower (M = 39.6, SE = 2.74), Congruent (M = 37.2, SE = 2.78), and No Feedback (M = 36.3, SE = 2.74).

Ratings in the Faster condition were significantly higher than those in the Congruent (estimate = -6.260, SE = 1.74, t(668) = -3.589, p = .0011), Slower (estimate = -3.872, SE = 1.69, t(671) = -2.289, p = .0448) and No Feedback (estimate = 7.185, SE = 1.68, t(668) = 4.274, p = .0001).

At *medium intensity* (0), Faster again produced the highest ratings (M = 59.4, SE = 2.76), followed by Congruent (M = 55.7, SE = 2.76), Slower (M = 54.9, SE = 2.75), and No Feedback (M = 50.7, SE = 2.75). Post hoc comparisons confirmed that Faster was significantly higher than both Congruent (estimate = -3.721, SE = 1.08, t(665) = -3.432, p = .0008) and Slower (estimate = -4.515, SE = 1.06, t(665) = -4.277, p < .0001), while all three active feedback conditions yielded significantly higher ratings than No Feedback (all ps < .0002).

At *high intensity* (+0.5), Faster again elicited the highest ratings (M = 75.4, SE = 3.38), followed by Congruent (M = 74.2, SE = 3.39), Slower (M = 70.3, SE = 3.37), and No Feedback (M = 65.2, SE = 3.37). The No Feedback condition was again significantly lower than all others (all ps < .0035), and Faster was significantly higher than Slower (estimate = -5.157, SE = 1.67, t(665) = -3.091, p = .0032). A significant difference was also observed between Congruent and Slower (estimate = 3.976, SE = 1.68, t(666) = 2.362, p = .0221), while the difference between Congruent and Faster was not significant (p = .4904).

Numeric pain scale of intensity ratings increased as a function of stimulus intensity in all feedback conditions. However, the rate of increase (slope) differed depending on the feedback type. The steepest increase was observed in the Congruent condition (b = 37.1, SE = 2.82, t(81) = 13.161, p < .0001), followed by Faster (b = 32.0, SE = 2.78, t(77.7) = 11.480, p < .0001), Slower (b = 30.7, SE = 2.74, t(72.5) = 11.212, p < .0001) and No Feedback (b = 29.0, SE = 2.73, t(71.7) = 10.612, p < .0001). Pairwise comparisons between slopes revealed that the increase in pain ratings was significantly steeper in the Congruent condition compared to Slower (estimate = 6.36, SE = 2.64, t(668) = 2.411, p = .048) and No feedback (estimate = 8.10, SE = 2.64, t(668) = 3.07, p = .0133. Differences between other slopes did not reach statistical significance (all ps > .11).

The interaction between **Feedback × Trial** was significant, *F*(3, 669.15) = 9.86, *p* < .001, indicating that the trajectory of Numeric pain scale of intensity ratings across trials varied depending on the type of feedback received (See Supplementary Figure 5).
To further unpack this interaction, the estimated slopes of Trial were computed for each feedback condition. Pain intensity ratings increased significantly over time in the Slower feedback condition (*b* = 7.69, *SE* = 3.37, 95% CI [0.97, 14.41], *t*(73.3) = 2.28, *p* = .0255), and even more robustly in the No Feedback condition (*b* = 19.61, *SE* = 3.68, 95% CI [12.31, 26.91], *t*(101.1) = 5.33, *p* < .0001).

Post hoc pairwise comparisons of slopes (FDR-corrected) confirmed that the rate of increase in pain intensity over trials was significantly higher in the No Feedback condition compared to Congruent (estimate = –13.48, *SE* = 3.93, *t*(673) = –3.43, *p* = .0018), Slower (estimate = –11.92, *SE* = 3.57, *t*(670) = –3.34, *p* = .0018), and Faster (estimate = –19.62, *SE* = 3.66, *t*(671) = –5.37, *p* < .0001). Additionally, the slope in the Slower condition was significantly higher than in the Faster condition (estimate = 7.70, *SE* = 3.34, *t*(668) = 2.30, *p* = .0323), while no significant differences were found among the active feedback conditions (all *ps* > .11).

To further explore how the effect of feedback on pain intensity evolved over time, estimated marginal means were computed at three levels of the standardized Trial variable (–0.5, 0, +0.5), representing early, middle, and late stages of the session. This analysis allowed us to assess whether the differences between feedback conditions were stable or changed over time.

At *early trials* (Trial = –0.5), pain intensity ratings were highest in the Faster condition (M = 59.6, SE = 3.33), followed by Congruent (M = 52.9, SE = 3.23), Slower (M = 51.3, SE = 3.23), and No Feedback (M = 41.2, SE = 3.31). Ratings in the No Feedback condition were significantly lower than in all other feedback conditions (all ps < .0001). Moreover, Faster feedback elicited significantly higher pain ratings than both Congruent (estimate = –6.73, SE = 2.09, t(666) = –3.23, p = .0016) and Slower (estimate = –8.33, SE = 2.08, t(666) = –3.99, p = .0001).

At *middle trials* (Trial = 0), the pattern remained consistent: Faster feedback again produced the highest pain ratings (M = 59.6, SE = 2.76), followed by Congruent (M = 56.0, SE = 2.77), Slower (M = 55.1, SE = 2.75), and No Feedback (M = 51.0, SE = 2.75). Ratings in the No Feedback condition were significantly lower than all others (all ps < .0005). Additionally, Faster was significantly higher than both Congruent (estimate = –3.66, SE = 1.09, t(666) = –3.37, p = .0010) and Slower (estimate = –4.48, SE = 1.05, t(665) = –4.26, p < .0001).

By *late trials* (Trial = 0.5), feedback-related differences in pain ratings had largely dissipated, with no statistically significant contrasts among conditions (all ps > .94). Nevertheless, Faster still elicited numerically higher pain ratings (M = 59.6, SE = 3.17) compared to Congruent (M = 59.0, SE = 3.44), Slower (M = 59.0, SE = 3.23), and No Feedback (M = 60.8, SE = 3.31), suggesting a potential residual effect of feedback on pain perception. However, this trend was no longer statistically significant, indicating that the modulatory influence of feedback on the Numeric Pain Scale of intensity ratings diminished over time.

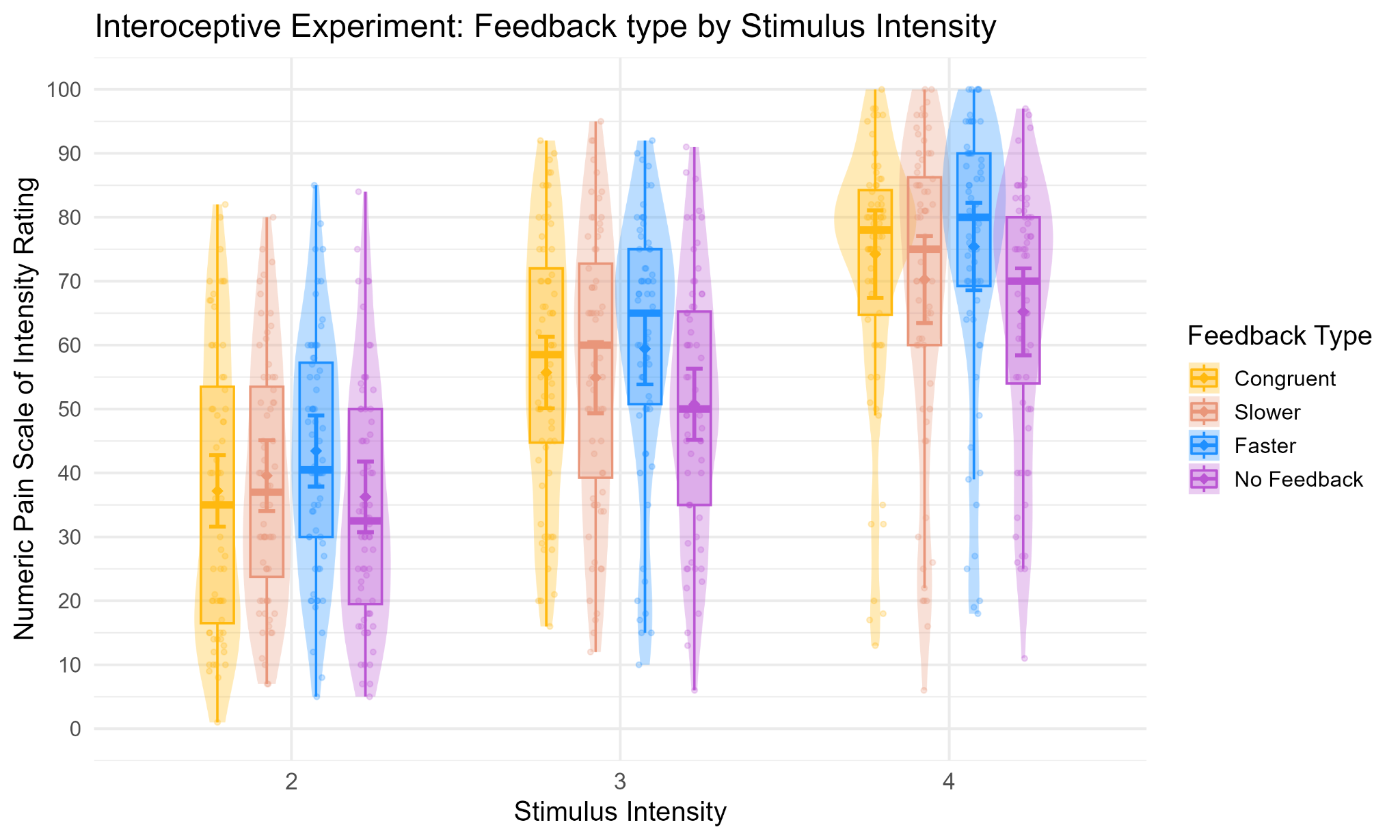

*Supplementary Figure 4.* Interoceptive experiment: Model-predicted Numeric Pain Scale of intensity ratings (NPS) for each stimulation intensity (StimInt 2–4), across feedback conditions (Feedback × Stimulus Intensity, F(3, 667.07) = 3.4631, p = .016). Violin plots and overlaid boxplots depict the distribution (violin) and interquartile range with median (box) of the observed data for each condition. Boxes are centred on the x-axis categories because they summarise the data within each stimulation intensity and feedback condition. Large coloured dots and error bars show the model-predicted estimated marginal means (EMMs) ± standard errors for each condition.

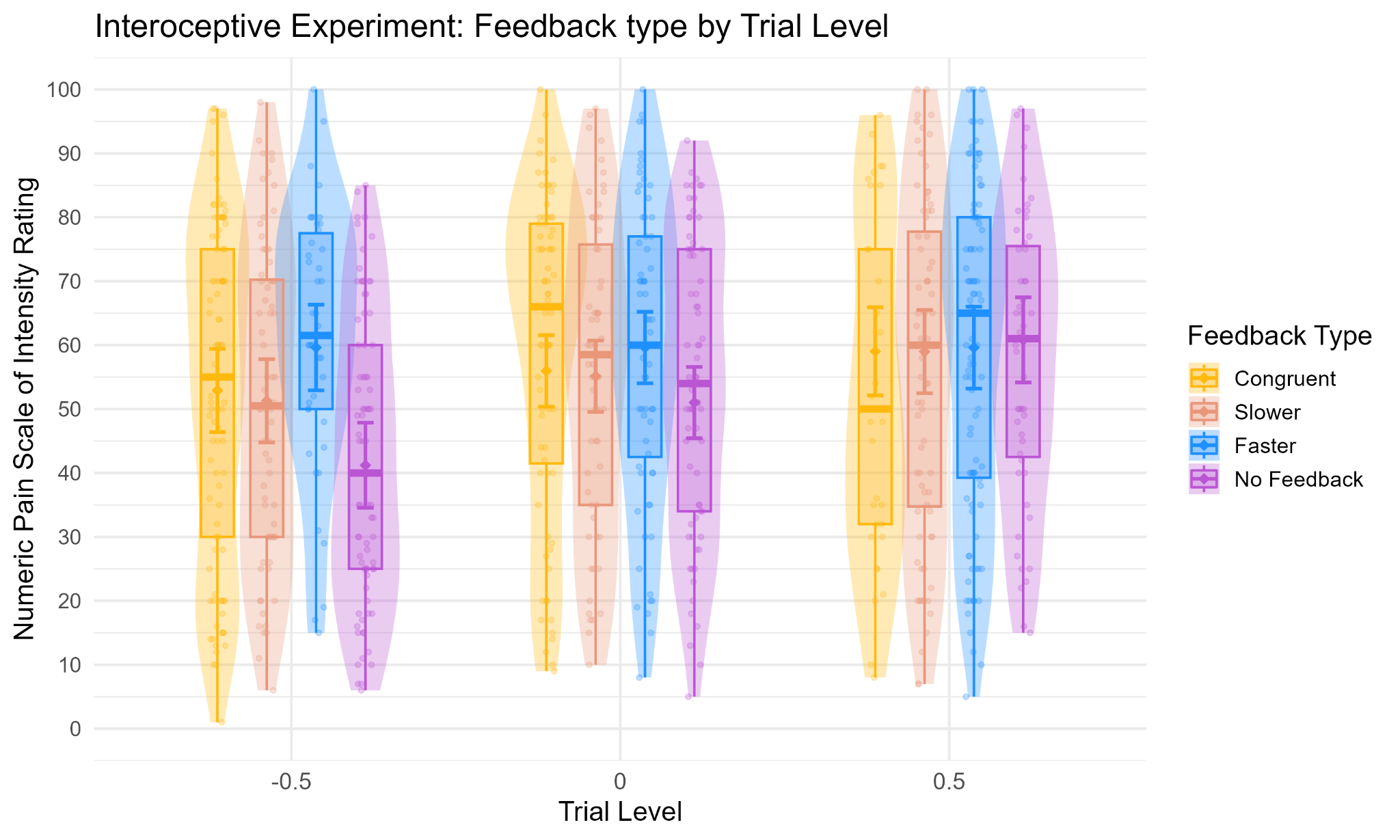

*Supplementary Figure 5.* Interoceptive experiment: Model-predicted Numeric Pain Scale of intensity ratings (NPS) as a function of standardized trial progression (Trial) across feedback conditions (Feedback × Trial, *F*(3, 669.15) = 9.86, *p* < .001). The Trial variable is modeled as a continuous, mean-centered predictor reflecting the temporal progression of the task (from -0.5 = early trials to +0.5 = late trials). Violin plots display the distribution and density of predicted ratings at early, middle, and late stages of the session.

#### 3. Exteroceptive experiment (within-subjects model)

##### 3.1 Heart rate

To examine variations in heart rate (HR) within the exteroceptive experiment, we applied general linear mixed-effects modeling (GLMM), the model was fitted using R (lme4 package; Bates et al., 2015; Bates et al., 2015) and was specified as follows:

𝐻𝑅∼𝐹𝑒𝑒𝑑𝑏𝑎𝑐𝑘×𝑇𝑟𝑖𝑎𝑙+(𝑇𝑟𝑖𝑎𝑙∣𝑆𝑢𝑏𝑗𝑒𝑐𝑡)

where Feedback and Trial were included as fixed effects, and a random slope for Trial at the subject level (Trial | Subject) to account for individual variability in HR trajectories over time. Model comparison indicated that the inclusion of a random slope significantly improved model fit (χ² = 11.426, *p* = .0435).

###### 3.1.1 Results

###### 3.1.1.1 Effects of interest

The Type III ANOVA revealed a significant **main effect of Feedback** on heart rate (HR), F(3, 596.03) = 44.43, p < .0001, indicating that HR levels varied depending on the type of feedback received. Compared to the Congruent condition (M = 76.5, SE = 1.67), HR was significantly higher in the No Feedback condition (M = 78.8, SE = 1.67; estimate = 2.19, SE = 0.25, t(597.5) = 8.92, p < .0001). In contrast, neither the Slower (M = 76.3, SE = 1.67; estimate = –0.29, SE = 0.24, t(594.9) = –1.21, p = .2285) nor the Faster feedback condition (M = 76.5, SE = 1.67; estimate = –0.03, SE = 0.24, t(595.1) = –0.11, p = .9088) significantly differed from the Congruent condition.

Post hoc comparisons further clarified these effects. Relative to all other conditions, the No Feedback condition elicited significantly higher HR values (all ps < .0001). Specifically, HR was significantly higher in the No Feedback condition compared to both the Slower (estimate = –2.50, SE = 0.24, t(598) = –10.29, p < .0001) and Faster conditions (estimate = –2.24, SE = 0.24, t(598) = –9.22, p < .0001).

*No significant differences were found among the active feedback conditions (all ps > .33)*, suggesting that the HR modulation was driven specifically by the absence of feedback, rather than differences among feedback types.

No other main effects or interactions reached statistical significance (See Supplementary Figure 6 for illustrative purposes).

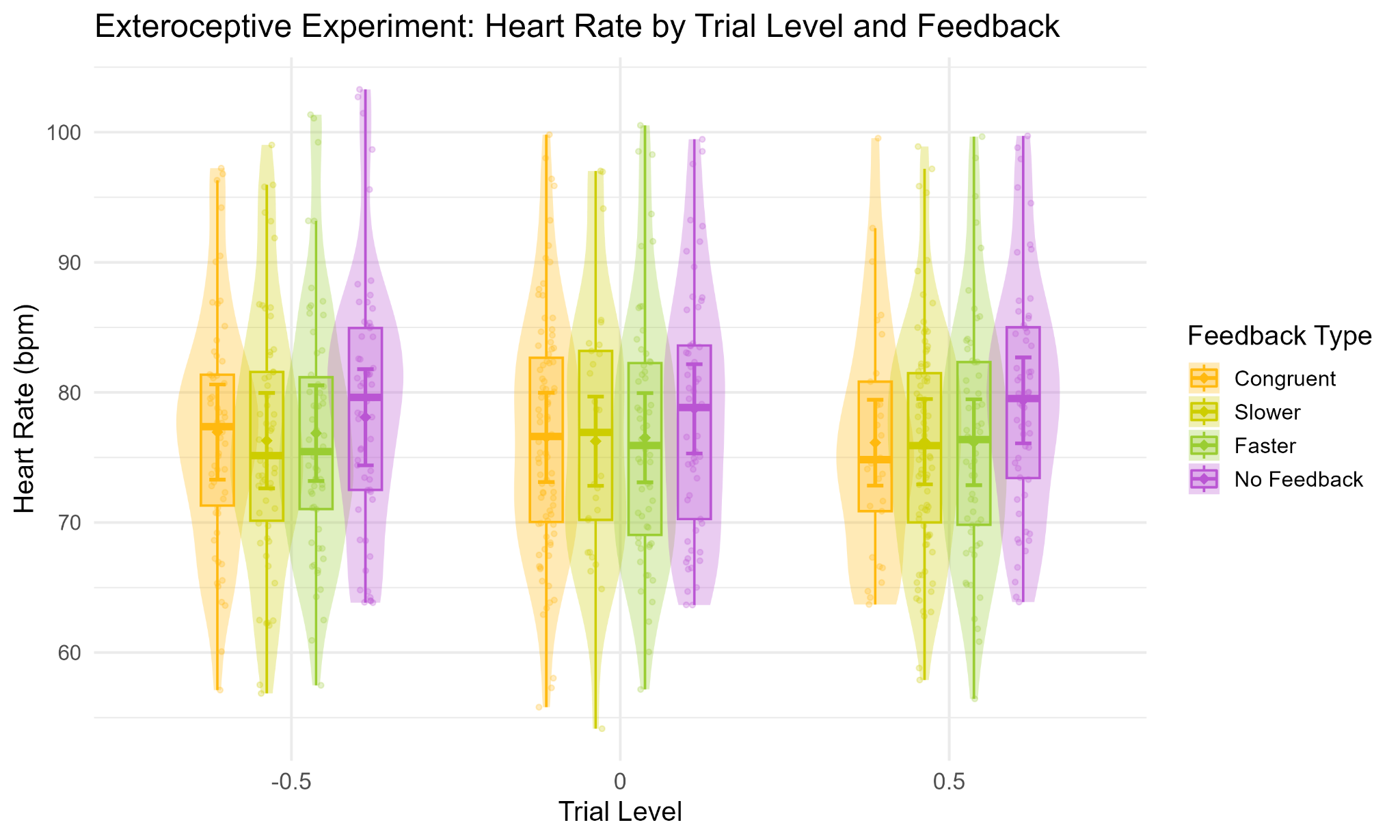

*Supplementary Figure 6.* Exteroceptive experiment: Model-predicted heart rate (HR) responses as a function of standardized trial progression (Trial) across feedback conditions in the Exteroceptive condition. The Trial variable is modeled as a continuous, mean-centered predictor capturing temporal dynamics within the session (from -0.5 = early trials to +0.5 = late trials). Violin plots show the distribution and density of predicted HR values at early, middle, and late trial points. Feedback x Trial was not significant, values are shown for consistency with the interoceptive experiment.

##### 3.2 Likert pain unpleasantness

To examine variations in pain unpleasantness ratings within the exteroceptive experiment, we applied general linear mixed-effects modeling (GLMM), the model was fitted using R (lme4 package; Bates et al., 2015; Bates et al., 2015) and was specified as follows:

LIKERT PAIN UNPLEASANTNESS RATINGS ~ Feedback × StimInt × Trial + (StimInt + Trial | Subject)

where Feedback, StimInt and Trial were included as fixed effects. A random slope for StimInt and Trial was included at the subject level (StimInt + Trial | Subject) to account for individual differences in pain perception across stimulus intensities and over time. Model comparison confirmed that including these random slopes significantly improved model fit (*χ²* = 51.538, *p* < 0.001).

###### 3.2.1 Results

###### 3.2.1.1 Effects of interest

The Type III ANOVA revealed a significant main effect of **Feedback** on pain unpleasantness ratings, *F*(3, 563.76) = 22.12, *p* < .0001, indicating that ratings varied as a function of the type of feedback received. Compared to the Congruent condition (M = 3.05, SE = 0.11), unpleasantness ratings were significantly lower in the No Feedback condition (M = 2.76, SE = 0.11; estimate = 0.29, SE = 0.05, *t*(565) = 6.00, *p* < .0001). In contrast, neither the Slower (M = 3.06, SE = 0.11; estimate = –0.01, SE = 0.05, *t*(564) = –0.20, *p* = .8434) nor the Faster condition (M = 3.10, SE = 0.11; estimate = –0.05, SE = 0.05, *t*(564) = –1.07, *p* = .4301) differed significantly from the Congruent condition.

Post hoc comparisons further clarified these effects. Ratings in the No Feedback condition were significantly lower than in all other feedback conditions (all *ps* < .0001). Specifically, unpleasantness ratings were significantly higher in the Faster condition compared to No Feedback (estimate = 0.34, SE = 0.05, *t*(564) = 7.17, *p* < .0001), as well as in the Slower condition compared to No Feedback (estimate = 0.30, SE = 0.05, *t*(564) = 6.31, *p* < .0001), and in the Congruent compared to the No feedback (estimate = 0.28, SE = 0.04, *t*(565) = 6.00, *p* < .0001.

*No significant differences emerged between active feedback conditions (all ps > .43)*, suggesting that the changes in pain ratings were driven specifically by the absence of feedback, rather than differences among feedback types.

###### 3.2.1.2 Additional unpredicted effects

A robust main effect of **Stimulus Intensity** also emerged, *F*(1, 28.27) = 190.00, *p* < .0001, indicating that unpleasantness ratings increased significantly with higher intensity stimulation. The fixed-effect estimate confirmed a strong positive association between stimulus intensity and unpleasantness ratings (b = 1.54, SE = 0.12, *t*(74.01) = 12.98, *p* < .0001), suggesting that more intense stimuli were consistently evaluated as more unpleasant.

Finally, a significant main effect of **Trial** was observed, *F*(1, 29.63) = 7.85, *p* = .0089, suggesting that unpleasantness ratings varied over the course of the experimental session. However, the fixed-effect estimate of the Trial coefficient did not reach statistical significance (b = 0.21, SE = 0.15, *t*(89.91) = 1.40, *p* = .165), indicating only a non-significant trend toward increasing unpleasantness ratings across trials. This apparent discrepancy likely reflects the presence of significant higher-order interactions with Feedback and Stimulus Intensity, which are described in detail in the following sections.

The interaction between **Stimulus Intensity × Feedback** was significant, *F*(3, 569.16) = 8.21, *p* < .0001, indicating that the effect of stimulus intensity on pain unpleasantness ratings varied depending on the type of feedback received (See Supplementary Figure 7). To further examine this interaction, estimated marginal means (EMMs) of unpleasantness ratings were computed at three levels of stimulus intensity: low (–0.5), medium (0), and high (+0.5).

At *low stimulus intensity* (–0.5), Slower feedback elicited the highest unpleasantness ratings (M = 2.56, SE = 0.12), followed by Faster (M = 2.40, SE = 0.12), Congruent (M = 2.28, SE = 0.12), and No Feedback (M = 2.17, SE = 0.12). Post hoc comparisons revealed that unpleasantness ratings in the Slower condition were significantly higher than in the Congruent (estimate = –0.28, SE = 0.08, *t*(570) = –3.64, *p* = .0009) and No Feedback conditions (estimate = 0.39, SE = 0.08, *t*(567) = 5.19, *p* < .0001). Ratings in the Faster condition were also significantly higher than in No Feedback (estimate = 0.23, SE = 0.08, *t*(567) = 3.07, *p* = .0045), but did not significantly differ from Congruent nor Slower (p > .05).

At *medium intensity* (0), the pattern shifted slightly, with Faster feedback yielding the highest ratings (M = 3.10, SE = 0.11), followed by Slower (M = 3.06, SE = 0.11), Congruent (M = 3.05, SE = 0.11), and No Feedback (M = 2.76, SE = 0.11). All three active feedback conditions produced significantly higher unpleasantness ratings than No Feedback (all *ps* < .0001). No significant differences were found among the active feedback conditions (all *ps* > .42).

At *high intensity* (+0.5), unpleasantness ratings peaked under Congruent (M = 3.82, SE = 0.14), followed by Faster (M = 3.80, SE = 0.14), Slower (M = 3.56, SE = 0.14), and No Feedback (M = 3.35, SE = 0.14). Ratings in the Congruent condition were significantly higher than in Slower (estimate = 0.26, SE = 0.08, *t*(567) = 3.44, *p* = .0013) and No Feedback (estimate = 0.47, SE = 0.08, *t*(567) = 6.14, *p* < .0001). Similarly, Faster elicited significantly higher ratings than both Slower (estimate = –0.24, SE = 0.07, *t*(566) = –3.24, *p* = .0019) and No Feedback (estimate = 0.45, SE = 0.08, *t*(567) = 5.98, *p* < .0001). No difference was found between Congruent and Faster (p = .8027).

Across all conditions, unpleasantness ratings increased significantly with higher stimulus intensity. However, the rate of increase (slope) varied depending on feedback. The steepest increase was observed in the Congruent condition (b = 1.54, SE = 0.12, *t*(74.5) = 12.96, *p* < .0001), followed by Faster (b = 1.40, SE = 0.12, *t*(70.2) = 11.97, *p* < .0001), No Feedback (b = 1.18, SE = 0.12, *t*(70.8) = 10.08, *p* < .0001), and Slower (b = 1.00, SE = 0.12, *t*(69.6) = 8.58, *p* < .0001). Pairwise comparisons of slopes confirmed that the Congruent condition showed a significantly steeper increase in unpleasantness compared to both Slower (estimate = 0.54, SE = 0.12, *t*(571) = 4.56, *p* < .0001) and No Feedback (estimate = 0.36, SE = 0.12, *t*(569) = 3.02, *p* = .0052). Faster also differed from Slower (estimate = 0.40, SE = 0.12, *t*(569) = 3.43, *p* = .0020), while the difference between Congruent and Faster did not reach significance (p = .2365), nor did the contrast between Faster and No Feedback (p = .0943).

The interaction between **Feedback × Trial** was significant, *F*(3, 568.85) = 11.47, *p* < .0001, indicating that the trajectory of unpleasantness ratings over time varied depending on the feedback condition (See Supplementary Figure 8). To investigate this effect, the slope of Trial was estimated within each feedback level. Unpleasantness ratings increased significantly over time in the No Feedback condition (b = 0.94, SE = 0.17, 95% CI [0.61, 1.27], *t*(139.1) = 5.63, *p* < .0001). In contrast, the slopes for Congruent (b = 0.21, SE = 0.15, 95% CI [–0.09, 0.50], *t*(90.3) = 1.40, *p* = .1649), Slower (b = 0.23, SE = 0.14, 95% CI [–0.04, 0.51], *t*(70.8) = 1.68, *p* = .0973) and Faster feedback (b = –0.14, SE = 0.16, 95% CI [–0.44, 0.17], *t*(109.4) = –0.87, *p* = .3843) were not statistically significant.

Post hoc comparisons of Trial slopes (FDR-corrected) confirmed that the increase in unpleasantness was significantly steeper in the No Feedback condition compared to Congruent (estimate = –0.73, SE = 0.18, *t*(571) = –4.07, *p* = .0001), Slower (estimate = –0.71, SE = 0.17, *t*(570) = –4.11, *p* = .0001), and Faster (estimate = –1.08, SE = 0.19, *t*(570) = –5.76, *p* < .0001). Additionally, the Slower condition showed a significantly steeper slope than the Faster condition (estimate = 0.37, SE = 0.16, *t*(567) = 2.28, *p* = .0343). No significant differences were found among the active feedback conditions (all *ps* > .05).

To examine how feedback differences evolved over time, estimated marginal means were computed at three levels of the standardized Trial variable: early (–0.5), middle (0), and late (+0.5).

At *early trials* (Trial = –0.5), unpleasantness ratings were highest in the Faster condition (M = 3.17, SE = 0.13), followed by Congruent (M = 2.95, SE = 0.12), Slower (M = 2.94, SE = 0.13), and No Feedback (M = 2.28, SE = 0.13). Ratings in the No Feedback condition were significantly lower than in all other conditions (all *ps* < .0001). Additionally, Faster feedback elicited significantly higher ratings than both Slower (*p* = .0215) and Congruent (*p* = .0215).

At the *midpoint* (Trial = 0), the pattern remained consistent: unpleasantness ratings were highest in the Faster condition (M = 3.10, SE = 0.11), followed by Slower (M = 3.06, SE = 0.11), Congruent (M = 3.05, SE = 0.11), and No Feedback (M = 2.75, SE = 0.11). Post hoc comparisons confirmed that the No Feedback condition was rated significantly lower than all others (all *ps* < .0001), while no other differences between active feedback conditions reached significance (all *ps* > .40).

By *late trials* (Trial = 0.5), unpleasantness ratings converged across conditions, with No Feedback (M = 3.23, SE = 0.15), Slower (M = 3.17, SE = 0.14), Congruent (M = 3.15, SE = 0.15), and Faster (M = 3.03, SE = 0.15) showing no statistically significant differences (all *ps* > .35). Despite numerical variability, these findings indicate that feedback-related effects on unpleasantness diminished over time.

These results suggest that feedback-related differences in unpleasantness ratings were most pronounced at the beginning of the session and progressively diminished over time. Notably, the No Feedback condition was associated with a sustained and significantly steeper increase in unpleasantness ratings over trials, highlighting its distinct temporal trajectory.

Additionally, the effect of Trial on pain unpleasantness ratings varied as a function of the actual intensity of the nociceptive stimulation, as indicated by a significant **Stimulus Intensity × Trial interaction**, F(1, 577.49) = 15.61, p < .001. To further investigate this interaction, estimated marginal trends of unpleasantness ratings across trials were computed at three levels of stimulus intensity: low (–0.5), medium (0), and high (+0.5). While unpleasantness ratings tended to increase over trials at all intensity levels, the rate of this increase became progressively steeper with higher stimulus intensities.

At low intensity, the slope was small and non-significant (b = 0.02, SE = 0.13, 95% CI [–0.24, 0.28], t(53.2) = 0.15, p = .8813). A significant positive trend emerged at medium intensity (b = 0.31, SE = 0.11, 95% CI [0.08, 0.54], t(29.7) = 2.80, p = .0089), which became markedly stronger at high intensity (b = 0.60, SE = 0.14, 95% CI [0.33, 0.88], t(68.0) = 4.38, p < .0001).

Pairwise comparisons between slopes (FDR-corrected) confirmed that the increase in unpleasantness ratings over trials was significantly steeper at higher stimulus intensities. Specifically, the slope at low intensity was significantly shallower than both medium (estimate = –0.29, SE = 0.074, t(578) = –3.94, p = .0001) and high intensity (estimate = –0.58, SE = 0.15, t(578) = –3.94, p = .0001). Moreover, the slope at medium intensity was also significantly lower than at high intensity (estimate = –0.29, SE = 0.074, t(578) = –3.94, p = .0001). These findings indicate that the cumulative effect of repeated nociceptive stimulation on perceived unpleasantness intensifies with increasing stimulation strength.

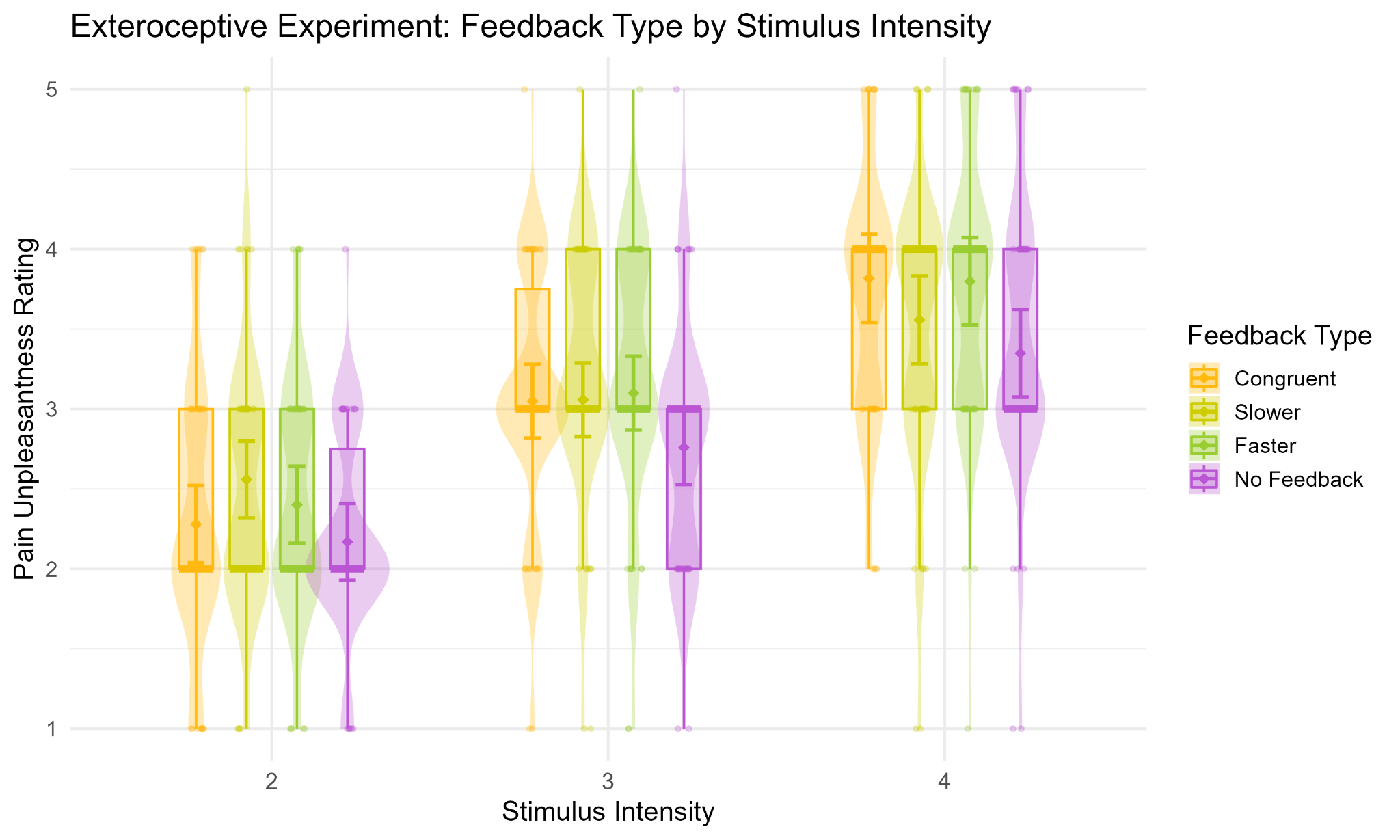

*Supplementary Figure 7.* Exteroceptive experiment: Model-predicted pain unpleasantness ratings for each stimulation intensity (StimInt 2–4), across feedback conditions (Stimulus Intensity × Feedback, F(3, 569.16) = 8.21, p < .0001). Violin plots and overlaid boxplots depict the distribution (violin) and interquartile range with median (box) of the observed data for each condition. Boxes are centred on the x-axis categories because they summarise the data within each stimulation intensity and feedback condition. Large coloured dots and error bars show the model-predicted estimated marginal means (EMMs) ± standard errors for each condition*]*

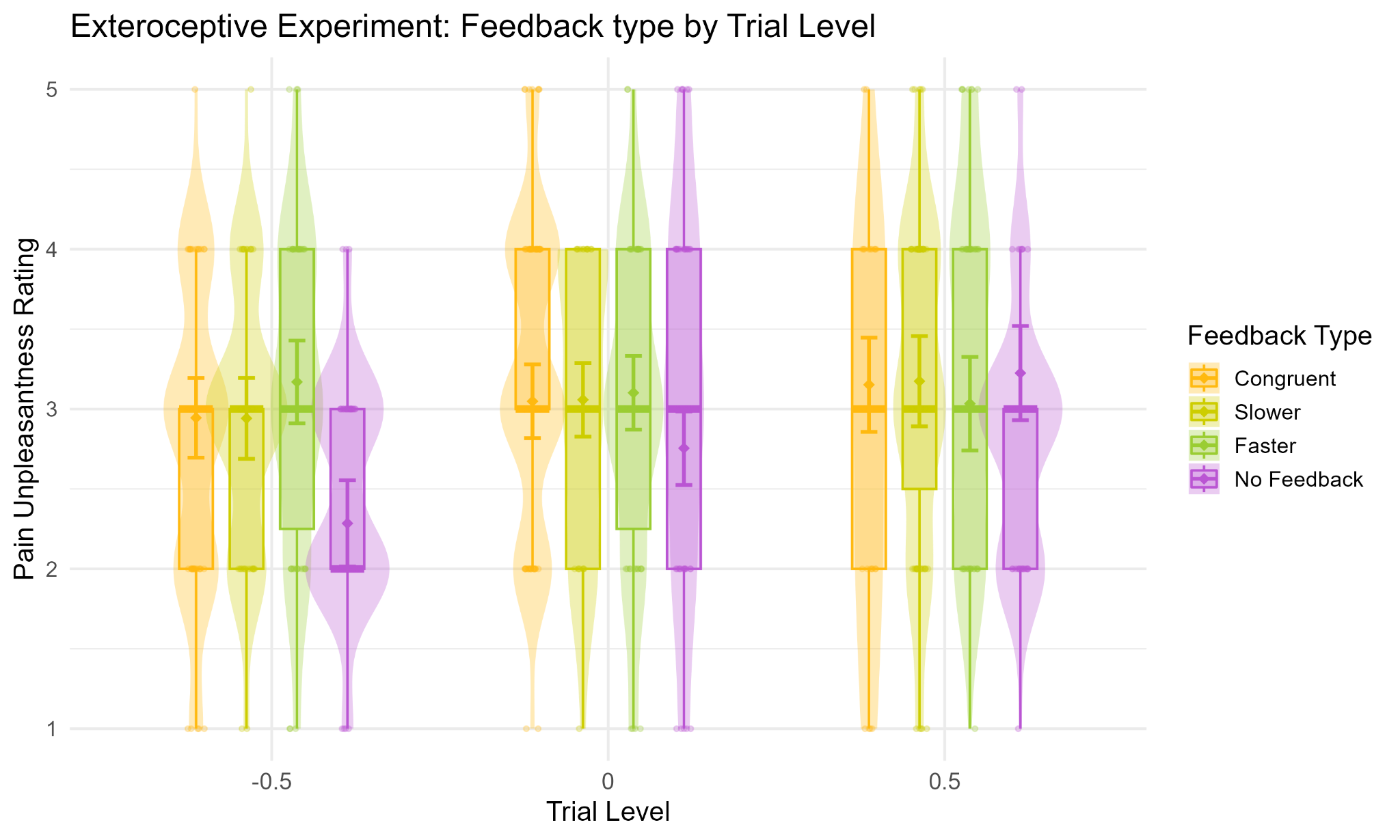

*Supplementary Figure 8.* Exteroceptive experiment: Model-predicted pain unplesantness ratings as a function of standardized trial progression (Trial) across feedback conditions (Feedback × Trial, *F*(3, 568.85) = 11.47, *p* < .0001). The Trial variable is modeled as a continuous, mean-centered predictor reflecting the temporal progression of the task (from -0.5 = early trials to +0.5 = late trials). Violin plots display the distribution and density of predicted ratings at early, middle, and late stages of the session

##### 3.3 Numeric Pain Scale of intensity ratings

To examine variations in pain intensity ratings within the exteroceptive experiment, we applied general linear mixed-effects modeling (GLMM), the model was fitted using R (lme4 package; Bates et al., 2015; Bates et al., 2015) and was specified as follows:

NUMERIC PAIN SCALE OF INTENSITY RATINGS ~ Feedback × StimInt × Trial + (StimInt + Trial | Subject)

where Feedback, StimInt, and Trial were included as fixed effects. A random slope for StimInt and Trial was included at the subject level (StimInt + Trial | Subject) to account for individual differences in pain perception across stimulus intensities and over time. Model comparison confirmed that including these random slopes significantly improved model fit (χ² = 56.623, p < < 0.001).

###### 3.3.1 Results

###### 3.3.1.1 Effects of interest

The results of the Type III ANOVA with Satterthwaite’s approximation revealed a significant main effect of **Feedback**, F(3, 565.43) = 34.72, p < .001, indicating that pain intensity ratings (Numeric pain scale of intensity ratings) varied across feedback conditions. Compared to the reference condition (i.e., Congruent feedback), Numeric pain scale of intensity ratings were significantly lower in the No Feedback condition (b = –8.47, SE = 1.17, t(565.85) = –7.23, p < .0001), while the Faster feedback condition showed a marginally significant increase in ratings (b = 2.23, SE = 1.17, t(564.96) = 1.91, p = .0562). The Slower feedback condition did not differ significantly from the Congruent condition (b = 0.84, SE = 1.17, t(565.11) = 0.72, p = .4727).

Post hoc comparisons further clarified these effects. Numeric pain scales of intensity ratings were significantly lower in the No Feedback condition compared to all other feedback conditions (all ps < .0001).

Specifically, intensity ratings were significantly higher in the Faster condition compared to No Feedback (estimate = 10.545, SE = 1.16, *t*(566) = 9.124, *p* < .0001), as well as in the Slower condition compared to No Feedback (estimate = 9.181, SE = 1.16, *t*(566) = 7.933, *p* < .0001,), and in the Congruent compared to the No feedback (estimate = 8.383, SE = 1.17, *t*(566) = 7.150, *p* < .0001). *No significant differences emerged between active feedback conditions (all ps > .10)*, suggesting that the changes in pain ratings were driven specifically by the absence of feedback, rather than differences among feedback types.

###### 3.3.1.2 Additional unpredicted effects

A significant main effect of **Trial** also emerged, F(1, 29.23) = 8.86, p = .0058, suggesting that pain intensity ratings changed throughout the experimental session. The fixed-effect estimate of the Trial coefficient indicated a significant positive trend (b = 7.86, SE = 3.64, t(83.06) = 2.16, p = .0337), consistent with a general increase in pain ratings over time.

Critically, a strong main effect of **Stimulus Intensity** was observed, F(1, 28.31) = 187.17, p < .001, indicating that pain intensity ratings increased as a function of the objective strength of nociceptive stimulation. The fixed-effect estimate confirmed a robust positive association (b = 37.90, SE = 3.15, t(60.25) = 12.05, p < .001), aligning with theoretical expectations and validating the experimental manipulation of stimulus intensity.

The **Feedback × Stimulus Intensity** interaction was significant, F(3, 570.65) = 3.00, p = .0301, indicating that the effect of stimulus intensity on pain ratings varied across feedback conditions. To further unpack this interaction, estimated marginal means (EMMs) of Numeric pain scale of intensity ratings were computed at three levels of stimulus intensity: low (–0.5), medium (0), and high (+0.5).

At *low stimulus intensity* (–0.5), the Slower condition elicited the highest pain ratings (M = 44.7, SE = 3.38), followed by Faster (M = 42.2, SE = 3.38), Congruent (M = 40.4, SE = 3.38), and No Feedback (M = 33.8, SE = 3.37). Post hoc comparisons revealed that the Slower condition yielded significantly higher ratings than Congruent (estimate = –4.24, SE = 1.87, t(570) = –2.27, p = .0351). Moreover,all active feedback conditions (Slower, Faster, Congruent) elicited significantly higher ratings than No Feedback (all ps < .001). No other contrasts reached significance.

At *medium intensity* (0), ratings followed a similar pattern: Faster produced the highest ratings (M = 61.6, SE = 2.74), followed by Slower (M = 60.2, SE = 2.74), Congruent (M = 59.4, SE = 2.74), and No Feedback (M = 51.1, SE = 2.74). While differences among active feedback conditions were not statistically significant (all ps > .09), each of them led to significantly higher ratings than the No Feedback condition (all ps < .0001).

At *high intensity* (+0.5), the highest ratings were reported in the Faster condition (M = 81.0, SE = 2.91), followed by Congruent (M = 78.4, SE = 2.93), Slower (M = 75.8, SE = 2.91), and No Feedback (M = 68.3, SE = 2.91). The No Feedback condition was significantly lower than all other feedback conditions (all ps < .0002), and pain ratings were significantly higher in the Faster compared to the Slower condition (estimate = –5.14, SE = 1.82, t(568) = –2.83, p = .0072). No significant difference was observed between Congruent and Faster (p = .1682).

Numeric pain scale of intensity ratings increased with stimulus intensity in all feedback conditions. However, the rate of increase (slope) differed across feedback types. The steepest increase was observed in the Faster condition (b = 38.8, SE = 3.14, t(59.4) = 12.36, p < .0001), followed closely by Congruent (b = 38.0, SE = 3.15, t(60.5) = 12.05, p < .0001), No Feedback (b = 34.5, SE = 3.12, t(58.2) = 11.07, p < .0001), and Slower (b = 31.1, SE = 3.13, t(58.7) = 9.96, p < .0001). Pairwise comparisons between slopes revealed that the increase in pain ratings was significantly steeper in the Faster condition compared to Slower (estimate = –7.61, SE = 2.86, t(571) = –2.66, p = .0484). All other comparisons between slopes were not statistically significant (all ps > .05).

The interaction between **Feedback × Trial** was significant, F(3, 569.80) = 12.13, p < .001, indicating that the trajectory of Numeric pain scale of intensity ratings ratings across trials varied depending on the type of feedback received.

To further explore this interaction, the estimated slopes of Trial were computed for each feedback condition. Pain intensity ratings increased significantly over time in the Congruent feedback condition (b = 7.91, SE = 3.65, 95% CI [0.66, 15.16], t(84.5) = 2.17, p = .0329) and even more markedly in the No Feedback condition (b = 23.53, SE = 4.13, 95% CI [15.36, 31.71], t(132.9) = 5.69, p < .0001). The Slower and Faster conditions did not show significant change over time (ps > .12), although a numerically positive (b = 5.39, SE = 3.45) and negative (b = -3.52, SE = 3.82) trend was observed for the Slower and Faster, respectively.

Post hoc pairwise comparisons of slopes (FDR-corrected) confirmed that the rate of increase in pain intensity was significantly higher in the No Feedback condition compared to Congruent (estimate = –15.63, SE = 4.38, t(571) = –3.57, p = .0008), Slower (estimate = –18.14, SE = 4.23, t(571) = –4.29, p = .0001), and Faster (estimate = –27.05, SE = 4.54, t(570) = –5.96, p < .0001). Additionally, Congruent feedback showed a significantly steeper increase over trials compared to Faster (estimate = 11.43, SE = 4.09, t(570) = 2.79, p = .0081), and Slower was significantly steeper than Faster (estimate = 8.91, SE = 3.92, t(568) = 2.27, p = .0280). No other slope comparisons reached significance (all ps > .5).

To examine how the effect of feedback on pain intensity evolved over time, estimated marginal means were computed at three levels of the standardized Trial variable (–0.5 = early, 0 = middle, +0.5 = late).

At *early trials* (Trial = –0.5), pain ratings were highest in the Faster condition (M = 63.5, SE = 3.25), followed by Slower (M = 57.6, SE = 3.19), Congruent (M = 55.6, SE = 3.12), and No Feedback (M = 39.3, SE = 3.40). Ratings in the No Feedback condition were significantly lower than all others (all ps < .0001), and Faster was significantly higher than both Congruent (estimate = –7.95, SE = 2.26, t(568) = –3.52, p = .0007) and Slower (estimate = –5.88, SE = 2.35, t(567) = –2.50, p = .0152).

At *mid trials* (Trial = 0), the pattern was similar: Faster again elicited the highest ratings (M = 61.7, SE = 2.73), followed by Slower (M = 60.3, SE = 2.73), Congruent (M = 59.5, SE = 2.74), and No Feedback (M = 51.0, SE = 2.73). All active feedback conditions resulted in significantly higher pain ratings than No Feedback (all ps < .0001), though differences among the active feedback types were not statistically significant (all ps > .08).

By *late trials* (Trial = 0.5), feedback-related differences had substantially diminished. Pain ratings were comparable across conditions : Congruent (M = 63.5, SE = 3.45), Slower (M = 63.0, SE = 3.28), No Feedback (M = 62.8, SE = 3.45), and Faster (M = 60.0, SE = 3.41), with no significant pairwise differences (all ps > .50). This indicates that while feedback had a strong modulatory effect on pain early in the session, this influence tended to fade as the session progressed.

Additionally, the effect of trial on pain ratings varied as a function of the actual intensity of the nociceptive stimulation, as indicated by a significant **Stimulus Intensity × Trial** interaction, F(1, 576.56) = 14.64, p < .001. To further investigate this interaction, estimated marginal trends of Numeric pain scale of intensity ratings across trials were computed at three levels of stimulus intensity: low (–0.5), medium (0), and high (+0.5). Pain ratings increased over trials at all intensity levels, but the rate of increase became progressively steeper with higher stimulus intensity.

At low intensity, the slope was small and not statistically significant (b = 1.44, SE = 3.20, 95% CI [–4.99, 7.87], t(51.4) = 0.45, p = .6551). A significant positive trend was observed at medium intensity (b = 8.28, SE = 2.78, 95% CI [2.60, 13.96], t(29.6) = 2.98, p = .0058), which became even more pronounced at high intensity (b = 15.11, SE = 3.41, 95% CI [8.31, 21.92], t(65.0) = 4.44, p < .0001).

Pairwise comparisons between slopes (FDR-corrected) confirmed that the increase in Numeric pain scale of intensity ratings over trials was significantly steeper at higher levels of stimulus intensity. Specifically, the slope at low intensity was significantly lower than both medium (estimate = –6.84, SE = 1.79, t(577) = –3.82, p = .0001) and high intensity (estimate = –13.67, SE = 3.58, t(577) = –3.82, p = .0001). Moreover, the slope at medium intensity was also significantly lower than at high intensity (estimate = –6.84, SE = 1.79, t(577) = –3.82, p = .0001). These findings suggest that the cumulative impact of repeated nociceptive stimulation on pain perception becomes progressively stronger with increasing stimulation intensity.

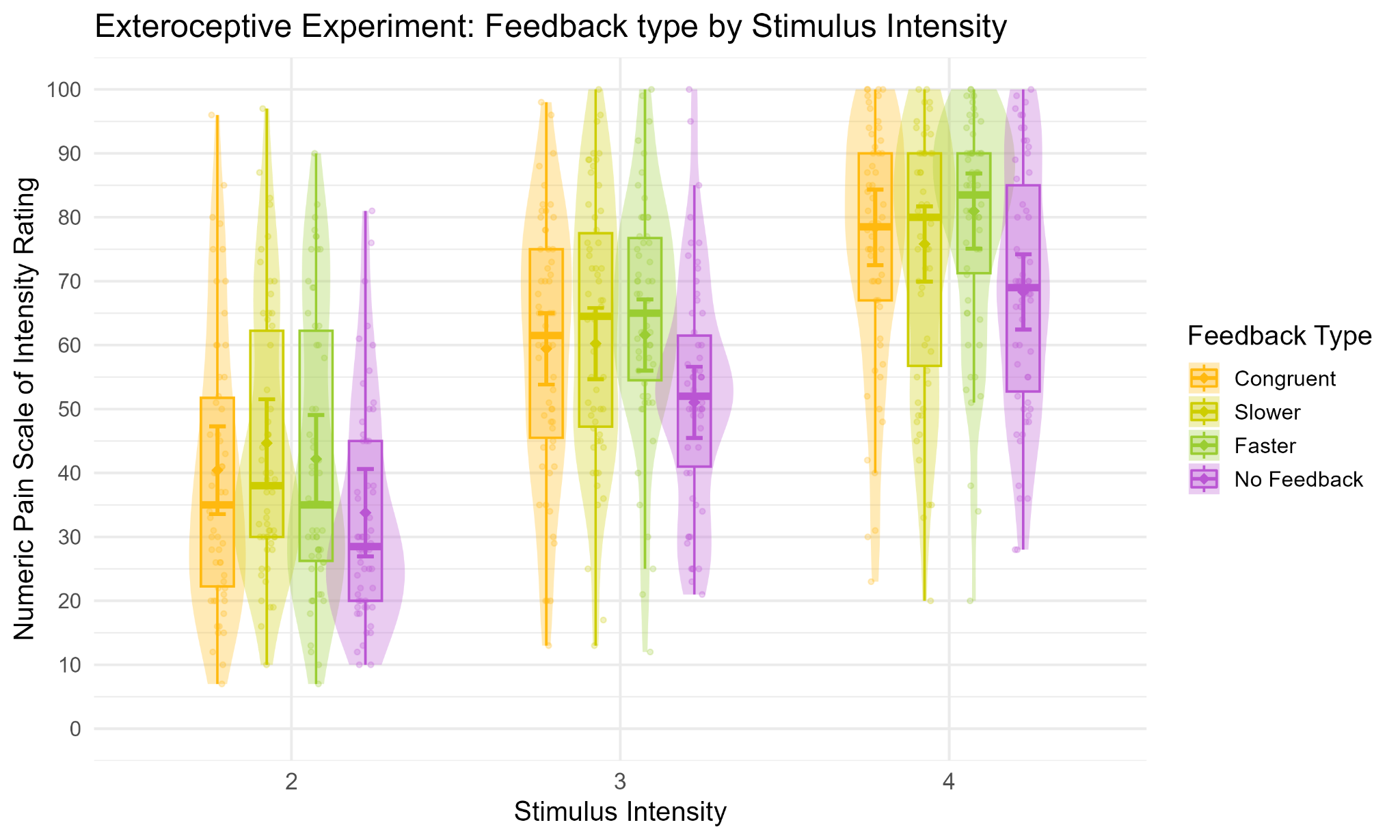

*Supplementary Figure 9.* Exteroceptive experiment: Model-predicted Numeric Pain Scale of intensity ratings (NPS) for each stimulation intensity (StimInt 2–4), across feedback conditions (Feedback × Stimulus Intensity, F(3, 570.65) = 3.00, p = .0301). Violin plots and overlaid boxplots depict the distribution (violin) and interquartile range with median (box) of the observed data for each condition. Boxes are centred on the x-axis categories because they summarise the data within each stimulation intensity and feedback condition. Large coloured dots and error bars show the model-predicted estimated marginal means (EMMs) ± standard errors for each condition.

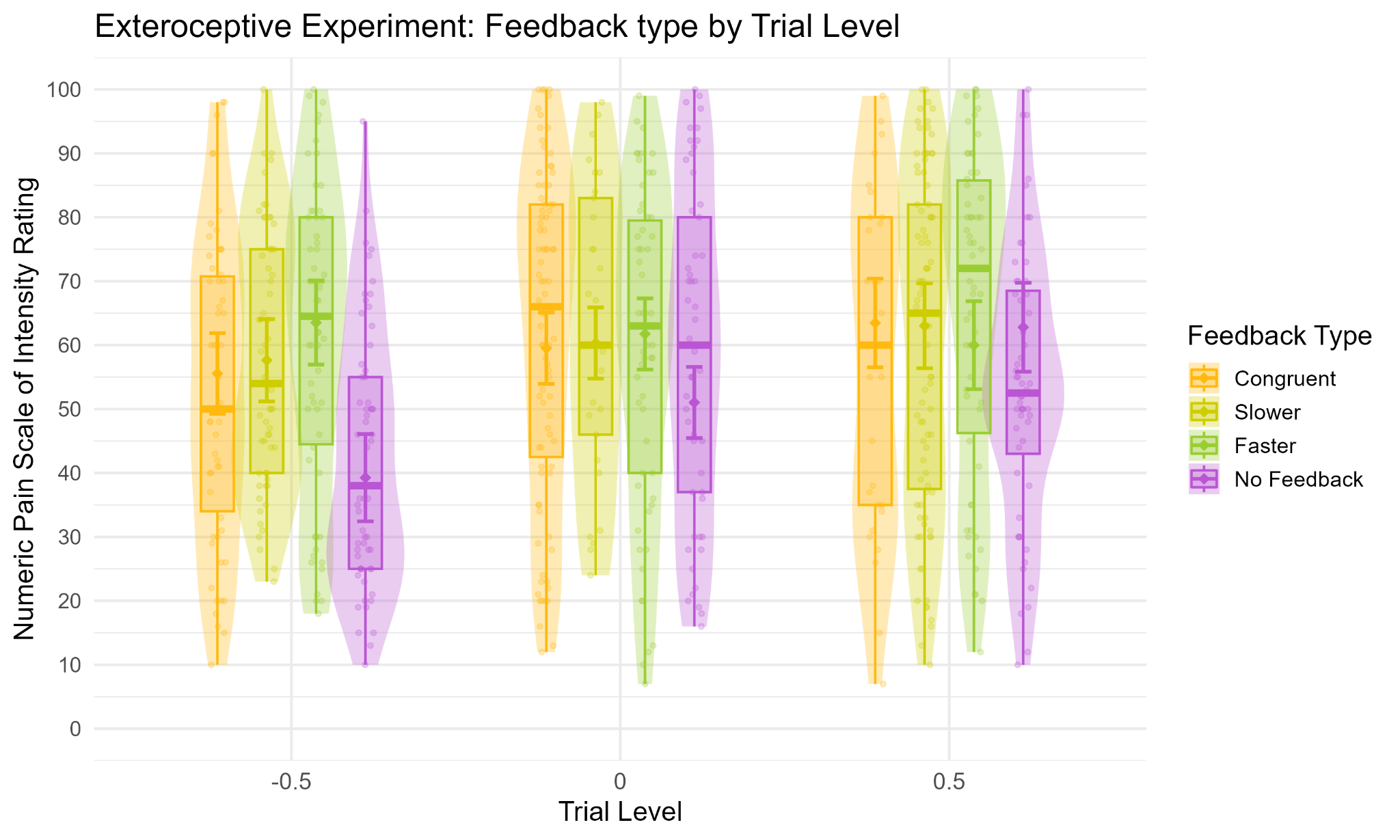

*Supplementary Figure 10.* Exteroceptive experiment: Model-predicted Numeric Pain Scale of intensity ratings (NPS) as a function of standardized trial progression (Trial) across feedback conditions (Feedback × Trial, *F*(3, 569.80) = 12.13, *p* < .001). The Trial variable is modeled as a continuous, mean-centered predictor reflecting the temporal progression of the task (from -0.5 = early trials to +0.5 = late trials). Violin plots display the distribution and density of predicted ratings at early, middle, and late stages of the session

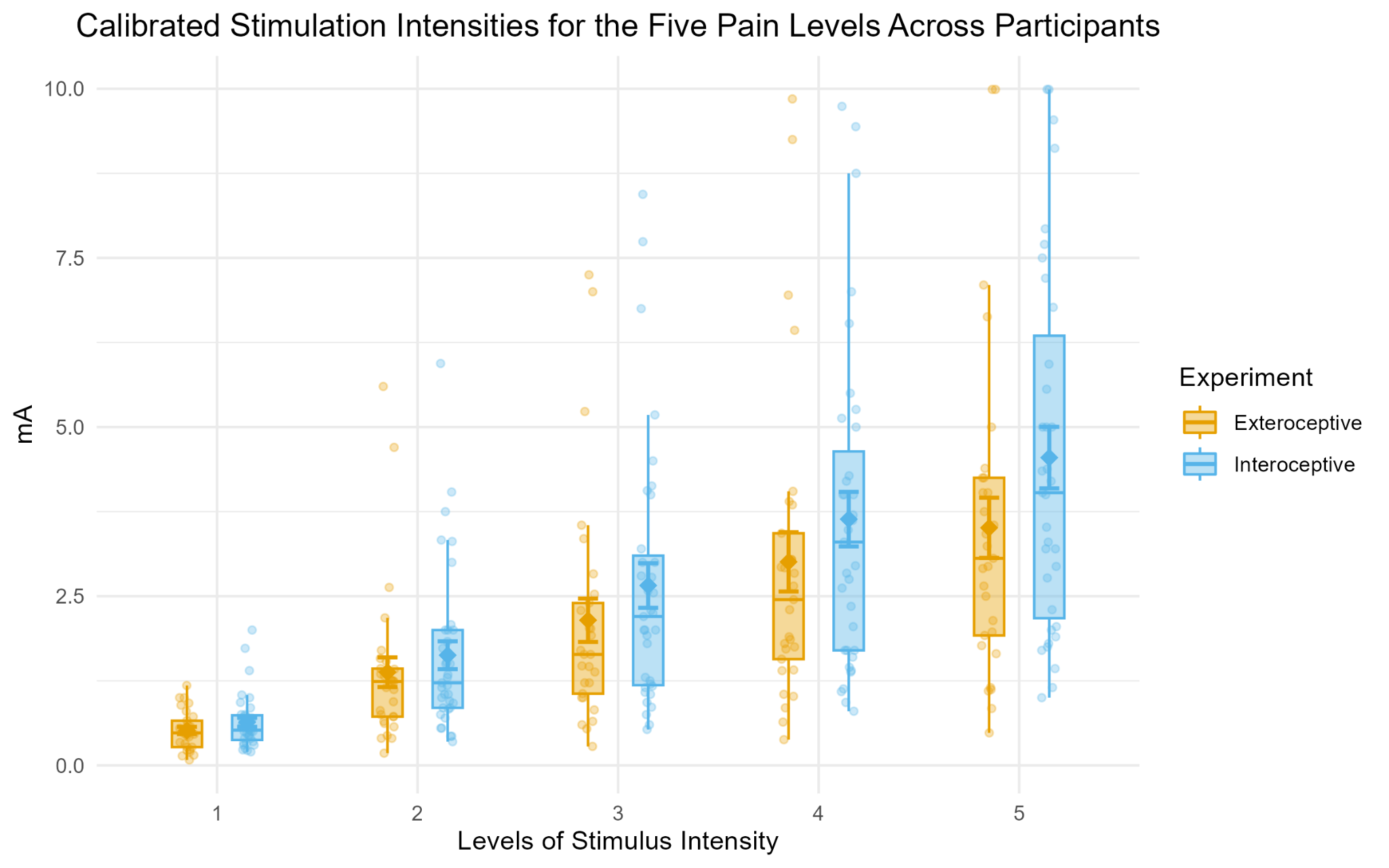

*Supplementary Figure 11.* Individually calibrated stimulation intensities (in mA) associated with five subjectively defined pain levels (NPS 10, 30, 50, 70, and 90), shown separately for the Exteroceptive and Interoceptive experiments. Each violin plot includes the distribution of individual values, boxplots with interquartile range and median, and the mean ± standard error.
